# White matter connections of human ventral temporal cortex are organized by cytoarchitecture, eccentricity, and category-selectivity from birth

**DOI:** 10.1101/2024.07.29.605705

**Authors:** Emily Kubota, Xiaoqian Yan, Sarah Tung, Bella Fascendini, Christina Tyagi, Sophie Duhameau, Danya Ortiz, Mareike Grotheer, Vaidehi S. Natu, Boris Keil, Kalanit Grill-Spector

## Abstract

Category-selective regions in ventral temporal cortex (VTC) have a consistent anatomical organization, which is hypothesized to be scaffolded by white matter connections. However, it is unknown how white matter connections are organized from birth. Here, we scanned newborn to 6-month-old infants and adults to determine the organization of the white matter connections of VTC. We find that white matter connections are organized by cytoarchitecture, eccentricity, and category from birth. Connectivity profiles of functional regions in the same cytoarchitectonic area are similar from birth and develop in parallel, with decreases in endpoint connectivity to lateral occipital, and parietal, and somatosensory cortex, and increases to lateral prefrontal cortex. Additionally, connections between VTC and early visual cortex are organized topographically by eccentricity bands and predict eccentricity biases in VTC. These data show that there are both innate organizing principles of white matter connections of VTC, and the capacity for white matter connections to change over development.

## Introduction

Human ventral temporal cortex (VTC) contains regions that are selective for categories that are important for our everyday lives such as faces^1^, bodies^2^, words^3^, and places^4^. A key debate in cognitive neuroscience is what structural and functional factors contribute to the consistent functional organization of VTC^5–10^. One central theory is that innate white matter connections between VTC and other parts of the brain may lead to the emergence of category-selective regions in consistent anatomical locations relative to cortical folds^11–16^. Indeed, recent research in adults has revealed that the white matter connections of VTC are highly regular with respect to cortical folds^17–19^, cytoarchitecture^20^, eccentricity^21^, and category-selectivity^11,12,22^. However, it is unknown which of these organizing principles of white matter connections of VTC are already present in infancy and if white matter connections remain stable from infancy to adulthood. Here, we address these gaps in knowledge by using anatomical and diffusion MRI in infants and adults to elucidate the organization principles of white matter connections of VTC from birth to adulthood.

We first consider multiple hypotheses of how white matter connections at birth may constrain the development of VTC function. The *category hypothesis* suggests that category-selective regions in VTC have category-specific connections forming specialized networks that enable processing of category-specific information^15,23,24^. While some studies have found that face-selective regions in VTC only emerge with visual experience^25,26^, and develop during the first year of life^27^, the existence of category-selective regions in the congenitally blind^28,29^, has led to the hypothesis that innate white matter connections might scaffold the development of category-selective regions in VTC. This theory is supported by studies showing that, in adults, white matter connectivity profiles predict the location of face, place, and word-selective regions in VTC^11,22,30^ and in children, white matter connectivity profiles at age five predict where word-selective regions will emerge at age eight^13^. Further, functional connectivity of resting state data in infants show category-specific patterns^31,32^, suggesting that category-specific networks may be present from birth. The category hypothesis thus predicts that white matter connections of VTC will be organized by category from birth.

The *cytoarchitecture hypothesis* suggests that white matter connections are linked to cytoarchitectonic areas – areas defined by their distribution of cell density across cortical layers^33–35^. VTC contains four cytoarchitectonic areas: FG1-FG4^36,37^, which have different neural hardware that is thought to support different functional computations. Notably, in both children and adults, regions with different category-selectivity within the same cytoarchitectonic area, e.g., face-selective and word-selective regions located in FG4, have similar white matter connectivity profiles^20^. But regions with the same category-selectivity located in different cytoarchitectonic regions, e.g., face-selective regions located in FG2 and FG4, respectively, have different white matter connectivity profiles^20^. The cytoarchitectonic hypothesis therefore predicts that white matter connections of VTC will be organized by cytoarchitecture from birth.

The *eccentricity-bias hypothesis* suggests that eccentricity biases in VTC are due to innate white matter connections with early visual cortex (EVC, union of V1-V2-V3), where faces and word-selective regions have more connections with foveal EVC and place-selective regions have more connections with peripheral EVC. This theory is supported by findings that in both children and adults, face and word-selective regions in VTC have a foveal bias^5,7,8,38^, whereas place-selective regions in VTC have a peripheral bias^5,7,8,38^. Additionally, patterns of functional connectivity between face- and place-selective regions in VTC and EVC in infant humans, neonate monkeys, and congenitally blind adults follow eccentricity bands^39–43^. That is, face-selective regions have higher functional connectivity to central eccentricity bands in EVC and place-selective regions have higher functional connectivity to peripheral eccentricity bands in EVC. Finally, in adults, white matter connections between VTC and EVC also correspond to eccentricity bands, with face-selective regions having more connections to central EVC eccentricities and place-selective regions having more connections to peripheral ones^21^. The eccentricity-bias hypothesis thus predicts that the subset of white matter connections between VTC to EVC will have a non-uniform distribution, where regions in VTC that have foveal bias in adults (faces and words) will have more connections with central bands in EVC from birth, and regions that have a peripheral bias (places) will have more connections with peripheral bands in EVC from birth. Crucially, when experiments only include the categories of faces and places, it is not possible to distinguish between the category and eccentricity hypotheses because there is one-one-mapping between category and eccentricity bias. In the present study, we include multiple categories that have a foveal bias (faces and words), which will allow us to distinguish the category from the eccentricity hypotheses.

In addition to the unknown organizational principles of white matter connections in infancy, it is also unknown if white matter connectivity profiles are stable or change over development. Accumulating evidence reveals that the large fascicles (axonal bundles that travel in parallel connecting distant parts of the brain) are present at birth^44–48^. Nonetheless, some fascicles continue to develop after birth. For example, the arcuate fasciculus (AF), which connects the VTC with lateral prefrontal cortex is not fully developed in infancy: it has a smaller cross section in children compared to adults^49^ and, and in infants, it reaches premotor cortex but not lateral prefrontal cortex as it does in adults^50^. Additionally, mature white matter connections in adulthood may depend on visual experience as visual deprivation during infancy and childhood leads to degradation of white matter tracts of the visual system^51–55^. However, it is unknown which aspects of VTC white matter connectivity profiles are innate and which aspects may develop from infancy to adulthood.

To address these gaps in knowledge, we obtained diffusion magnetic resonance imaging (dMRI) and anatomical MRI in infants and adults and evaluated the organization and development of white matter connections of category-selective regions of VTC.

## Results

We collected MRI data from 43 newborn to 6-month-old infants (26 longitudinally) during natural sleep using 3T MRI over 75 sessions, as well as from 21 adults (28.21±5.51 years), obtaining whole brain anatomical MRI and multishell dMRI data in each participant. We implemented several quality assurance measures to ensure high quality data (Methods: Quality Assurance). Two infant sessions were excluded because of missing diffusion data, and six infant sessions were excluded after quality assurance. We report data from 67 infant sessions (42 infants (16 female, 26 male); 21 longitudinal infants; timepoints in Supplementary Figure 1) and 21 adult sessions (17 female, 4 male) with no significant differences in data quality (see Methods). In total, we report data from 88 sessions: 23 sessions from newborns (mean age±standard deviation: 28.6±10.2 days), 23 sessions from 3-month-olds (106.9±19.3 days), 21 sessions from 6-month-olds: (189.0±15.8 days), and 21 sessions from adults (28.21±5.51 years). Anatomical MRIs were used to segment the brain to gray and white matter, define the gray-white matter boundary to seed the tractography, and create cortical surface reconstructions. dMRI data was used to derive the whole brain white matter connectome of each individual and session.

To determine how white matter connections are organized in infants and adults and how they may constrain the development of regions in VTC, we project adult functional regions of interest (fROIs) to the native space of each individual and determine the white matter connections of each fROI. To do so, we use cortex based alignment^56^ to project maximal probability maps (MPMs) of six adult category-selective regions from independent data^20^ into each individual brain and session. The fROIs are: mFus-faces (face-selective region located near anterior end of the mid fusiform sulcus), pFus-faces (posterior fusiform face-selective region), OTS-bodies (body-selective region located on the occipital temporal sulcus), CoS-places (place-selective region located on the collateral sulcus), mOTS-words (word-selective region located on the occipital temporal sulcus), and pOTS-words (posterior OTS word-selective region). MPMs of fROIs selective for faces, bodies, and places (mFus-faces, pFus-faces, OTS-bodies and CoS-places) are bilateral, and MPM fROIs selective for words (mOTS-words and pOTS-words) are in the left hemisphere only (Fig 1A,B).

**Figure 1.**
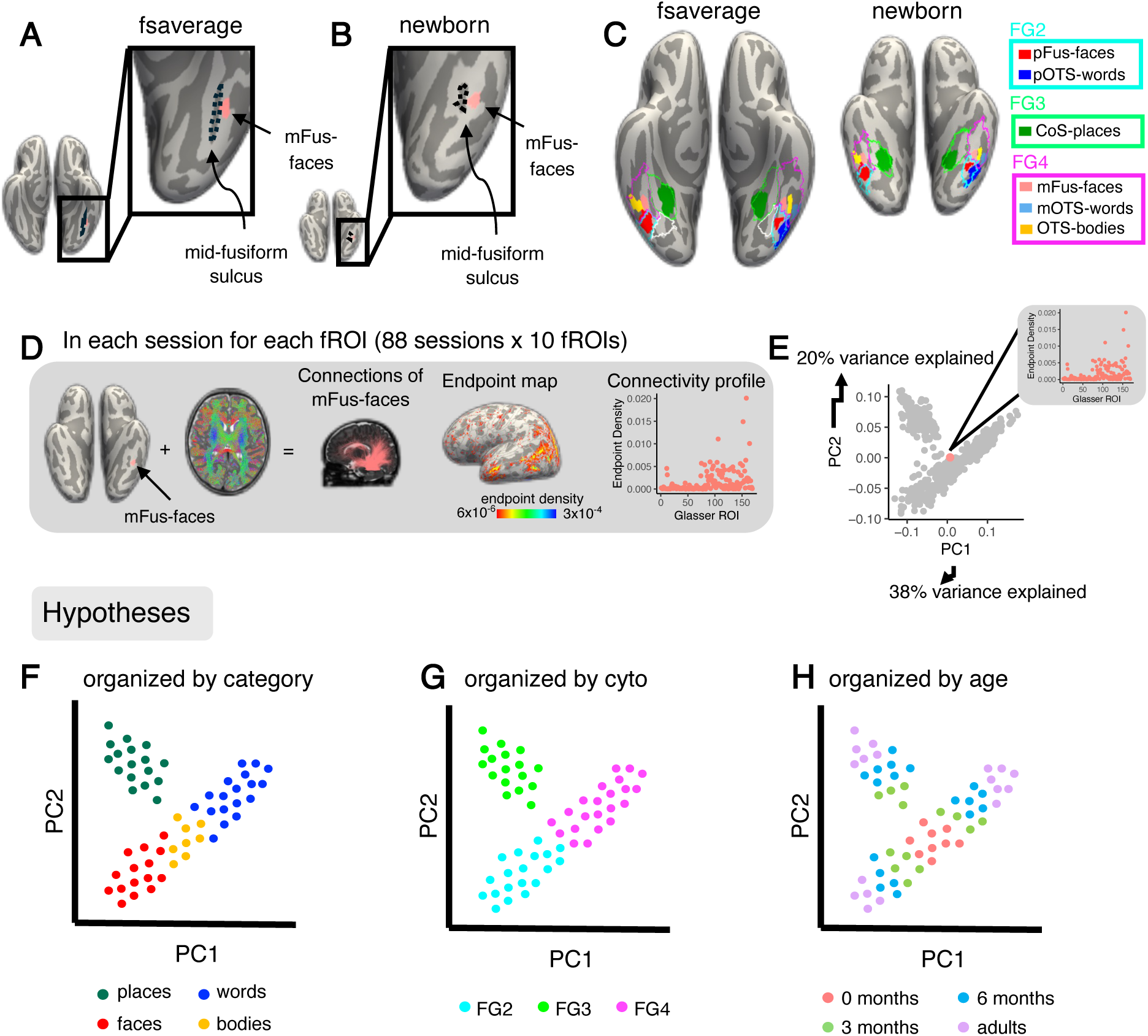
Functional atlas and analysis pipeline. A) mFus-faces aligns to the mid-fusiform sulcus in the fsaverage brain space (left) and B) the newborn brain space (right). C) *Left:* Functional region of interest (fROI) atlas from 28 adults in^20^ and Atlas of cytoarchitectonic areas from^59^ projected to the fsaverage brain; *Right:* the same atlases projected into an example newborn’s brain (17 days old). Solid colors represent fROIs. Salmon *pink:* mFus-faces (mid fusiform face-selective region); *Light blue:* mOTS-words (mid occipital temporal sulcus word-selective region); *Yellow:* OTS-bodies (occipital temporal sulcus body and limb-selective region); *Red:* pFus-faces (posterior fusiform face-selective region); *Blue: p*OTS-words (posterior occipital temporal sulcus word-selective region). Green: CoS-places (collateral sulcus place-selective region). Outlines represent cytoarchitectonic areas^59^. *Cyan:* FG2; *Green:* FG3; *Magenta:* FG4. Both fROIs and cytoarchitectonic areas align to the expected anatomical landmarks in the newborn. All individuals in Supplementary Figs 2-9. D) Pipeline to define white matter connectivity profiles for each fROI. *Left:* example connections of mFus-faces in an individual newborn; *Middle*: endpoint map on the cortical surface; *Right:* quantification of endpoint density within each Glasser ROI. E) Principal Components Analysis plot across the first two principal components: explaining 38% and 20% of the variance, respectively. Each dot represents a single connectivity profile in a single subject; colored dot represents the connectivity profile shown in (D). (F-H) Schematics illustrating what the data may look like in PC space according to the predictions of three hypotheses. Each dot represents a connectivity profile for a given fROI and session. (F) Category hypothesis. *Green:* place-selective, *blue:* word-selective, *red:* face-selective, *yellow:* body-selective. G) Cytoarchitectonic hypothesis. *Cyan:* FG2, *green:* FG3, *magenta:* FG4. H) Age hypothesis. *Salmon pink:* 0-months, *green:* 3 months, *blue:* 6 months, *purple:* adults. *FG*: fusiform gyrus.

To verify that fROIs are not proportionally different in size across age groups, which could affect the resulting white matter connectivity profiles, we measured the surface area of fROIs relative to the surface area of each hemisphere in all infant and adult sessions. We tested if the relative fROI size changes over development (as development plateaus, we use a logarithmic model: fROI size/surface area ∼ log10(age in days)). We find no relationship between the surface area of fROIs relative to hemisphere surface area and age (t(878) = 8.77×10^−6^, p = 0.88, adjusted R^2^ = −0.001, 95% CI = [−0.0001,0.0001]).

To ensure that fROIs and cytoarchitectonic areas align to the expected anatomical landmarks in each infant and adult session, we visually inspected each session to ensure that there is a consistent relationship between projected ROIs and anatomical landmarks. First, we checked whether the collateral sulcus (CoS) and mid fusiform sulcus (MFS) are present in both hemispheres. We found that the anatomical landmarks were present in nearly each infant (CoS: 67/67, MFS: 66/67) and adult (CoS: 21/21, MFS: 19/21) session. Next, we tested whether CoS-places aligns to the intersection of the anterior lingual sulcus and CoS^6,9^, mFus-faces aligns to the anterior tip of the MFS^57^, and whether the boundaries between FG1 and FG2 and between FG3 and FG4 are aligned to the MFS, as in prior studies^20,58^. We find that both fROIs and cytoarchitectonic areas are aligned to the expected anatomical landmarks in nearly each infant (fROIs; LH: 66/67, RH: 65/67, cytoarchitectonic areas; LH: 67/67, RH: 62/67) and adult (fROIs: LH: 21/21, RH: 21/21, cytoarchitectonic areas; LH: 20/21, RH: 21/21) (Supplementary Figs 2-9). Therefore, in both infants and adults there is a consistent relationship between MPMs of category-selective regions and cytoarchitectonic areas^20,58^, where pOTS-words and pFus-faces are within FG2, CoS-places is within FG3, and mFus-faces, OTS-bodies, and mOTS-words are within FG4 (Fig 1C).

We use a data-driven approach to determine the organizing principles of white matter connections of VTC. As the category and cytoarchitecture hypotheses make predictions about connectivity across the whole brain, whereas the eccentricity-bias hypothesis makes predictions only about connectivity with EVC, we first test if whole brain connectivity profiles are organized by category-selectivity or cytoarchitecture. In addition, we test if whole brain connectivity profiles are organized by age to determine if connectivity profiles are stable or changing over development. We separately examine the eccentricity-bias hypothesis in the next section.

We derived each fROI’s white matter connections by intersecting each fROI with the whole brain white matter connectome derived from dMRI in each session (Fig 1D). We quantified the white matter connectivity profile by measuring the endpoint density of each fROI’s connections on the whole brain as the proportion of connections ending in each of 169 Glasser Atlas ROIs^60^ (180 Glasser ROIs excluding 11 VTC ROIs, Fig 1D). This resulted in 880 unique connectivity profiles (10 fROIs x 88 sessions across infants and adults). Then we used principal component analysis (PCA) to reduce the dimensionality of the connectivity profiles. The first 10 principal components (PCs) explain 98% of the variance in the data, with the first and second components explaining 38% and 20%, respectively.

To visualize how connectivity profiles cluster, we plot the connectivity profiles of all fROIs and sessions in PC space (Fig 1E). The plot consists of 880 dots including data from all infants and adults, where each dot represents a single connectivity profile of an fROI from one session. Note that each white matter connectivity profile has several non-mutually exclusive features: the category-selectivity of the fROI, the cytoarchitectonic area in which the fROI is located, and the age of the participant. Therefore, we can label each connectivity profile by each of these features (category, cytoarchitecture, age group) and test if connectivity profiles cluster by one or more of these features. For example, if connectivity profiles are organized by category, then connectivity profiles of fROIs with the same category-selectivity would be nearby in PC space and separate from connectivity profiles of fROIs with a different category-selectivity (Fig 1F). However, if connectivity profiles are organized by cytoarchitecture, then connectivity profiles of fROIs in the same cytoarchitectonic area would be nearby in PC space (Fig 1G). Further, if connectivity profiles change across development, then connectivity profiles of newborns may be different than other age groups and connectivity profiles of adults may be most distinct (Fig 1H).

Labeling connectivity profiles by fROIs’ preferred category reveals some clustering by category (Fig 2A). For example, connectivity profiles of face- and place-selective ROIs tend to be clustered and separate from others. However, clustering by category is imperfect, as connectivity profiles of word-selective regions split into two clusters, and there is overlap between the connectivity profiles of word- and body-selective fROIs. In contrast, labeling connectivity profiles by fROIs’ cytoarchitectonic area reveals a strikingly clear organization whereby connectivity profiles of each of the fusiform gyrus (FG) cytoarchitectonic areas FG2, FG3, FG4 form a different cluster with a distinct connectivity profile (Fig 2B). Examination of the PC loadings (Supplementary Fig 10) and the coefficients across the first 2 PCs suggests that connectivity profiles of fROIs in different cytoarchitectonic areas largely vary in their connections to early visual, lateral occipito-temporal, and anterior temporal cortex. Finally, labeling connectivity profiles by participant’s age group reveals that connectivity profiles are intermixed across age groups with no clear organization by age across the first 2 PCs (Fig 2C).

**Figure 2.**
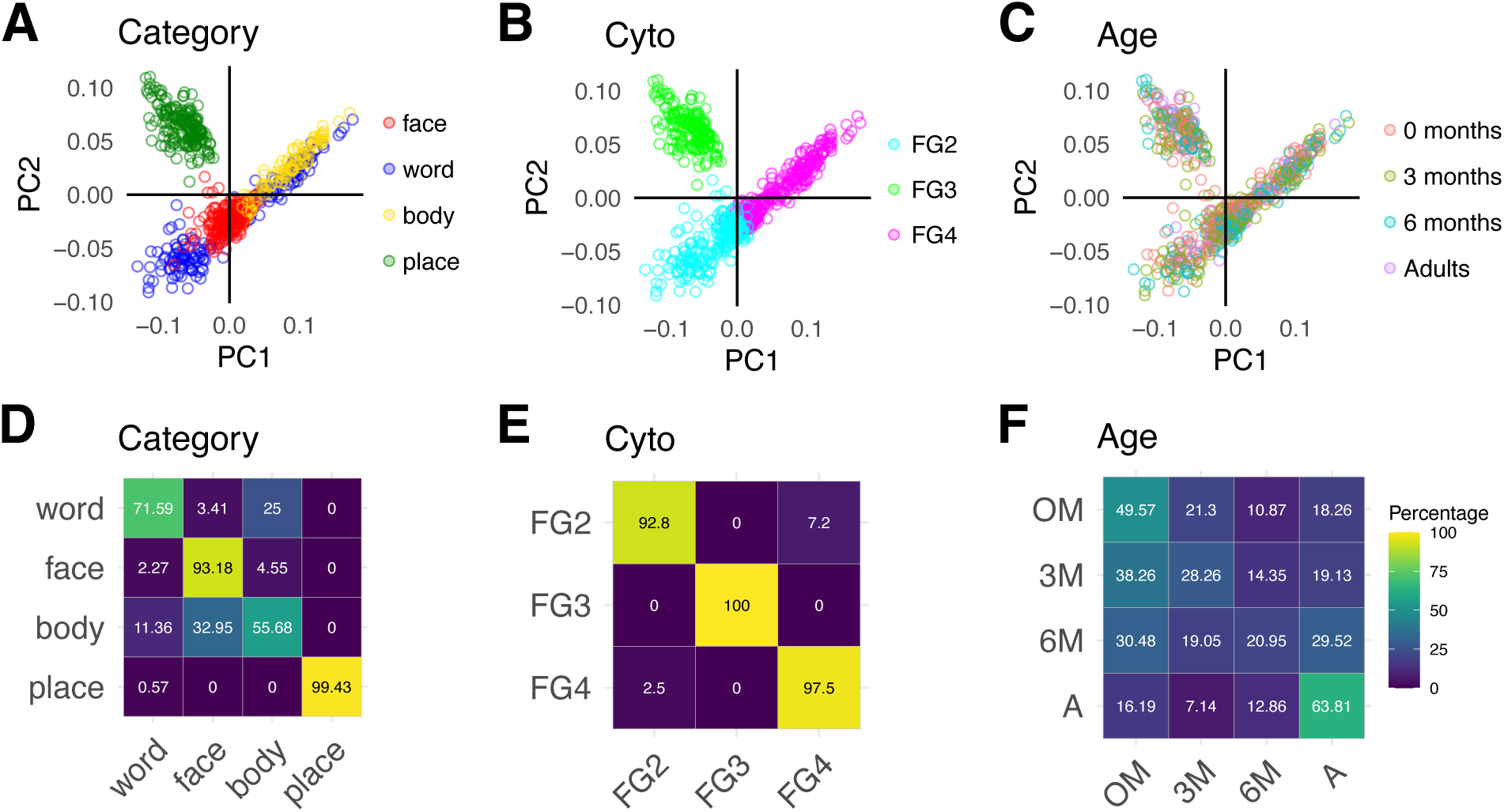
White matter connections are organized by cytoarchitecture and category from infancy. (A-C) White matter connectivity profiles of all participants (n = 88) and fROIs (n=10) projected to the first two principal components (PC1 and PC2); each dot is a connectivity profile of a single fROI and session. The same data is shown in all three panels, except that they are colored by different features. (A) Category: *Red*: faces, *green:* places, *blue*: words, *yellow:* bodies, (B) Cytoarchitectonic area: *Cyan:* FG2, *green:* FG3, *magenta:* FG4. *FG:* fusiform gyrus. (C) Age group: *Salmon:* newborn, *green:* 3 months, *teal:* 6 months, *purple*: adults. (D-F) Classification accuracy of a feature from white matter connectivity profiles, using multinomial logistic regression with leave-one-out-participant cross validation. In D-F, Rows sum to 100. Color depicts the percentage of samples classified within each bin; brighter colors indicate higher percentage of samples (see color bar). On diagonal values are percentage correct classification and off diagonal values are percentage incorrect classification (D) Confusion matrix for classification of category. (E) Confusion matrix for classification of cytoarchitectonic area. (F) Confusion matrix for classification of age group. Overall classification accuracy is found in Supplementary Figure 11.

To utilize the information across all 10 PCs, we use a leave-one-out classification approach to test if a classifier trained on white matter connectivity profiles of n-1 participants can predict the category preference, cytoarchitectonic area, or the participant’s age group from the left-out participant’s connectivity profiles. We find that category (mean accuracy±SD = 83%±10%, 95% CI = [81%, 85%], t(87) = 54.68, p < 0.001, Cohen’s d = 5.83), cytoarchitecture (mean accuracy±SD= 97%±5%, 95% CI = [95%, 98%], t(87) = 109.6, p < 0.001, Cohen’s d = 11.68), and age (mean accuracy±SD = 41%± 27%, 95% CI = [35%, 46%], *t*(87) = 5.48, *p* < 0.001, Cohen’s d = 0.58) are all classified significantly above chance (Supplementary Fig 11). Classification of cytoarchitecture is significantly higher than classification of category preference (odds ratio= 5.96, 95% CI [3.98 8.93], binomial logistic regression) and age (odds ratio = 41.51, 95% CI [28.16, 61.19], binary logistic regression). Confusion matrices reveal almost no error for cytoarchitecture classification (Fig 2E), but some category errors that arise from confusion of word-and body-selectivity as well as confusion of body and face-selectivity (Fig 2D). The confusion matrix for age classification (Fig 2F) reveals that newborns and adults are classified the best, that 3-month-olds are often confused for newborns, and that 6-month-olds are equally confused across all age groups. This suggests that there is heterogeneity in the connectivity profiles of 6-months-olds, and that around 6 months of age, the white matter connections of some infants may start to become more adult-like, while connections in others resemble 3-month-olds and even newborns.

As there are unbalanced training sets (e.g., two face-selective vs. one body-selective fROI), we performed an additional analysis where we created a training set that had an equal number of examples for each label by sampling the training set with replacement. We find similar classification results with balanced examples in the training data (Supplementary Fig 12).

To assess if the high classification performance of cytoarchitecture is due to higher anatomical proximity of fROIs within the same cytoarchitectonic area compared to those in distinct cytoarchitectonic areas, we repeated the classification analysis on connectivity profiles of equidistant disk ROIs that are either in the same (FG4) or different (FG2/FG4) cytoarchitectonic areas (Supplementary Fig13A,B). We reasoned that if anatomical proximity explains classification results, then classification accuracy would be reduced once distance is controlled. However, even when distance was held constant, cytoarchitecture is classified from the connectivity profiles with high accuracy (Supplementary Fig13C, mean accuracy±SD=95%±22%, t(87) = 31.62, p < 0.001, Cohen’s d = 3.37, 95% CI = [0.92,0.98]), suggesting that organization of connectivity profiles by cytoarchitecture is not just due to anatomical distance between fROIs.

Interestingly, both category and cytoarchitecture classification are not significantly different across age groups (comparison between newborns, 3-month-olds, and 6-month-olds compared to adults (newborns: odds ratio =1.20, 95% CI [0.77, 1.89], 3-months: odds ratio =1.01, 95% CI [0.65, 1.56], 6-months: odds ratio =1.05, 95% CI [0.67, 1.65], binomial logistic regression). Indeed, examination of the organization structure across the first 2 PCs separately for each age group reveals similar organization across age groups (Supplementary Fig 14). These analyses suggest that organizational features of white matter connections by cytoarchitecture and category persist across development.

### Are Connections Between VTC and EVC Organized Retinotopically?

To test the eccentricity-bias hypothesis we next examined the subset of connections between each fROI and early visual cortex (EVC). We reasoned that if connections between VTC and EVC are organized by eccentricity from birth, then fROIs that overlap with foveal representations in adults (pFus-faces, pOTS-faces, mFus-faces, OTS-bodies, mOTS-words) would have more white matter connections to central than peripheral eccentricities in EVC in infancy, and fROIs that overlap with peripheral representations in adults (CoS-places) would have more connections to peripheral than central eccentricities in VTC in infancy. To test these predictions, we measured the distribution of white matter endpoints between each of the VTC category-selective fROIs and three eccentricity bands in EVC: 0-5°, 5-10°, and 10-20° and compared across age groups.

We first visualized the white matter connections between each fROI and EVC, coloring the white matter connections according to the endpoint eccentricity in EVC. This visualization reveals a striking orderly arrangement of the connections from each eccentricity band to each of the fROIs in both infants and adults (Fig 3A and 3B; 3D visualization: Supplementary Videos 1-4). Specifically, in all age groups, newborns to adults, there is a lateral to medial arrangement of connections such that connections to EVC central eccentricities (red, 0-5°) are more lateral, and connections to more peripheral EVC eccentricities are more medial (green, 5-10° to blue, 10-20°; Fig 3A,B).

**Figure 3.**
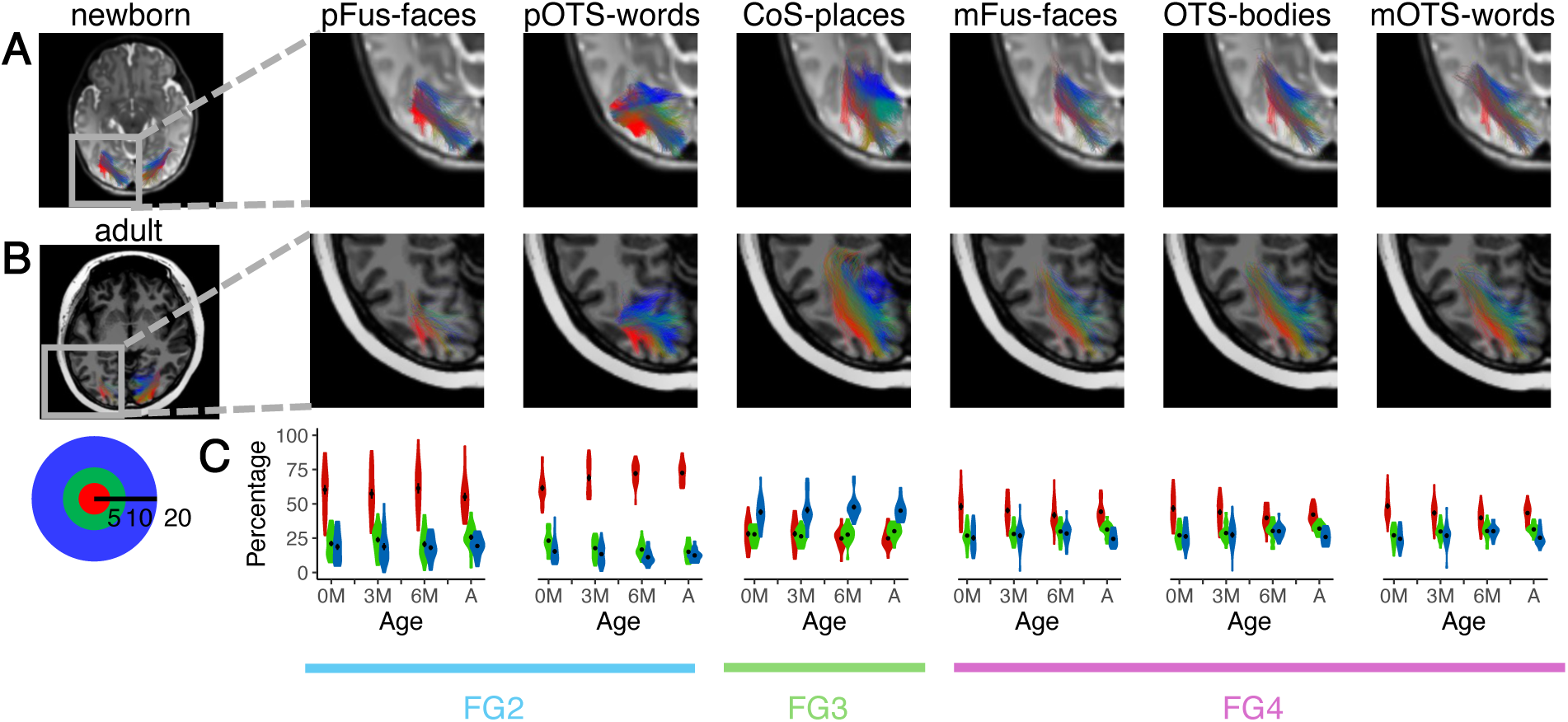
White matter connections of VTC are organized by eccentricity from birth. Connections between each fROI and early visual cortex (EVC, union of V1, V2, V3) in an example newborn (A) and an example adult (B). Here we show data for the left hemisphere as it includes all 6 fROIs; data for the right hemisphere is in Supplementary Fig 56. Connections are colored by the endpoint eccentricity band in EVC: *Red:* 0-5°, *green*: 5-10°, *blue:* 10-20°. (C) Quantification of the percentage of endpoints in each eccentricity band by age group for each fROI. Violin plots indicate distributions of the percentage of connections for each age group and eccentricity band. *Black dot and error bars*: mean± standard error of the mean. *Red:* 0-5°, *green*: 5-10°, blue: 10-20°, see inset. *0M:* newborns (n = 23), *3M:* 3 month-olds (n = 23), *6M:* 6 month-olds (n = 21), *A*: adults (n = 21). *Bottom horizontal lines:* cytoarchitectonic area.

We then quantified the distributions of endpoint connections to EVC for each fROI and age group. Consistent with the predictions of the eccentricity-bias hypothesis, in all age groups, pFus-faces, pOTS-words, mFus-faces, OTS-bodies, and mOTS-words have more connections to the EVC 0-5° eccentricity band than the more peripheral eccentricities (Fig 3C) and CoS-places has more connections to the EVC 10-20° eccentricity band than to the more central eccentricities, which may be supported by the medial occipital longitudinal tract^61^ (Fig 3C). Additionally, the connections from VTC to EVC by eccentricity band also mirror the cytoarchitectonic organization of the VTC connectivity profiles. Specifically, across all age groups: (i) pFus-faces and pOTS-words, which are located in cytoarchitectonic area FG2, have ∼60% of their connections to the EVC 0-5° eccentricity band, whereas (iii) CoS-places, which is located in FG3, has ∼40% of its connections to the EVC 10-20° eccentricity band, and (iii) mFus-faces, OTS-bodies, and mOTS-words in FG4 have 40-50% of their EVC connections in the central 5° (Fig 4C).

**Figure 4.**
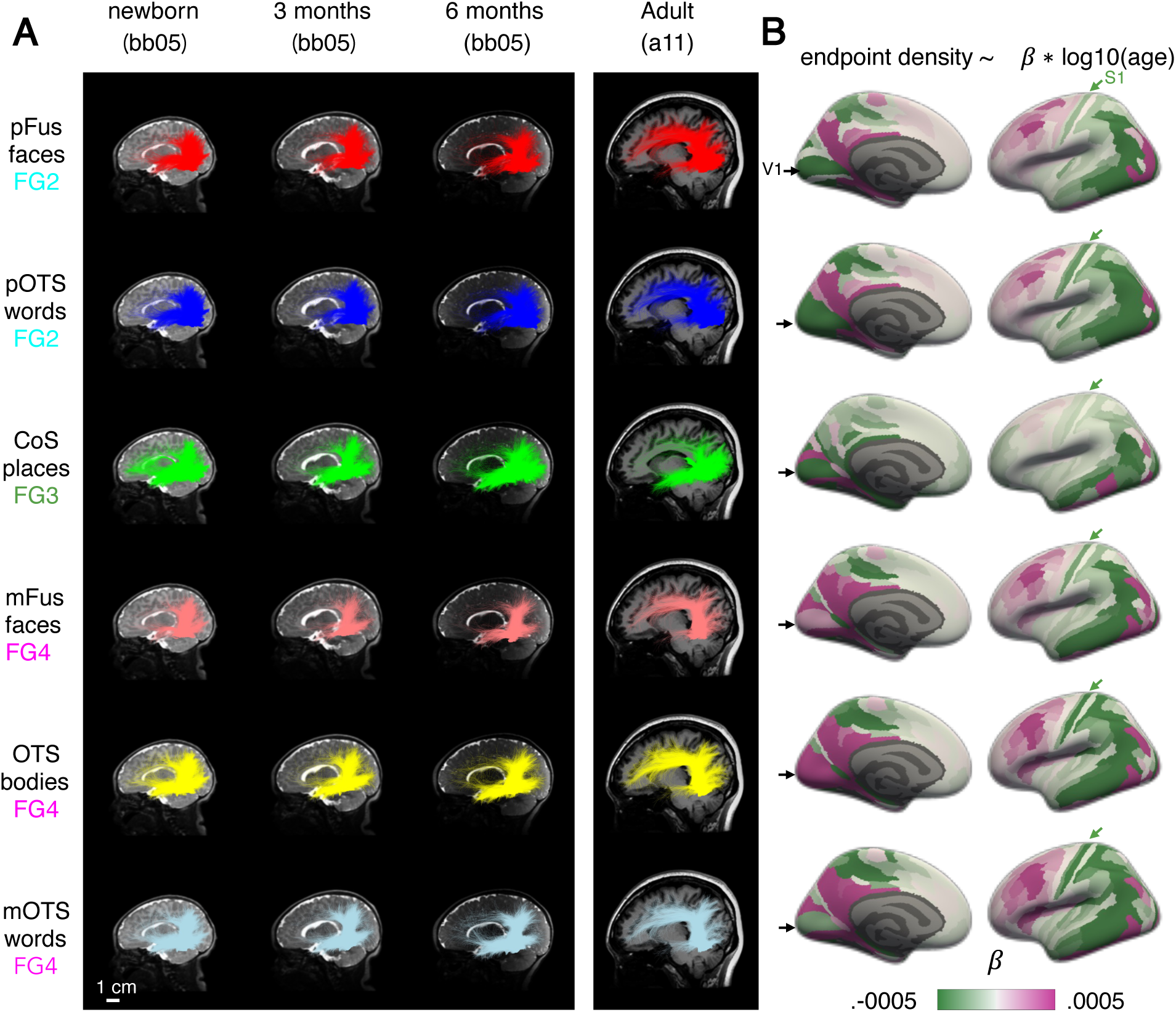
Connectivity of VTC develops from infancy to adulthood. (A) Connections of each fROI in an individual infant across three time points (newborn, 3 months, 6 months) next to an example adult shown in a sagittal cross section. Brains are shown to scale, scale bar, bottom left. Each row is a different fROI ordered by cytoarchitectonic area; from top to bottom: FG2 to FG4. (B) Change in endpoint density over development for each fROI’s connectivity profile. The color of the Glasser ROI indicates the development in endpoint density over time, namely the slope of the regression: endpoint density ∼ log10(age in days). *Green*: decreasing endpoint density over development, *magenta:* increasing endpoint density over development. *Black arrow:* primary visual area (V1); *Green arrow:* primary somatosensory area (S1). Each row shows the development of white matter connections of an fROI; rows ordered identical to (A). Here we show data for the left hemisphere; data for the right hemisphere is in Supplementary Fig 55.

We further tested how eccentricity is related to the features of cytoarchitecture, category, and age that we examined in the whole brain analysis. To do so, for each fROI we fit a linear model predicting the percentage of connections to the central 5° by cytoarchitecture, category-selectivity, and age group. As the percentage of connections to each eccentricity band is proportional, we use the percentage of connections to the central 5° as the independent variable, where a greater percentage connections to the central 5° indicate more connections to central eccentricity bands, and fewer connections to the central 5° indicate more connections to peripheral eccentricity bands.

We find a main effect of cytoarchitecture (F(2,504) = 390.02, p < 0.001, η^2^ = 0.61), suggesting that regions that are in distinct cytoarchitectonic areas have different percentages of connections with the central 5° of EVC. In particular, the majority of connections between FG2 and EVC are in the central 5°, but FG3 has a minority of connections in the central 5° (mean±sd: FG2: 63.57%±14.41%, FG3: 26.58%±8.33%, 4: 43.97%±9.34%). Additionally, there is a main effect of category-selectivity (F(2,504) = 8.32, p < 0.001, η^2^ = 0.03), reflecting that regions with different category-selectivity also have different percentages of connections with the central 5° of EVC. Face- and word-selective regions have a majority of connections with the central 5°, but place-selective regions have only a minority (mean±sd: faces: 51.70%±15.02%, words: 56.21%±15.73%, places: 26.58%±8.33%, bodies: 43.23%±9.05%). In contrast, there is no main effect of age group (F(3,504) = 1.14, p = 0.33, η^2^ = 6.77×10^−3^), and no significant interaction between category and age (F(6,504) = 0.93, p = 0.47, η^2^ = 0.01). There are, however, significant interactions between cytoarchitecture and category (F(1,504) = 24.55, p < 0.001, η^2^ = 0.05), cytoarchitecture and age (F(6,504) = 3.59, p = 0.002, η^2^ = 0.04), and between cytoarchitecture, category, and age (F(3,504) = 2.76, p = 0.04, η^2^ = 0.02). Post-hoc tests reveal that the cytoarchitecture by category interaction is driven by differences in percentage of connections with the central 5° between pFus-faces and pOTS-words in FG2 (t(87) = 5.68, p < 0.001, Bonferroni corrected, Cohen’s d = 0.75, mean difference = −10.09, 95% CI = [−13.63, −6.56]), as well as differences in percentage of connections to the central 5° between mFus-faces and OTS-bodies in FG4 (t(87) = 3.68, p = .0016, Bonferroni corrected, Cohen’s d = 0.17, mean difference = 1.08, 95% CI = [0.76, 2.55]). Further post-hoc tests reveal that differential development is due to a decrease in the percentage of connections from FG4 with the EVC central 5° over development (β = −2.75, 95% CI = [−4.34, −1.16], *t*(163.09) = −3.39, *p* = 0.0027, Bonferroni corrected), and a differential development within FG2 where there is a significant increase in the connections of pOTS-words with central 5° of EVC over development (β = 3.95, 95% CI = [1.75, 6.14], t(81.26) = 3.52, p = 0.0042, Bonferroni corrected), but no significant development in the percentage of pFus-faces connections with the central 5° (β = −0.78, 95% CI = [−4.05, 2.49], *t*(85.90) = −0.47, *p* = 1, Bonferroni corrected). These results suggest that in addition to cytoarchitecture, and category, white matter connections of VTC are organized topographically by eccentricity from birth and that the differential proportion of connections between VTC and different eccentricity bands from birth mirrors the eccentricity bias of both cytoarchitectonic areas and category-selective regions in VTC.

### Do Connections of VTC Develop from Infancy to Adulthood?

In addition to revealing the organizing principles of white matter connections by cytoarchitecture, category, and eccentricity, we also found evidence for development: we were able to classify the age group of the participant above chance from the whole connectivity profile, and we found some developmental changes in connectivity with EVC. To qualitatively assess how white matter connections of VTC change over development, we visualized the white matter connections of all category-selective fROIs in all participants. Fig 4 shows an example infant at newborn, 3 months, and 6 months as well as an example adult (connections of all participants in Supplementary Figs 15-54).

Consistent with the prior analyses, within an age group, white matter connections of fROIs in the same cytoarchitectonic area were similar. For example, in the example newborn, pFus-faces and pOTS-words, both in FG2, are characterized by vertical connections to dorsal occipital cortex and longitudinal connections to both the anterior temporal lobe and early visual cortex (Fig 3A, top 2 rows). However, we also observe developmental changes. For example, pFus-faces and pOTS-words appear to have more abundant vertical connections in infants than adults, but more connections to the frontal cortex in adults than infants (Fig 3A, Supplementary Figs 23-30, 51-54 all participants). We see a similar pattern for the white matter connections of fROIs in FG4: mFus-faces, OTS-bodies, and OTS-words show similar connectivity profiles at each timepoint (e.g., newborn), yet we also observe developmental differences: vertical connections appear more abundant in newborns than adults, but connections to the frontal cortex appear to be more abundant in adults than infants (Fig 3A, bottom row, Supplementary Figs 15-22, 31-38,47-50). For CoS-places, located medially in FG3, there is a different connectivity profile and a different developmental pattern: in newborns, CoS-places has abundant frontal, vertical, and longitudinal connections to both occipital and anterior temporal lobes, but in adults there appear to be fewer frontal and vertical connections (Fig 3A-third row, Supplementary Figs 39-46).

To quantify developmental changes, we tested whether each fROI’s endpoint density, quantified as the proportion of connections in each Glasser ROI, varies as a function of age (fROI endpoint density ∼ log10(age in days) x Glasser ROI, linear model). As expected, fROIs’ endpoint density vary across the brain (significant main effect of Glasser ROI; *F*s > 74.94, *p*s < 0.001; statistics in Supplementary Table 1) reflecting that fROIs have non-uniform connectivity across the brain. Additionally, fROIs’ endpoint density differentially vary with age across the brain (significant age by Glasser ROI interaction; *F*s > 3.54, *p*s < 0.001; statistics in Supplementary Table 1; repeated with correction for gestational age in Supplementary Table 2). As we find a differential development across the brain, for each fROI, we calculate the change in endpoint density across age per Glasser ROI (endpoint density ∼ log10(age in days), linear model). Then, we visualize the slope of this model for each Glasser ROI (Fig 4D). A positive slope (magenta) indicates that the endpoint density within the Glasser ROI increases from infancy to adulthood, whereas a negative slope (green) indicates that endpoint density decreases with age.

This analysis reveals both developmental increases and decreases in endpoint density with distinct spatial characteristics. First, there are common developmental patterns for face, word, and body fROIs (Fig 4-right): endpoint densities to lateral occipital and dorsal occipito-parietal visual Glasser ROIs as well as somatosensory areas decrease (Fig 4-green arrows), but endpoint densities to lateral prefrontal Glasser ROIs increase (Fig 4). There is also differential development of white matter connections of fROIs in FG2 (pOTS-words, pFus-faces) compared to FG4 (mFus-faces mOTS-words, and OTS-bodies): the former show developmental decreases in endpoint densities in V1 and early visual cortex more broadly (Fig 4-medial view) but the latter show developmental increases. Finally, for CoS-places, which is in FG3, we find mostly developmental decreases in endpoint densities in both visual and orbitofrontal cortex with increases in endpoint densities in ventral temporal lateral regions.

Finally, we tested whether development was more similar for fROIs with the same cytoarchitectonic area or same category-selectivity. We used a bootstrapping procedure to estimate the developmental slopes (Methods) and evaluated the similarity between white matter development by calculating the pairwise correlation between the development slopes across Glasser ROI for each pair of fROIs. We then fit a linear model to test whether the correlation was higher for fROIs with the same cytoarchitecture or category-selectivity (linear model: Fisher transformed correlation ∼ cytoarchitecture + category, adjusted R^2^ = 0.81). We find a significant main effect of cytoarchitecture (t(14997) = 248.0, p < 0.001, 95% CI = [0.88, 0.89]), reflecting that regions within the same cytoarchitectonic area had more similar developmental slopes (correlation±sd: .78±11) than regions in different cytoarchitectonic areas (correlation±sd: .25±13). There was also a significant main effect of category (t(14997) = 32.7, p < 0.001, 95% CI = [0.14, 0.16]). However, here we find that regions with the same category-selectivity had less similar developmental slopes (correlation±sd: .36±16) than regions with different category-selectivity (correlation±sd: .40±.28). These analyses suggest that although the relationship between white matter and cytoarchitecture is consistent over development, connectivity profiles develop in parallel within each cytoarchitectonic area, with both developmental increases and decreases in endpoint densities.

## Discussion

Here, we use anatomical and diffusion MRI in infants within the first six months of life and in adults to determine the organizing principles and developmental trajectories of the white matter connections of regions in VTC. We find that white matter connections are organized by cytoarchitecture, eccentricity, and category from birth. We also find evidence for development of white matter connections of VTC, with increasing endpoint density to lateral frontal cortex, and decreasing endpoint density to lateral occipital, parietal, somatosensory, and orbitofrontal cortex. These findings have important implications for understanding the interplay between nature and nurture on both white matter connections and functional brain development.

### Cytoarchitecture and eccentricity are organizing principles

Here, we tested theories that suggest that innate white matter connections support the organization of VTC. When we use the term innate, we refer to the organization of white matter connections at birth, prior to the experience of patterned vision outside the womb. A predominant theory suggests that the consistent organization of category-selective regions in VTC is due to innate white matter connections that support category-specific processing^11,12,15,31^. While we find that in infancy white matter connections are organized to a certain extent by category, especially for faces and places, several of our empirical findings suggest that cytoarchitecture and eccentricity are more parsimonious organizing principles of white matter connectivity from infancy to adulthood. Even as both category and cytoarchitecture can be classified from white matter connectivity profiles irrespective of age, classification of cytoarchitecture is higher than category-selectivity. In addition, the subset of connections between VTC and early visual cortex are organized topographically by eccentricity bands and show similar eccentricity biases for functional regions that are selective for different categories but located in the same cytoarchitectonic area. Finally, connectivity profiles of functional regions in the same cytoarchitectonic area largely develop in parallel from infancy to adulthood.

This organization by cytoarchitecture and eccentricity suggests that these are general principles underlying the white matter organization of the visual system, which may predate and constrain the coupling of white matter connections and brain function. That is, our data suggest a theoretical shift: rather than humans being born with specialized connections to support category-specific processing, cytoarchitectonic boundaries and eccentricity gradients are the innate organizing principles of white matter connections of the visual system. Indeed, in children and adults cytoarchitecture and eccentricity are linked as the boundary between lateral and medial cytoarchitectonic areas in VTC align to the transition between foveal and peripheral representations in VTC^20,57,58,62^. As we find that white matter connections are organized by eccentricity from birth, one functional prediction is that eccentricity biases in VTC^7,8^ will also be present from birth. Nonetheless, we underscore that white matter connections start developing before birth during the second and third trimesters of gestation^63^. Thus, an important direction for future research is understanding how the prenatal environment in the womb^63^ and retinal waves^64^ prior to birth contribute to the organization of white matter connections of the visual system at birth.

As cytoarchitectonic^35,59,65–68^ and retinotopic organization^7,8,21,69–71^ are prevalent across the entire visual system, we hypothesize that these two organizing principles may underlie the white matter organization of the visual system more broadly. This hypothesis can be tested in future research, for example, in the parietal cortex that contains a series of retinotopic^72,73^ and cytoarchitectonic^66^ areas. Critically, as cytoarchitecture^35,60,7435,75–78^, and topography^79,80^ are also key features distinguishing brain areas beyond the visual system, e.g., language^81^, our findings raise the possibility that white matter connections are innately organized by cytoarchitecture and topographic gradients even beyond the visual system and throughout the entire brain. This hypothesis can be tested in future research leveraging large anatomical and diffusion MRI datasets that are being collected in infants^82–86^.

### Development of white matter connections of VTC

While we find innate organization of white matter connections, we also find that the connectivity profiles of fROIs within the same cytoarchitectonic area develop in parallel from infancy to adulthood. Endpoint density from VTC to the frontal lobe showed both developmental increases and decreases. Endpoint density from FG2 and FG4 in lateral VTC to prefrontal cortex increased over development but from FG3 in medial VTC to orbitofrontal cortex decreased. The former finding is consistent with observations that the arcuate fasciculus^87^ is underdeveloped in infants, only reaching the precentral gyrus and premotor cortex in newborns, and not reaching lateral prefrontal cortex as it does in adults^50,88^.

Additionally, we find developmental decreases in endpoint density between VTC and lateral occipital, parietal, and somatosensory cortex. Our observations in the human visual system mirror results from a large body of tracer studies in cats and macaques that documented exuberant connections between V1, V2, and V3^89–91^ as well as between early visual areas and somatosensory areas^92^. These exuberant connections are present early in development and then eliminated by adulthood^90,92–94^. Our data not only suggest that exuberant connections may exist in the human visual system, but also suggest the possibility that connections between VTC and both the lateral and dorsal visual streams^17,95–98^ may decrease over development.

We acknowledge that our *in-vivo* dMRI measurements do not enable us to make inferences about the cellular and molecular underpinnings of the development of white matter connectivity profiles, and there are multiple possible underlying causes of the developmental effects that we observed. For example, increases in connectivity may reflect a pathway that exists at birth and becomes myelinated over development^45^, or a pathway that exists at birth and becomes expanded over development^49^. Likewise, the observation of decreasing connections may reflect decreases in connectivity or increases in crossing fibers. Future studies can examine if developmental increases in endpoint density may be related to increased myelination^99–102^, axonal sprouting^103–105^, and glial proliferation^100,102^ that are associated with activity-dependent white matter plasticity^106,107^. Additionally, future studies can test whether developmental decreases in endpoint density may be related to elimination of exuberant long range connections^90,92–94^.

### Implications for theories of functional development of VTC

Our findings that the white matter connections of VTC have both innate organization and the capacity to change over development raise questions about the relationship between white matter and function in atypical development. One question is whether lack of visual experience may reshape white matter connectivity profiles of VTC and consequently function^14^. For example, it is unknown if lack of visual experience may lead to the preservation of connections between VTC and somatosensory cortex, which in turn, may enable VTC to respond to haptic inputs in individuals who are congenitally blind^108^. Critically, our framework enables measuring the fine-grained white matter connections of VTC in individual infants longitudinally^109^, increasing accuracy and precision in measuring the interplay between white matter connections, functional regions, and anatomy across development. Our framework thus opens opportunities not only for evaluating development of white matter associated with functional regions in large infant datasets^82–86^ but also for early identification and assessment of developmental disorders associated with VTC such as autism^110–113^, Williams syndrome^114^, congenital prosopagnosia^115,116^, and dyslexia^117,110–113^.

Together, the present study advances our understanding of white matter organization in the visual system suggesting that cytoarchitecture, eccentricity, and category are organizing principles from birth, even as aspects of the white matter develop from infancy to adulthood. These data have implications not only for theories of cortical functional development, but also have ramifications for early identification of atypical white matter development.

## Methods

### Participants

The study was approved by the Institutional Review Board of Stanford University and complies with all ethical regulations. Adult participants and parents of infant participants provided written informed consent prior to their scan session. Both infant and adult participants were paid $25/hour for participation.

### Expectant parent and infant screening procedure

Expectant parents and their infants in our study were recruited from the San Francisco Bay Area using social media platforms. We performed a two-step screening process. First, parents were screened over the phone for eligibility based on exclusionary criteria designed to recruit a sample of typically developing infants. Second, eligible expectant mothers were screened once again after giving birth. Exclusionary criteria were as follows: recreational drug use during pregnancy, significant alcohol use during pregnancy (> 3 instances of alcohol consumption per trimester; more than 1 drink/instance), taking prescription medications for a disorder involving psychosis or mania during pregnancy, insufficient written and spoken English ability to understand study instructions, or learning disabilities. Exclusionary criteria for infants were preterm birth (<37 gestational weeks), low birthweight (<5 lbs 8 oz), congenital, genetic, and neurological disorders, visual problems, complications during birth that involved the infant (e.g., NICU stay), history of head trauma, and contraindications for MRI (e.g., metal implants).

43 full term infants following a typical pregnancy (gestational weeks: 39.31±1.32, mean±sd) participated in a total of 75 MRI sessions. Two sessions were excluded for missing diffusion data, and six sessions were excluded for too much motion (see Quality Assurance). We report data from 42 infants over 67 sessions (21 longitudinal): sex: 16 female, 26 male; race and ethnicity: 4 Asian, 5 Hispanic, 11 Multiracial, and 22 White participants; age: n=23 newborns (28.56 ± 10.21days, mean±sd), n=23 3-month-olds (106.91 ± 19.33 days), and n=21 6-month-olds (189.05 ±15.77 days).

We also collected data from 21 adults: sex: 17 female, 4 male; race and ethnicity: 12 Asian, 1 Hispanic, 2 Multiracial, and 6 White participants; age: M=28.21 years, SD=5.51 years. All adult sessions met inclusion criteria. No statistical methods were used to pre-determine sample sizes, but our sample sizes are similar to previous publications^45^.

### MRI acquisitions

Scanning sessions were scheduled in the evenings around infants’ bedtime and were done during natural sleep. Infant data were acquired on a 3T GE Ultra High Performance (UHP) scanner (GE Healthcare, Waukesha, WI) equipped with a customized 32-channel infant head-coil^118^.

Hearing protection included soft wax earplugs, and MRI compatible neonatal noise attenuators (https://newborncare.natus.com/products-services/newborn-care-products/nursery-essentials/minimuffs-neonatal-noise-attenuators), and headphones (https://www.alpinehearingprotection.com/products/muffy-baby) that covered the infant’s ears. During sessions with newborns, an MR-safe plastic immobilizer (MedVac, www.supertechx-ray.com) was used to stabilize the infant and their head position. When the infant was asleep, the caregiver placed the infant on the scanner bed. Weighted bags were placed at the edges of the bed to prevent side-to-side movements. Pads were also placed around the infant’s head and body to stabilize head position. An experimenter stayed inside the MR suite with the infant during the entire scan.

During scan, experimenters monitored the infant using an infrared camera that was affixed to the head coil and positioned for viewing the infant’s face. Experimenters stopped the scan if the infant showed signs of waking or distress or excessively moved; scans were repeated if there was excessive head motion.

### Anatomical MRI acquisition

We obtained T1-weighted MRI data (GE’s BRAVO sequence) for each infant and adult session with the following parameters: TE = 2.9ms; TR = 6.9ms; voxel size = 0.8x 0.8 x0.8 [mm^3^]; FOV = 20.5 cm; Scan time: 3:05 min. We obtained T2-weighted MRI data for each infant session as the T2-weighted contrast between different tissue types is better than T1-weighted images for young infants. T2-weighted (GE’s CUBE sequence) parameters: TE = 124ms; TR = 3650 ms; voxel size = 0.8x 0.8x 0.8 [mm^3^]; FOV = 20.5 cm; Scan time: 4:05 min.

### Anatomical MRI processing for infant sessions

The T1-weighted and T2-weighted images from each individual were aligned to a plane running through the commissures (AC-PC transformed)using rigid body alignment (FSL FLIRT^119^; https://web.mit.edu/fsl_v5.0.10/fsl/doc/wiki/FLIRT(2f)UserGuide.html). Then, we used iBEAT V2.0^120^ to segment the white and gray matter. White matter segmentation was then manually fixed for errors using ITKgray^121^. We used the manually edited segmentation file to reconstruct the cortical surface with Infant FreeSurfer^122^.

### Anatomical MRI processing for adult sessions

The T1-weighted images were aligned to a plane running through the commissures (AC-PC transformed). Tissue segmentation was done with Freesurfer v7.2^123^ .

### Diffusion MRI (dMRI) acquisition

dMRI used the following parameters: multi-shell, #diffusion directions/b-value = 9/0, 30/700, 64/2000; TE = 75.7 ms; TR = 2800 ms; voxel size = 2×2×2 [mm^3^]; number of slices = 60; FOV = 20 cm; in-plane/through-plane acceleration = 1/3; scan time: 5:08 min. We also acquired a short dMRI scan with reverse phase encoding direction and only 6 b = 0 images.

### Diffusion MRI processing

dMRI data was preprocessed using MRtrix3^124^ (https://github.com/MRtrix3/mrtrix3) in accordance with prior work from the human connectome project and our lab ^45,125,126^. Data were denoised using principal component analysis^127^. We used FSL’s top-up tool (https://fsl.fmrib.ox.ac.uk/) and one image with reverse phase-encoding to correct for susceptibility-induced distortions. We used FSL’s eddy tool to perform eddy current and motion correction, where outlier slices were detected and replaced^128^. Finally we performed bias correction using ANTS^129^ (https://picsl.upenn.edu/software/ants/). The preprocessed dMRI data were aligned to the T2-weighted anatomy for infants, and T1-weighted anatomy for adults using whole-brain rigid body registration. Alignment was checked manually.

### Generating White Matter Connectomes

We generated a whole brain white matter connectome in each session using MRTrix3^124^. Voxel-wise fiber orientation distributions (FODs) were calculated using constrained spherical deconvolution (CSD). We used the Dhollander algorithm^130^ to estimate the three-tissue response function. We computed FODs separately for the white matter and CSF. As in past work^45,125^, the gray matter was not modeled separately, as white and gray matter do not have sufficiently distinct b-value dependencies to allow for a clean separation of the signals. Finally, we generated a whole brain white matter connectome for each session using MRTrix3 tckgen. We used a FOD amplitude threshold of 0.05. Tractography was optimized using the gray/white matter segmentation from anatomical MRI data (Anatomically Constrained Tractography; ACT^131^). For each connectome, we used probabilistic fiber tracking with the following parameters: iFOD2 algorithm, step size of 0.2 mm, minimum length of 4 mm, maximum length of 200 mm, and maximum angle of 15°. Streamlines were randomly seeded on this gray-matter/white-matter interface, and each connectome consisted of 5 million streamlines. This procedure is critical for accurately identifying the connections that reach fROIs, which are in the gray matter^132^.

### Quality Assurance

To evaluate the quality of dMRI data we implemented three quality assurance measures: (i) We measured the number of outliers (dMRI volumes with signal dropout measured by FSL’s eddy tool). The exclusion criterion was >5% outlier volumes as in past work^20^. (ii) We visualized in mrView the fractional anisotropy (FA) colored by direction of the diffusion tensors to validate the expected maps. That is, the existence of between hemispheric connections through the corpus callosum, and the inferior-superior directionality of the cerebral spinal tract. (iii) We used babyAFQ^22^ to identify the bundles of the brain and validate that the major bundles known to be present at birth^45,46,133^ can be found in each session.

These quality assurance measures lead to the following data exclusions:

i. No adult sessions were excluded for outliers. Six infant sessions were excluded because they had >5% outliers. After exclusion, there were no significant differences in outlier volumes between infants and adults (infants: mean percent outliers±SD = 0.45% ± 0.43%; adults: mean percent outliers±SD = 0.29% ± 0.10%; t(86) = 1.70, p = .092).
ii. We identified the corpus callosum with left-right directionality and corticospinal tract with inferior-superior directionality in all infant and adult sessions; no data was excluded by this measure.
iii. We identified the major bundles in the infant brain using babyAFQ, and in adults using AFQ. In each infant and adult session, we identified the arcuate (AF), posterior arcuate (pAF), vertical occipital (VOF), inferior longitudinal (ILF), superior longitudinal (SLF), corticospinal (CST), cingulate (CC), forceps major (FcMa), and forceps minor (FcMi), as such, no data were excluded. This confirms the quality of dMRI data and our tracking procedure as we could identify the major white matter bundles that are known to exist from birth in all sessions.

### Cytoarchitectonic Areas of VTC

We used the Rosenke Atlas^59^ that contains maximum probability maps (MPMs) of 8 cytoarchitectonic regions of the human ventral visual stream, created from 10 post mortem adult samples published in^37,134^. We used cortex based alignment in FreeSurfer^56^ (https://freesurfer.net/, mri_label2label) to map the MPM of four cytoarchitectonic areas of the ventral stream FG1, FG2, FG3, FG4 to each cortical surface for each of the 88 sessions in the study. Two independent raters (SD, DO) manually checked each brain to examine if the mid fusiform sulcus (MFS) was the boundary between FG3/FG4 and FG1/FG2 as reported in prior studies^5,58^. In places where there were disagreements between the raters, data was checked by KGS. The analysis confirmed that the boundaries between the cytoarchitectonic areas aligned with the MFS (see examples in Fig 1c) except for five infants in the right hemisphere, and one adult in the left hemisphere.

### Category-Selective Regions of VTC

A functional atlas of category-selective regions of VTC was created from independent fMRI data in 28 adults (11 females and 17 males, ages 22.1–28.6 years, mean±SD = 24.1±1.6 years)from a previous study^20^. Category-selective regions were defined in each of the individuals using the fLoc experiment containing low-level and familiarity-controlled gray level images of items from 10 visual categories^135^. Category-selective regions were defined using a voxel-level t-statistic contrasting each category of interest with all other categories (t > 3, no spatial smoothing), and anatomical criteria^5,6,135^. We defined two face-selective regions on the fusiform gyrus of each hemisphere, mFus-faces and pFus-faces (contrast: faces > bodies, limbs, characters, objects, places). mFus-faces is located near the anterior tip of the mid fusiform sulcus (MFS), pFus-faces is in the posterior fusiform. We defined OTS-bodies in both hemispheres as the region on the occipital temporal sulcus selective for bodies (bodies and limbs > characters, faces, objects, places). We defined CoS-places in both hemispheres as the region in the intersection between the collateral sulcus (CoS) and anterior lingual sulcus (ALS) that was selective for places (places > faces, characters, objects, bodies, limbs). Finally, we identified mOTS-words and pOTS-words as the regions in the occipital temporal sulcus (OTS) that were selective for words (psuedowords > faces, scenes, objects, bodies, limbs), where pOTS-words in on the posterior end of the OTS lateral to pFus-faces, and mOTS-words is more anterior and lateral to mFus-faces. After identifying the functional regions within each participant, we mapped the regions to the fsaverage template brain space using cortex based alignment^56^. We then created probabilistic maps for each fROI where each vertex was the probability of a participant having the fROI at that location. We thresholded the probabilistic maps at .2 and then created MPMs, where in the case that two fROIs had probabilistic values > .2 at the same vertex, the vertex was assigned to the fROI with the highest probability (as in^136^). MPM-fROIs of word-selective regions were only found in the left hemisphere^137,138^.

To map fROIs to individual participants’ cortical space for the 88 sessions of the main experiment, we used cortex based alignment in Freesurfer^56^ (https://freesurfer.net/, mri_label2label). Two independent raters (DO and SD) then visually checked each surface to ensure that the fROIs were aligned to the expected anatomical landmarks in each individual participant. The raters checked whether mFus-faces aligned to the mid fusiform sulcus (MFS) and whether CoS-places aligned to the junction of the ALS and CoS. When there was disagreement, KGS then checked whether the fROIs aligned to anatomical landmarks. Category-selective fROIs aligned to these anatomical landmarks in all hemispheres except for one infant in which left hemisphere CoS-places did not align to the junction or the ALS and CoS, and two infants in which right hemisphere mFus-faces did not align to the MFS. Supplementary Figs. 2-9 show the mapping of each fROI in each participant.

We tested whether fROI surface area relative to brain surface varied with age as follows:

(1) (fROI surface area)/(brain surface area) ∼ log10(age in days).

We found no significant differences between infants and adults on fROI surface area relative to brain surface area (effect of age: t(878) = 8.77×10^−6^, p = 0.88)), reflecting that fROIs were the same size relative to the size of the brain regardless of age.

### Identifying functionally defined white matter connections

To identify the white matter connections of each fROI, we intersected it with each individual’s whole brain connectome using an open source software package, FSuB-Extractor^109^ (https://github.com/smeisler/fsub_extractor). The software takes in fROIs in the native space of each participant and projects them along the surface normal into the gray-matter-white-matter interface. It then restricts the fROI to the gray-matter-white-matter interface and then selects all streamlines that intersect with the fROI using a radial search with a search distance of 3mm.

### Defining connectivity profiles

For each fROI, we define its connectivity profile. To do so, we took the white matter connections of each fROI, projected their endpoints to the cortical surface using tract density imaging (TDI) with MRTrix3^124^, and calculated the distribution of these white matter endpoints across the cortical surface. We transformed the TDI output into a distribution of endpoints by dividing the endpoint map by the total number of endpoints. This results in an endpoint density map that sums to 1 for each fROI and participant. We then used the Glasser Atlas^60^ to quantify how the endpoints were distributed across the brain. We chose to use the Glasser Atlas because it covers the whole cortical surface of each hemisphere, and because it divides cortex into meaningful parcellations according to functional and anatomical criteria. For each fROI, we define the white matter connectivity profile or the endpoint density in each region in the Glasser Atlas. The Glasser Atlas consists of 180 regions per hemisphere. As in prior work^20^, we excluded 11 VTC regions of the Glasser Atlas to avoid quantifying looping fibers from the seed fROI. Therefore, each connectivity profile consisted of the endpoint density across 169 regions in the Glasser Atlas within the same hemisphere of the fROI.

### Principal Components Analysis

Because our connectivity profiles are high-dimensional (169), we used principal component analysis (PCA function in MATLAB) to reduce the dimensionality of the connectivity profiles. We conducted PCA on the 880 (10 fROIs x 88 sessions connectivity profiles) x 169 (Glasser ROIs) endpoint connectivity matrix. We used find_curve_elbow in R (https://rdrr.io/cran/pathviewr/man/find_curve_elbow.html) to find the elbow of the curve of principal components vs. variance explained. We found that the elbow of the curve was at 10 principal components, and that the first 10 principal components explained 98% of the variance in the data. In Figure 2, we plot the first principal component (explains 38% of the variance) vs the second principal component (explains 20% of the variance). Each dot represents a single connectivity profile in a single subject and session; The connectivity profiles (dots) can be coded by different features (cytoarchitectonic area, category-preference, or participant’s age).

### Classification Analysis

We used a n-way leave-one-out classifier to test if we could predict different features (cytoarchitecture, category, or age) of a held-out connectivity profile. For each feature (cytoarchitecture, category, age), we used multinomial logistic regression (multinom function from the nnet package in R^139^) fit on all data, excluding all sessions from the held-out subject to account for the longitudinal nature of the data to predict the feature of the held out connectivity profile (e.g., probability of belonging to FG2, FG3, or FG4 for the cytoarchitecture classification). We used a winner-take-all approach and assigned the connectivity profile to the label with the highest probability. We repeated the process for each connectivity profile (leave-one-out cross-validation). We then calculated classification accuracy by comparing the predicted classification to the ground truth. We performed three separate classifications: one predicting cytoarchitecture (FG2/FG3/FG4), one predicting category (face/word/body/place), and one predicting age group: (newborn/3 months/6 months/adult).

We compared classification accuracy across classification tasks by using a binomial logistic regression (glm in R (https://www.rdocumentation.org/packages/stats/versions/3.6.2/topics/glm)) with the following model:

(1) classification accuracy(1/0) ∼ classification task (cytoarchitecture/category/age)

As our results could be affected by training sets with different numbers of examples for each label, we used a bootstrapping procedure to create a balanced training set. To do so, we sampled the training set (all data except for the held-out subject) with replacement 200 times for each label (e.g., FG2/FG3/FG4) to create a balanced training set. We then fit a multinomial logistic regression on the balanced training set to predict the feature of the held-out connectivity profile. We then calculated classification accuracy by comparing the predicted classification to the ground truth. We repeated this procedure for the cytoarchitecture, category, and age classification.

### Connections to Eccentricity Bands in Early Visual Cortex

To test how each fROI connects to different eccentricity bands in early visual cortex (EVC), we created a region of interest corresponding to EVC from the union of V1, V2, and V3 ROIs from the Benson Atlas^140^ (https://github.com/noahbenson/neuropythy). We then used the average retinotopic map in 21 adults from a previously published paper^141^ to divide EVC into three eccentricity bands, 0-5°, 5-10°, and 10-20°. Using cortex-based alignment in FreeSurfer (mri_label2label) we aligned each of these EVC eccentricity band ROIs to each individual participant’s native space. We then used FSuB-Extractor ^109^ (https://github.com/smeisler/fsub_extractor), to identify the white matter connections between each fROI and each eccentricity band in VTC. We divided the number of streamlines connecting to each eccentricity band by the total number of streamlines between the fROI and EVC to estimate the percentage of streamlines connected to each eccentricity band. To test if the percentage of streamlines to each eccentricity band differs by cytoarchitecture, category, and age, we fit the following model using the lm function in R (https://www.rdocumentation.org/packages/stats/versions/3.6.2/topics/lm):

(2) percentage of streamlines in central 5° ∼ cytoarchitecture x category x age group

After finding a significant cytoarchitecture by category interaction, we performed post-hoc tests to determine which fROIs were driving the interaction. To do so, we used paired t-tests to test whether the percentage of streamlines in the central 5° differed for fROIs within the same cytoarchitectonic area for the following pairs of fROIs: pFus-faces and pOTS-words in FG2, mFus-faces and OTS-bodies in FG4, mFus-faces and mOTS-words in FG4, and OTS-bodies and mOTS-words in FG4.

After finding a significant cytoarchitecture by age interaction, we performed post-hoc tests to test whether the percentage of streamlines in the central 5° differed as a function of age for fROIs within each cytoarchitectonic area using the following linear mixed model (using lmer function from the lmerTest package^142^ in R) separately for each cytoarchitectonic area (FG2, FG3, FG4), using a random intercept for each subject to account for the longitudinal nature of the data:

(3) percentage of streamlines in central 5° ∼ log10(age) + (1|subject)

After finding a significant interaction between cytoarchitecture, category, and age we performed post-hoc tests to test whether the percentage of streamlines to the central 5° changed over development using the following linear mixed model (using lmer function from the lmerTest package^142^ in R) separately for each fROI (pFus-faces, pOTS-words, CoS-places, mFus-faces, OTS-bodies, mOTS-words), using a random intercept for each subject to account for the longitudinal nature of the data:

(4) percentage of streamlines in central 5° ∼ log10(age) + (1|subject)

As development is expected to asymptote across the lifespan and typically follows a logarithmic function of age, to quantify how white matter connectivity profiles change with age, we fit a linear model (lm in R (https://www.rdocumentation.org/packages/stats/versions/3.6.2/topics/lm)) predicting endpoint density of each fROI as a function of Glasser ROI and log10(age in days) across the brain. Age is a continuous variable and Glasser ROI is a categorical variable. As endpoint density was normalized to sum to one across the brain, it is not appropriate to model a random intercept for each subject.

(5) fROI endpoint density ∼ log10(age in days) x Glasser ROI

After finding a significant Glasser ROI by age interaction (Supplementary Table 1), for each fROI, we fit linear models (lm in R https://www.rdocumentation.org/packages/stats/versions/3.6.2/topics/lm) relating endpoint density vs log10(age) separately for each Glasser ROI to quantify the development of endpoint density within each Glasser ROI. The regression slope quantifies the rate of the development:

(6) fROI endpoint density ∼ log10(age in days).

We visualize the regression results in Fig 4B, where each Glasser ROI is colored by the slope of the regression. Data is reported for the left hemisphere in the main text and Fig 3 because all 6 fROIs are found consistently in the left hemisphere. Data from 4 fROIs in the right hemisphere is reported in Supplementary Figure 55. Supplementary Tables 3-8 provide all the developmental slopes per Glasser ROI.

### Calculating the similarity between developmental slopes

To test whether development was more similar for regions within the same cytoarchitectonic area or category-selectivity we estimated the similarity between developmental slopes of white matter connectivity of fROI pairs using a bootstrapping procedure.

First, we calculated the developmental slopes of endpoint connectivity across the brain using a bootstrapping procedure. For each fROI, and for each of 1000 iterations we used 75% of the data (66 randomly selected connectivity profiles), and calculated the developmental slope within each of the 169 Glasser ROIs using the following model (lm in R https://www.rdocumentation.org/packages/stats/versions/3.6.2/topics/lm):

(7) fROI endpoint density ∼ log10(age in days)

Then for each pair of fROIs we calculated the pairwise correlation between the vector of 169 slopes across the bootstrap iterations. Finally, we fit a linear model (lm in R https://www.rdocumentation.org/packages/stats/versions/3.6.2/topics/lm)) to test whether the Fisher transformed correlation between development slopes was higher for fROIs with the same cytoarchitecture and category-selectivity than for fROIs with different cytoarchitecture and category-selectivity. We calculated the Fisher transform to ensure normality (as correlations are bounded between −1 and 1); category and cytoarchitecture are categorical variables (same: 1; different: 0).

(8) FisherZ(correlation) ∼ category + cytoarchitecture

## Data availability

The data to make the figures, tables, and statistics associated with this manuscript is available here: https://github.com/VPNL/bbVTCwm/tree/main/data

## Code availability

The code to analyze the data, compute statistics, and make the individual figure elements is available here: https://github.com/VPNL/bbVTCwm/. The code folder contains the R code used to generate all other figures and statistics in the figures/ and statistics/ subdirectories. The code used to preprocess the data and perform the analyses are included in the analyses/ subdirectory. The label files for the fROIs and the EVC ROIs are provided in the labels folder. The supplement folder contains code to generate Supplementary Figures.

## Acknowledgements

This work was funded by Stanford Wu Tsai Neurodevelopment big idea and accelerator grants, as well as NIH grants R01EY033835 and R01EY022318 to KGS; the National Science Foundation Graduate Research Fellowship (grant number DGE-1656518) to EK; the Deutsche Forschungsgemeinschaft (DFG, German Research Foundation – project number 222641018 – SFB/TRR 135 TP C10), as well as “The Adaptive Mind’’, funded by the Excellence Program of the Hessian Ministry of Higher Education, Science, Research and Art to MG; the Deutsche Forschungsgemeinschaft (DFG, German Research Foundation – grant INST 169/22–1), the Excellence Program of the Hessian Ministry of Higher Education, Science, Research and Art (grants: 2/16/519/03/09.001(0001)/101 and LOEWE/4TP//519/05/02.002(0004)/107) to BK. The funders had no role in study design, data collection and analysis, decision to publish or preparation of the manuscript.

## Author Contributions

EK: designed the analyses, wrote the code and data analysis pipelines, analyzed the data, and wrote the manuscript. XY: participated in the design and data analysis and collected the data. ST, BF, CT collected the data, segmented each brain anatomy image into gray and white matter, and created cortical surface reconstructions. SF and DO: validated the alignment between fROIs, cytoarchitectonic areas and anatomical landmarks. MG: participated in the data analyses. VSN: participated in the design and data analysis and collected the data. BK: designed the infant coil used for data collection. KGS: oversaw all parts of the research: design, data analysis, and wrote the manuscript. All authors read and gave feedback on the manuscript.

