## Supplementary Fig for "White matter connections of human ventral temporal cortex are organized by cytoarchitecture, eccentricity, and category-selectivity from birth"

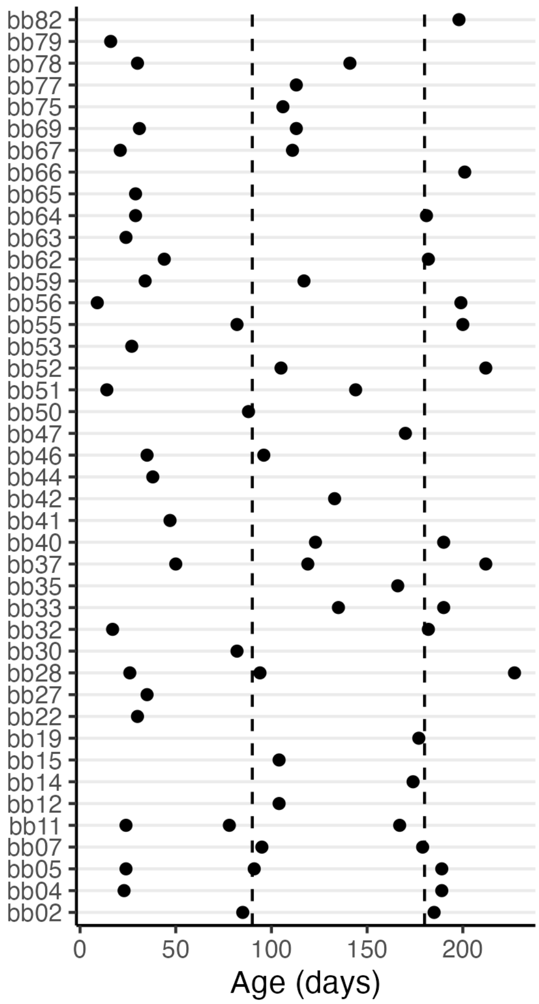


**Supplementary Figure 1.** Summary of the longitudinal visits for each infant participant. X-axis is the age in days at visit, y-axis is the subject number. Each dot represents a visit. Each infant was scanned one, two, or three times with visits near the three month and six month time points (vertical dashed lines at 90 days (~3 months) and 180 days (~6 months).

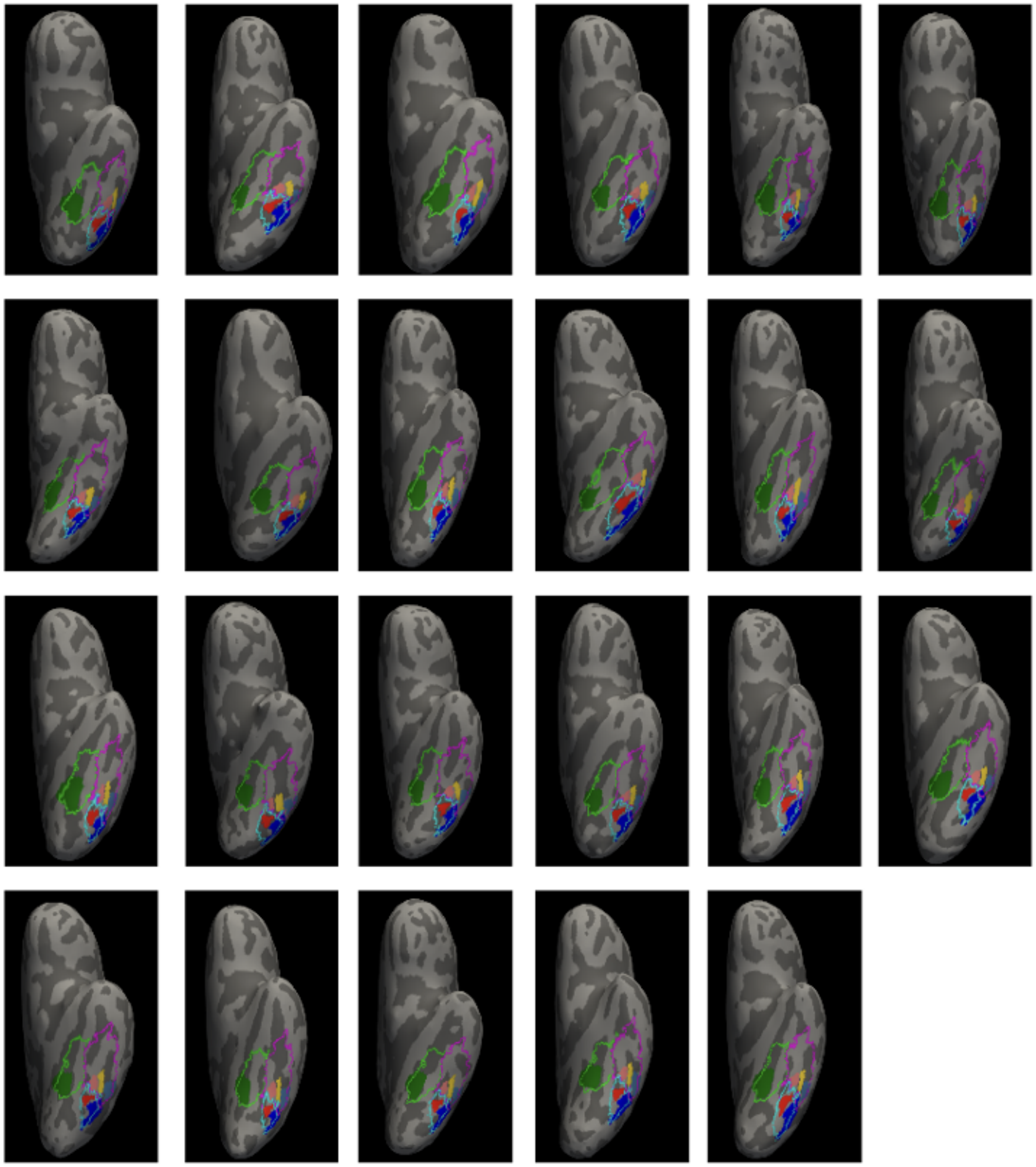

**Supplementary Figure 2.** Left hemisphere maximal probability map (MPM) of functional regions of interest (fROIs) mapped to the brain of each individual infant subject in the 0-month old age group ordered by age from the upper left to the lower right. Colors indicate functional ROI: *green:* CoS-places; *pink:* mFus-faces, *red:* pFus-faces; *yellow:* OTS-bodies; *light blue:* mOTS-words*;* *blue:* pOTS-words. Outlines indicate cytoarchitectonic area: *cyan:* FG2, *green:* FG3, *magenta:* FG4.

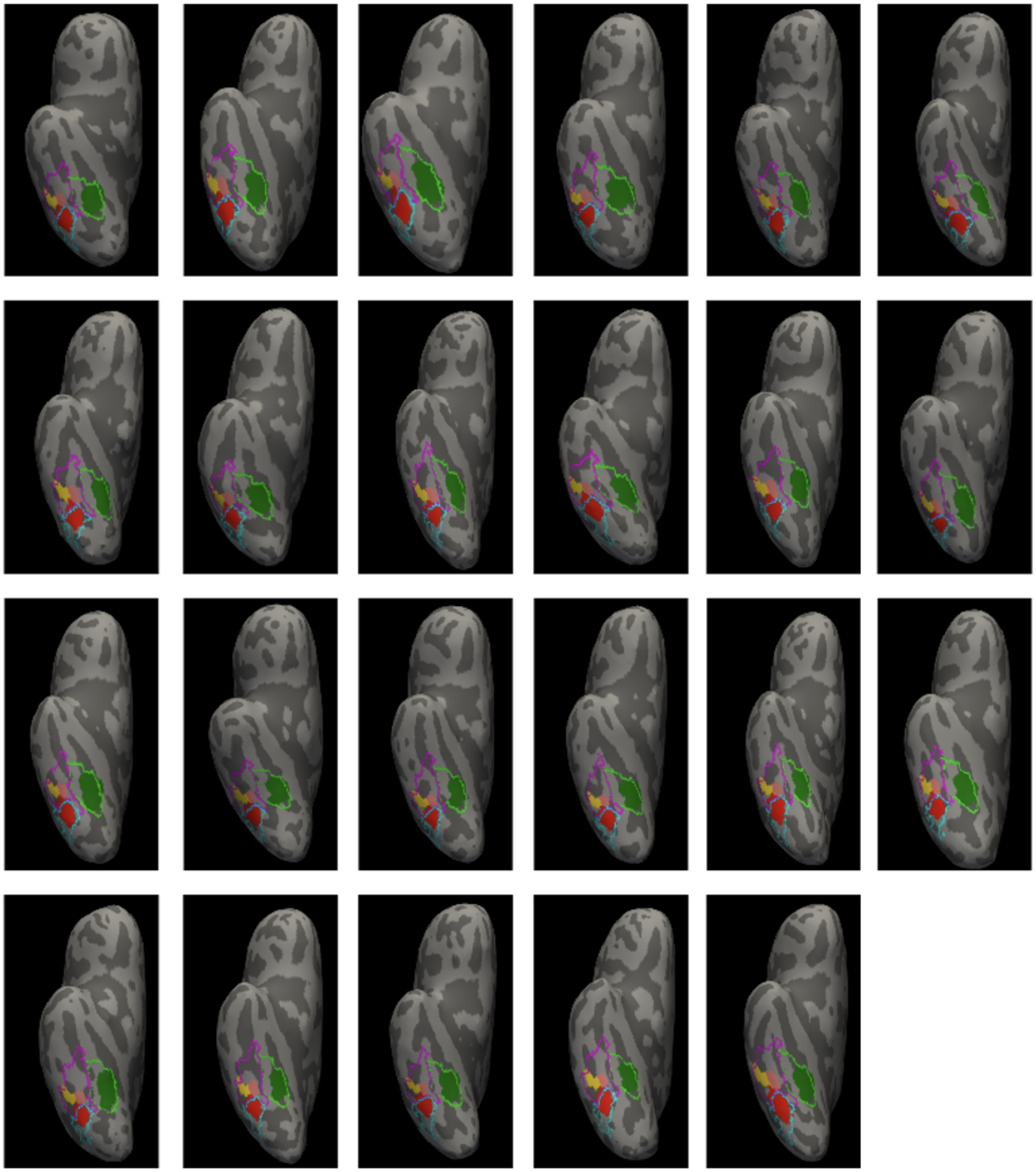

**Supplementary Figure 3.** Right hemisphere MPM ROIs mapped to the brain of each individual infant subject in the 0-month old age group ordered by age. Colors indicate functional ROI: *green:* CoS-places; *pink:* mFus-faces, *red:* pFus-faces; *yellow:* OTS-bodies; *light blue:* mOTS-words; *blue:* pOTS-words. Outlines indicate cytoarchitectonic area: *cyan:* FG2, *green:* FG3, *magenta:* FG4.

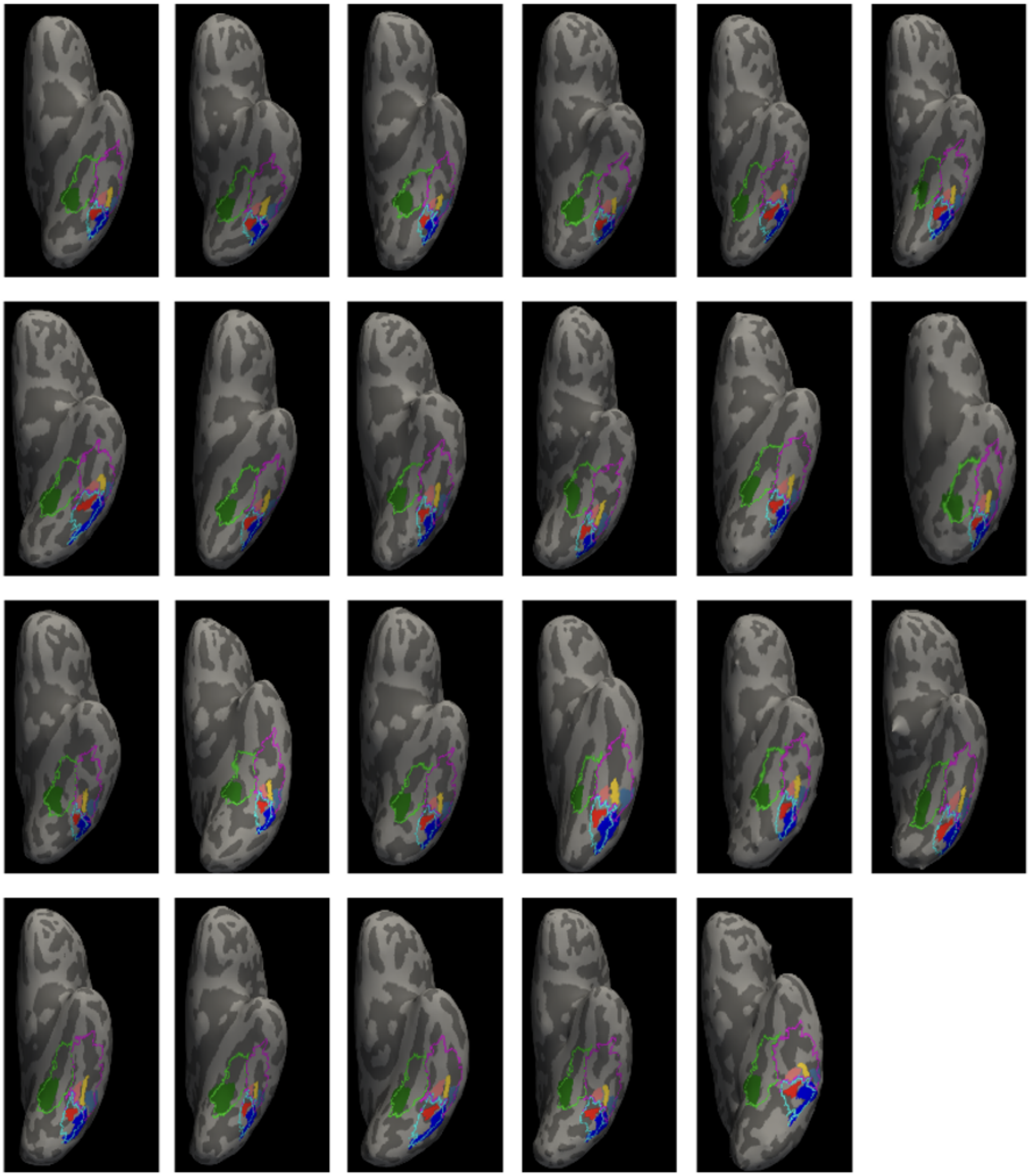

**Supplementary Figure 4.** Left hemisphere MPM fROIs mapped to the brain of each individual infant subject in the 3-month old age group ordered by age. Colors indicate functional ROI: *green:* CoS-places; *pink:* mFus-faces, *red:* pFus-faces; *yellow:* OTS-bodies; *light blue*: mOTS-words; *blue:* pOTS-words. Outlines indicate cytoarchitectonic area: *cyan:* FG2, *green:* FG3, *magenta:* FG4.

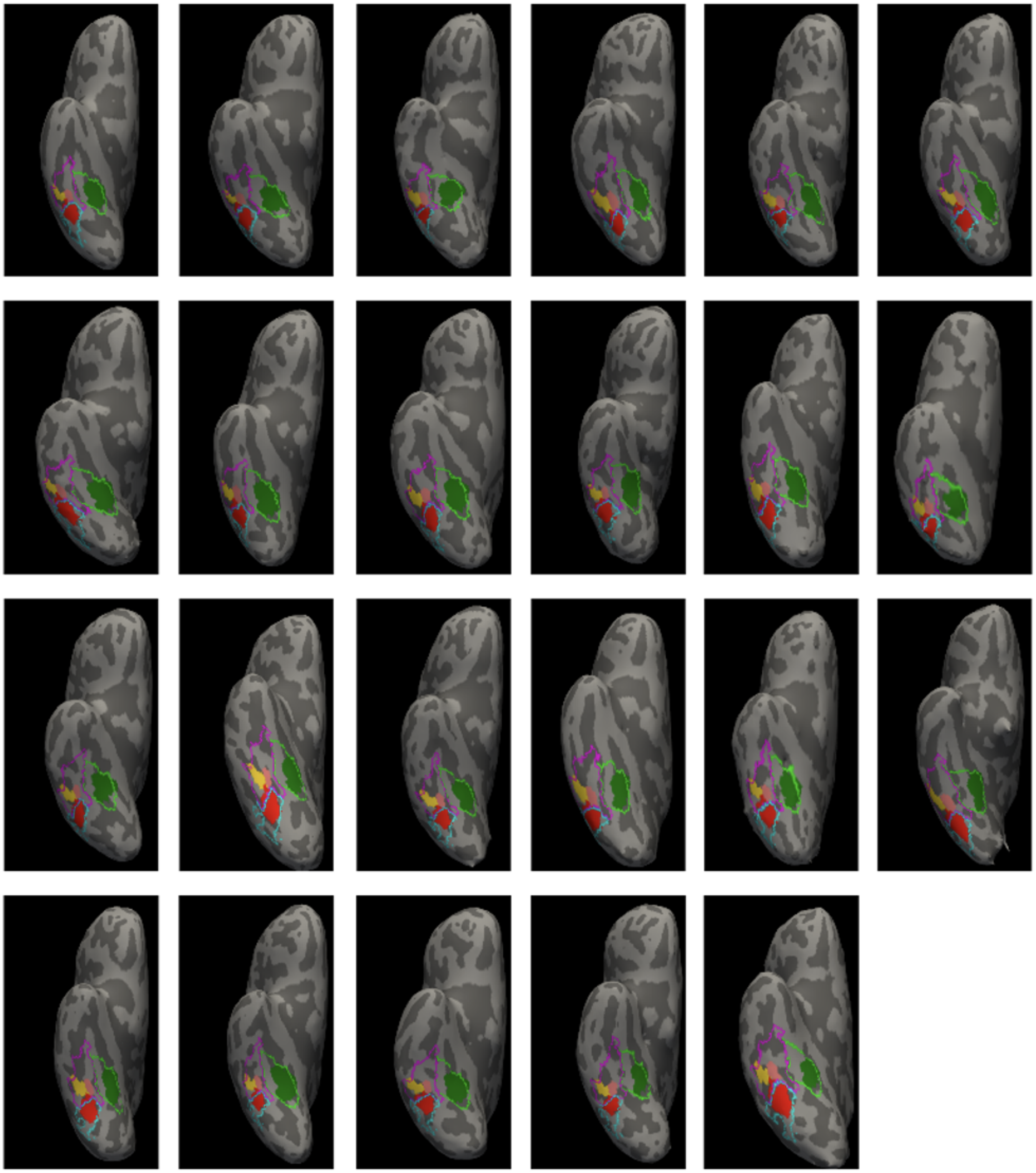

**Supplementary Figure 5.** Right hemisphere MPM ROIs mapped to the brain of each individual infant subject in the 3-month old age group ordered by age. Colors indicate functional ROI: *green:* CoS-places; *pink:* mFus-faces, *red:* pFus-faces; *yellow:* OTS-bodies; *light blue:* mOTS-words; *blue:* pOTS-words. Outlines indicate cytoarchitectonic area: *cyan:* FG2, *green:* FG3, *magenta:* FG4.

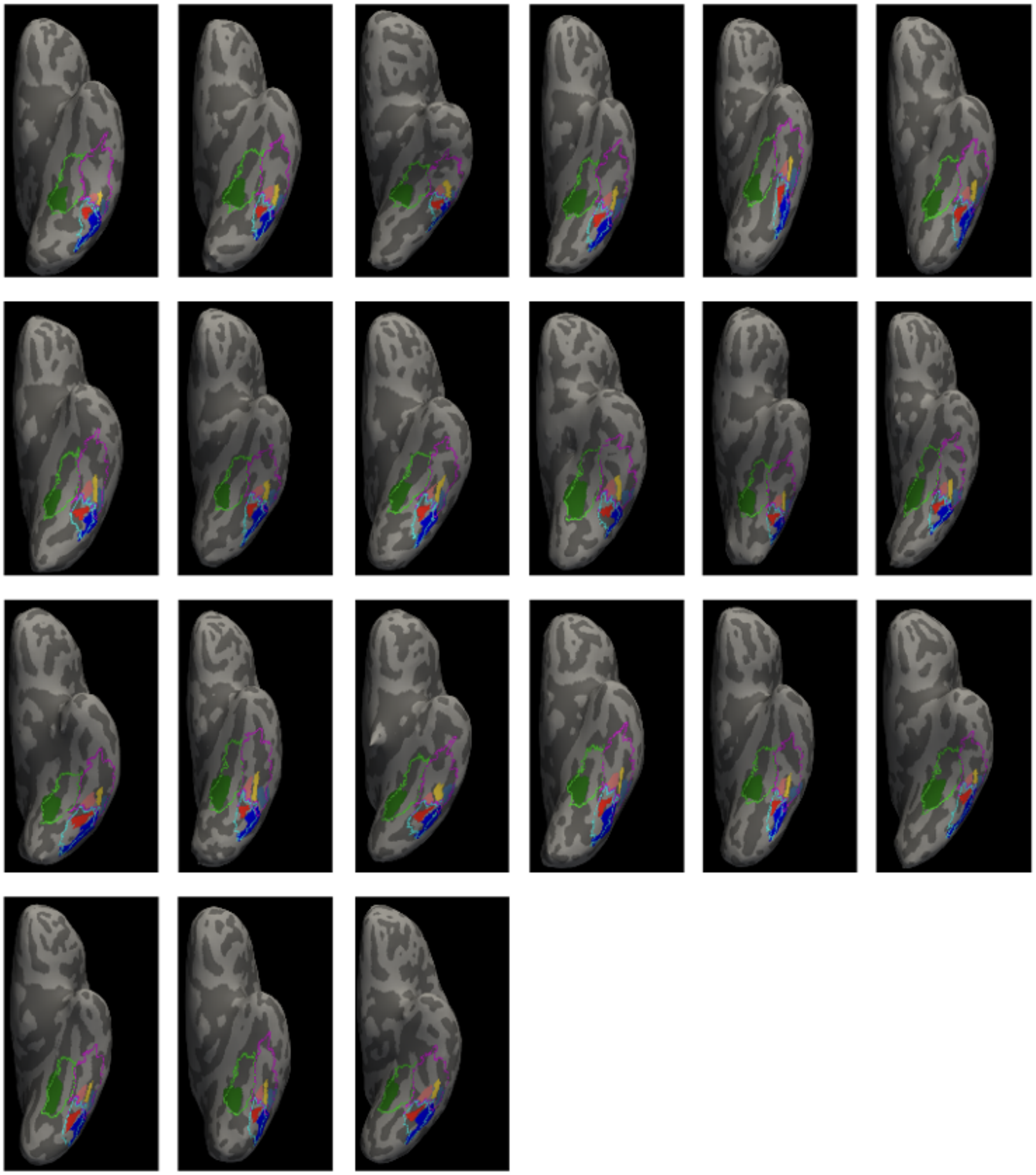

**Supplementary Figure 6.** Left hemisphere MPM fROIs mapped to the brain of each individual infant subject in the 6-month old age group ordered by age. Colors indicate functional ROI: green: CoS-places; pink: mFus-faces, red: pFus-faces; yellow: OTS-bodies; light blue: mOTS-;words; blue: pOTS-words. Outlines indicate cytoarchitectonic area: *cyan:* FG2, *green:* FG3, *magenta:* FG4.

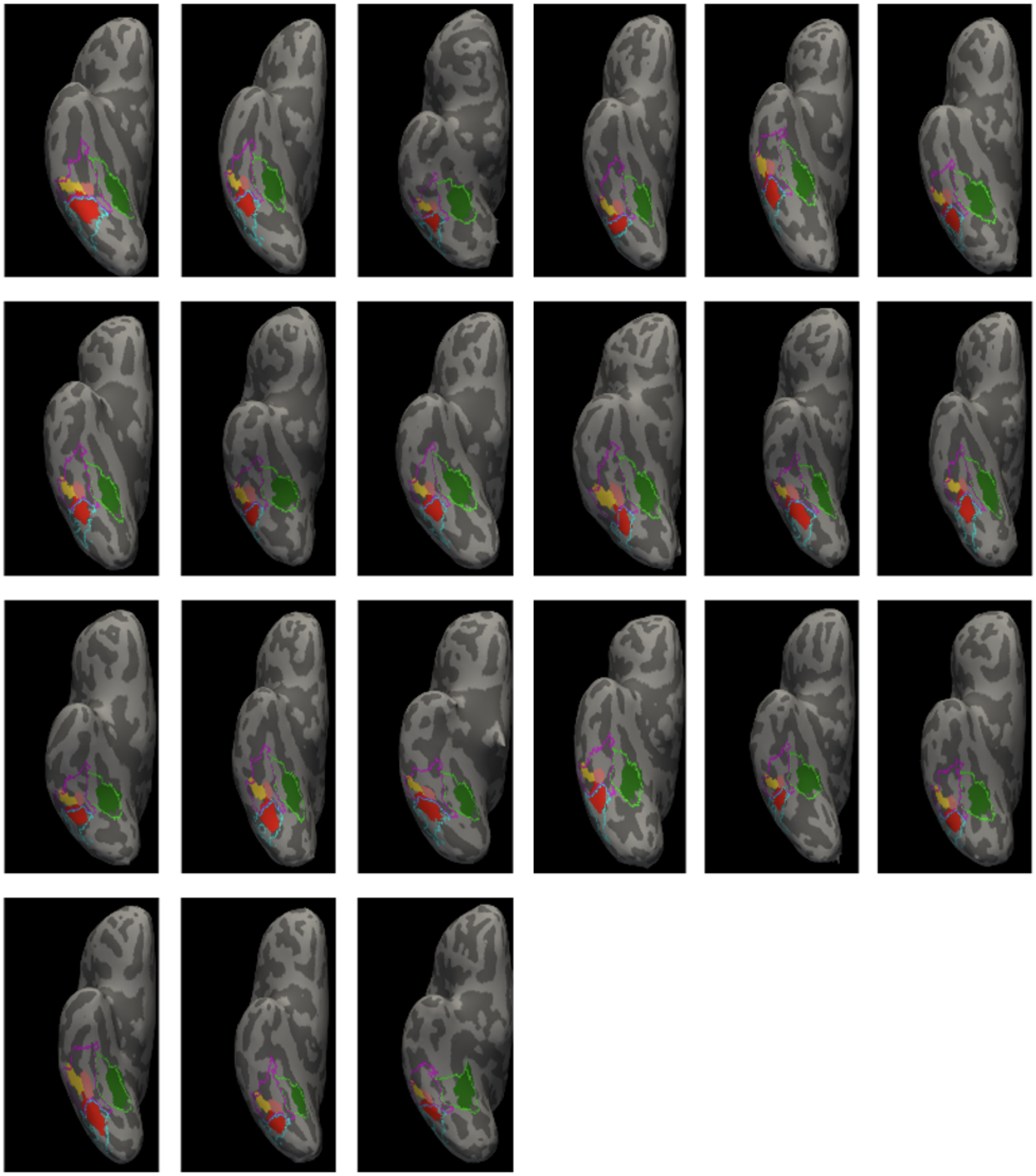

**Supplementary Figure 7.** Right hemisphere MPM fROIs mapped to the brain of each individual infant subject in the 6-month old age group ordered by age. Colors indicate functional ROI: green: CoS-places; pink: mFus-faces, red: pFus-faces; yellow: OTS-bodies; light blue: mOTS-words; blue: pOTS-words. Outlines indicate cytoarchitectonic area: *cyan:* FG2, *green:* FG3, *magenta:* FG4.

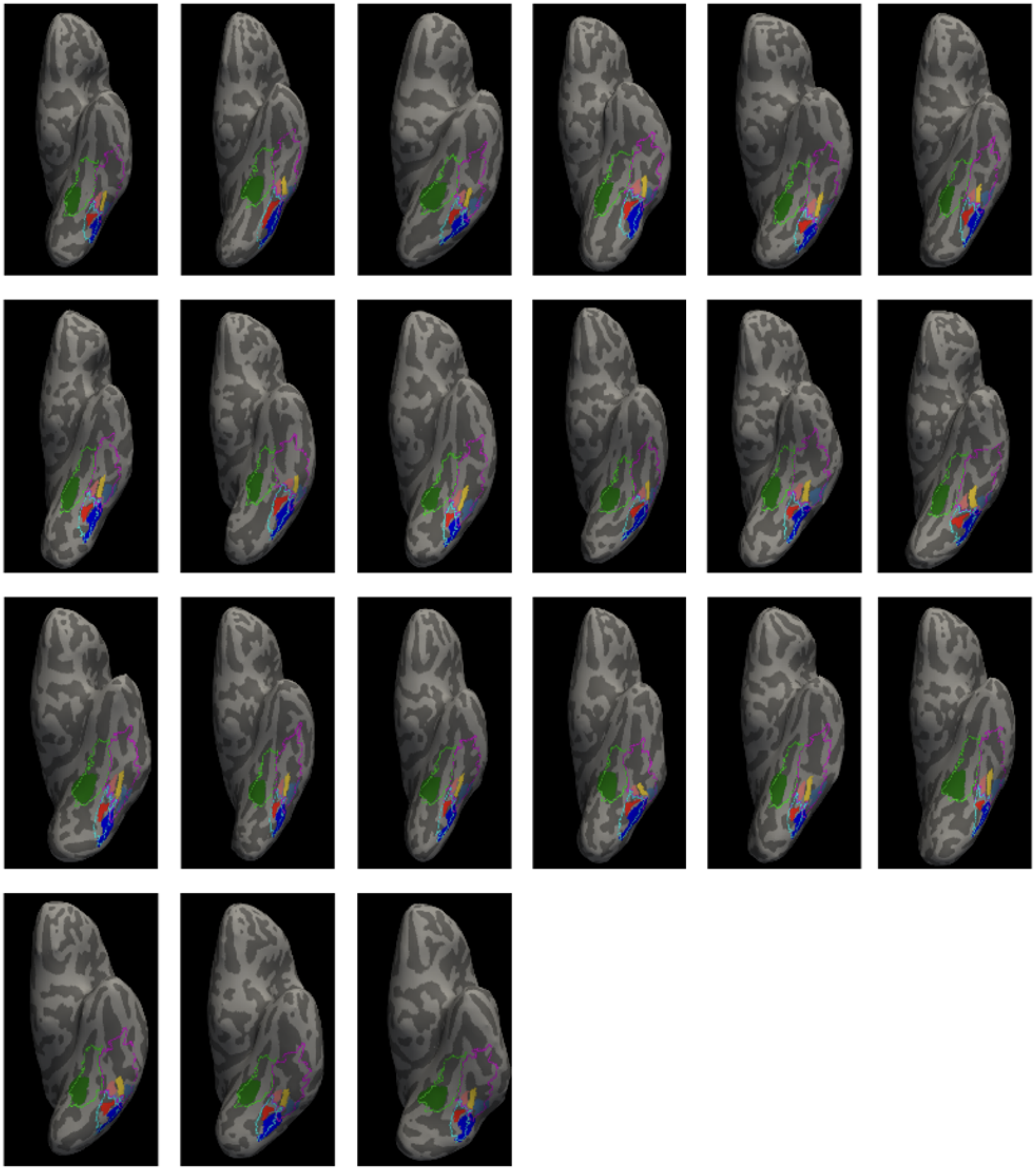

**Supplementary Figure 8.** Left hemisphere MPM ROIs mapped to the brain of each individual subject in the adult age group ordered by age. Colors indicate functional ROI: green: CoS-places; pink: mFus-faces, red: pFus-faces; yellow: OTS-bodies; light blue: mOTS-words; blue: pOTS-words. Outlines indicate cytoarchitectonic area: *cyan:* FG2, *green:* FG3, *magenta:* FG4.

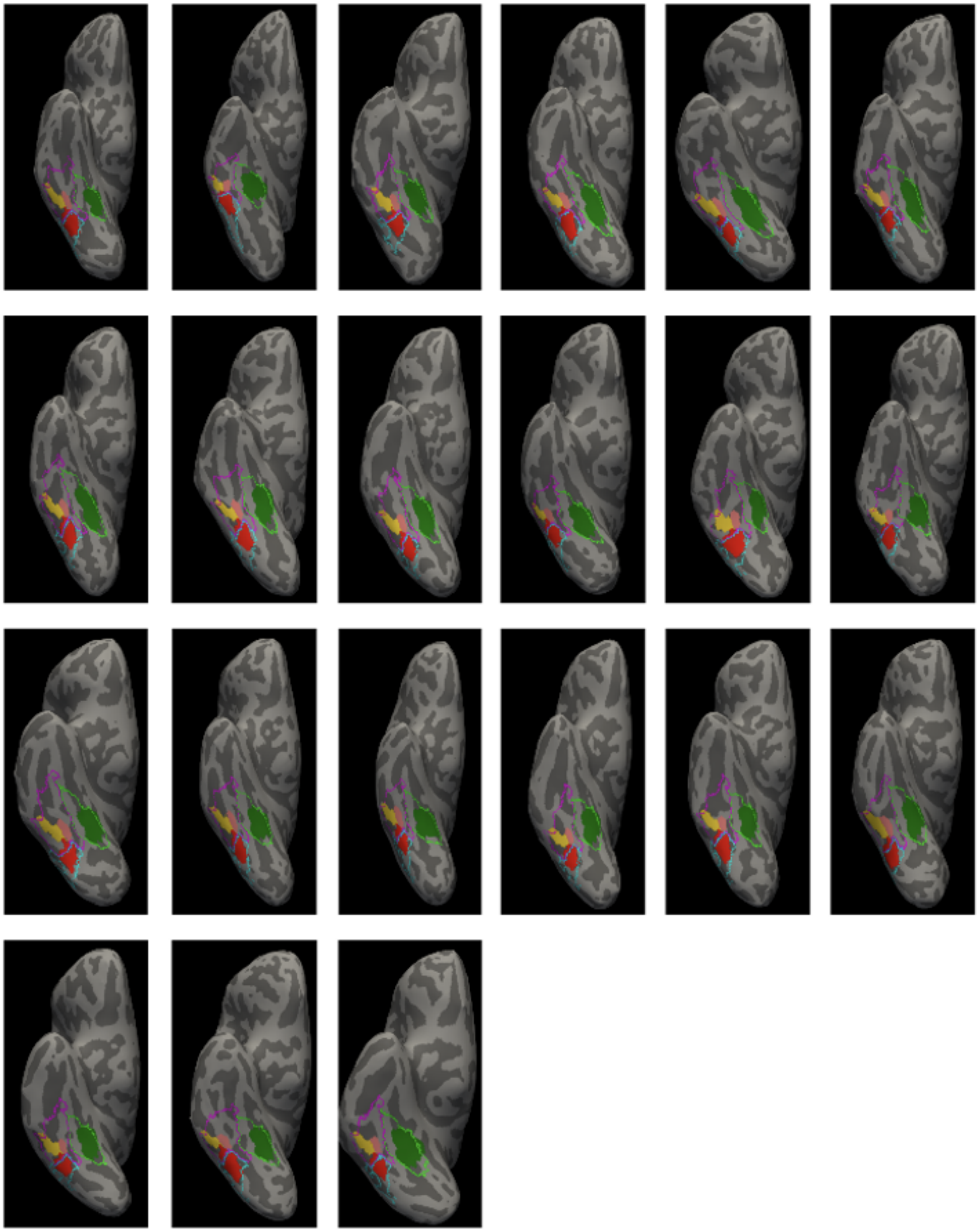

**Supplementary Figure 9.** Right hemisphere MPM fROIs mapped to the brain of each individual infant subject in the adult old age group ordered by age. Colors indicate functional ROI: green: CoS-places; pink: mFus-faces, red: pFus-faces; yellow: OTS-bodies; light blue: mOTS-words; blue: pOTS-words. Outlines indicate cytoarchitectonic area: *cyan:* FG2, *green:* FG3, *magenta:* FG4.

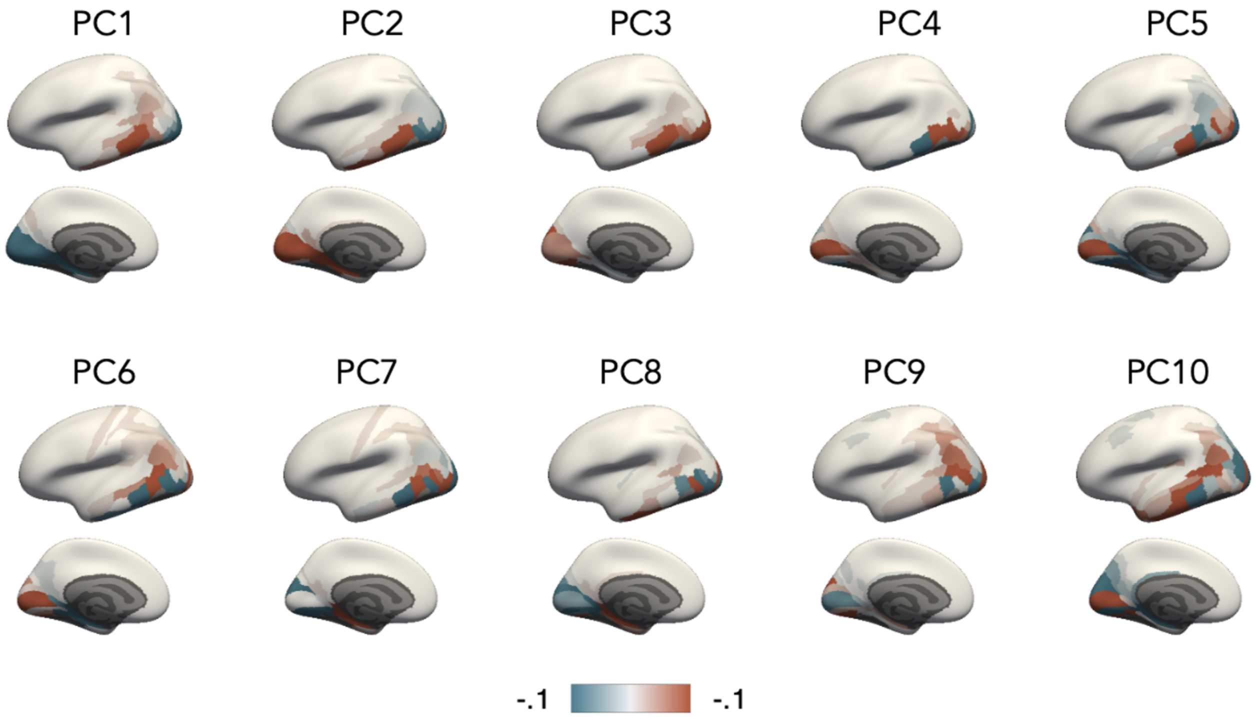


**Supplementary Figure 10.** Loadings of each of the 10 first principal components (PCs) across Glasser ROIs. Colors indicate the coefficient (see color bar).

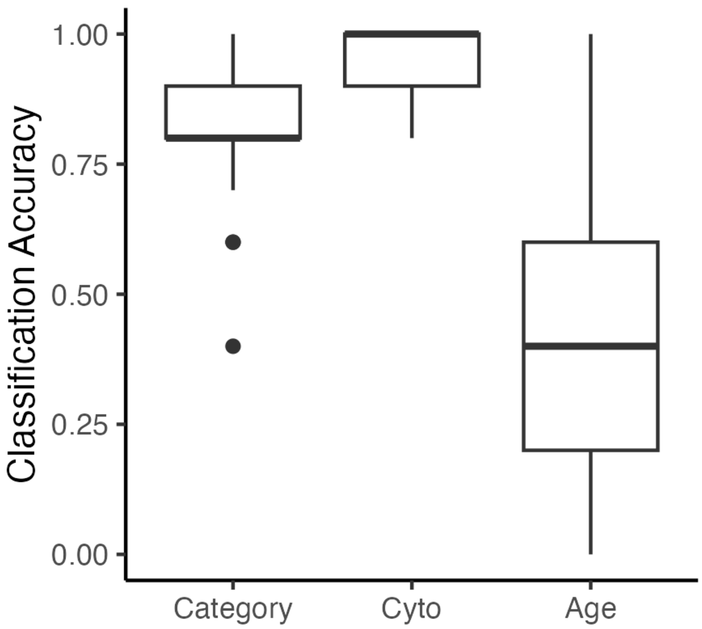


**Supplementary Figure 11.** Boxplots depicting distribution of average leave-one-out classification accuracy of category, cytoarchitecture (Cyto), and age from white matter connectivity profiles (n=88). Center line indicates median, whiskers depict upper and lower quartiles, and points indicate outliers.

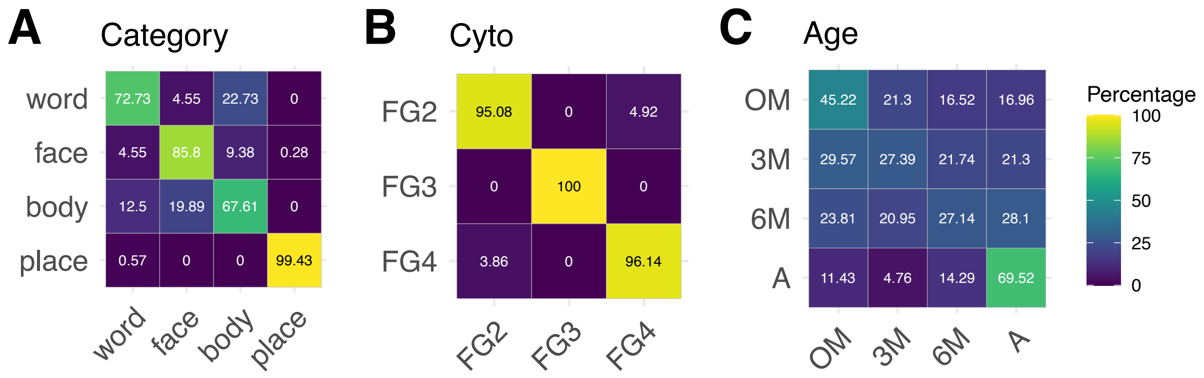


**Supplementary Figure 12.** Confusion matrices for classification of the white matter connectivity profiles of VTC fROIs (n=10) in each participant (n=88) using a balanced training set with 200 examples per each label (sampled with replacement). A) Category classification. B) Cytoarchitecture classification. C) Age classification. All panels: Rows sum to 100. Color depicts the percentage of samples classified within each bin; brighter colors indicate higher percentage of samples (see color bar). On diagonal values are % correct classification and off diagonal values are % incorrect classification.

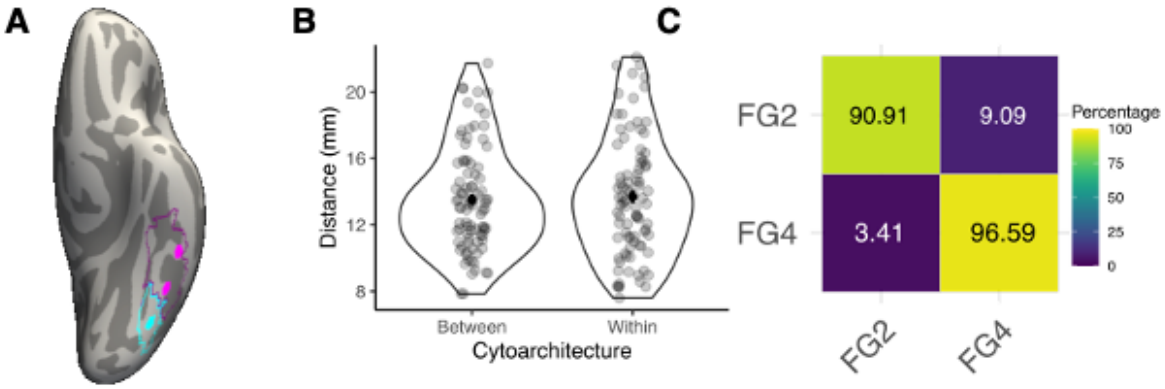


**Supplementary Figure 13.** When holding distance between fROIs constant, cytoarchitecture can be successfully classified from connectivity profiles. A) Disks placed on the cortical surface such that the middle disk is equidistant from two disks, one within the same cytoarchitectonic area (FG4; magenta) and the other in a different cytoarchitectonic area (FG2; cyan). B) Violin plots depicting distance between the ROIs within the same cytoarchitectonic area or between cytoarchitectonic areas, showing no difference in distance ($t(87) = .49$, $p = 0.62$); Black dot and error bar indicate mean± standard error of the mean; transparent dots (n = 88) indicate individual participants C) Confusion matrices depicting leave-one-out cross validation classification accuracy of cytoarchitecture classification in each participant (n=88) in each control ROI (n=3). Rows sum to 100. Color depicts the percentage of samples classified within each bin; brighter colors indicate higher percentage of samples (see color bar). On diagonal values are percentage correct classification and off diagonal values are percentage incorrect classification. Cytoarchitecture is classified successfully and significantly above chance (mean accuracy ± SD=95% ± 13%, $t(87)= 31.62$, $p = 2.2x10^{-16}$).

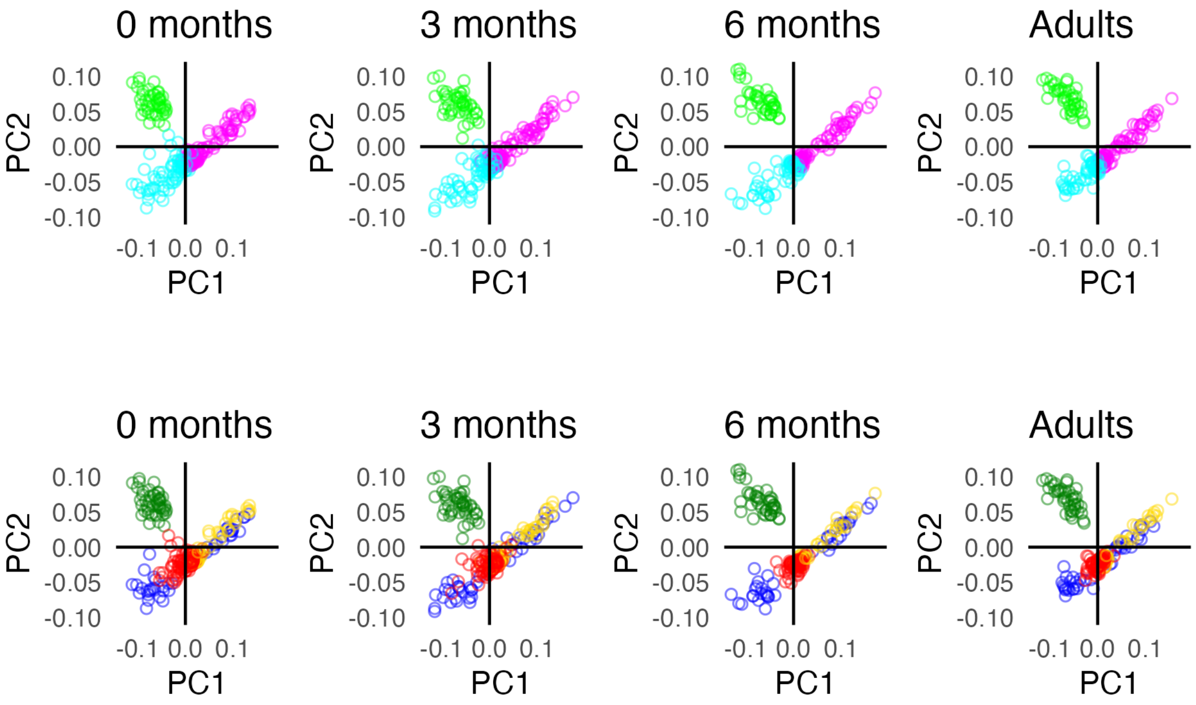


**Supplementary Figure 14.** White matter connectivity profiles of all participants and fROIs projected to the first two principal components (PC1 and PC2). Each dot is a connectivity profile. Each column is an age group. Top row: colored by cytoarchitectonic area: *cyan:* FG2, *green:* FG3, *magenta:* FG4. Bottom row: colored by category: *red*: faces, *green:* places, *blue*: words, *yellow:* bodies. Mean classification accuracy±SD for cytoarchitecture : 0 months: 97%± 16%, 3-months: 97%± 18%, 6-months: 96%±20%, adults: 97%±18%. Mean classification accuracy±SD for category: 0 months: 84%± 37%, 3-months: 82%± 39%, 6-months: 83%±37%, adults: 81%±39%.

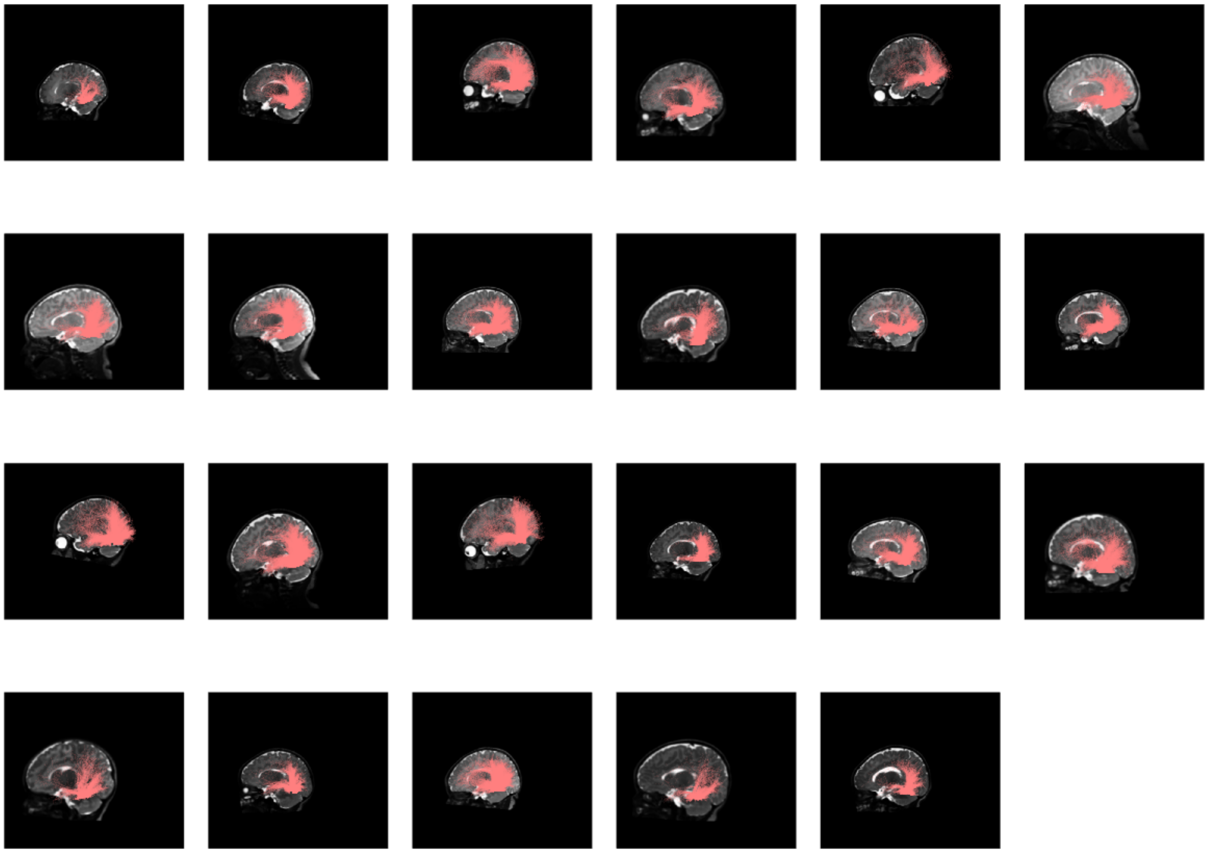

**Supplementary Figure 15.** White matter connections of mFus-faces in the 0-month age group in the left hemisphere. Each panel is a different individual and session. Panels are organized by age from top left to bottom right, with the youngest in the top left.

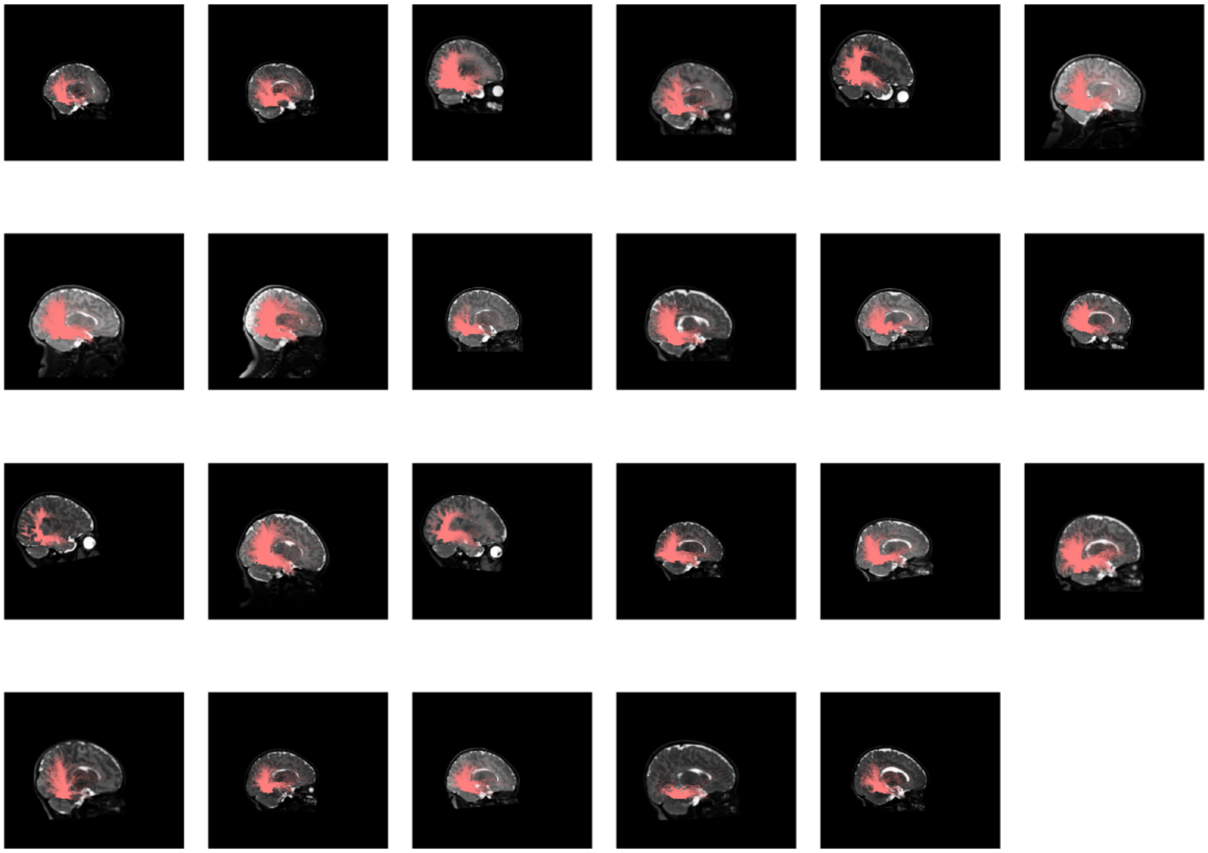

**Supplementary Figure 16.** White matter connections of mFus-faces in the 0-month age group organized by age in the right hemisphere.

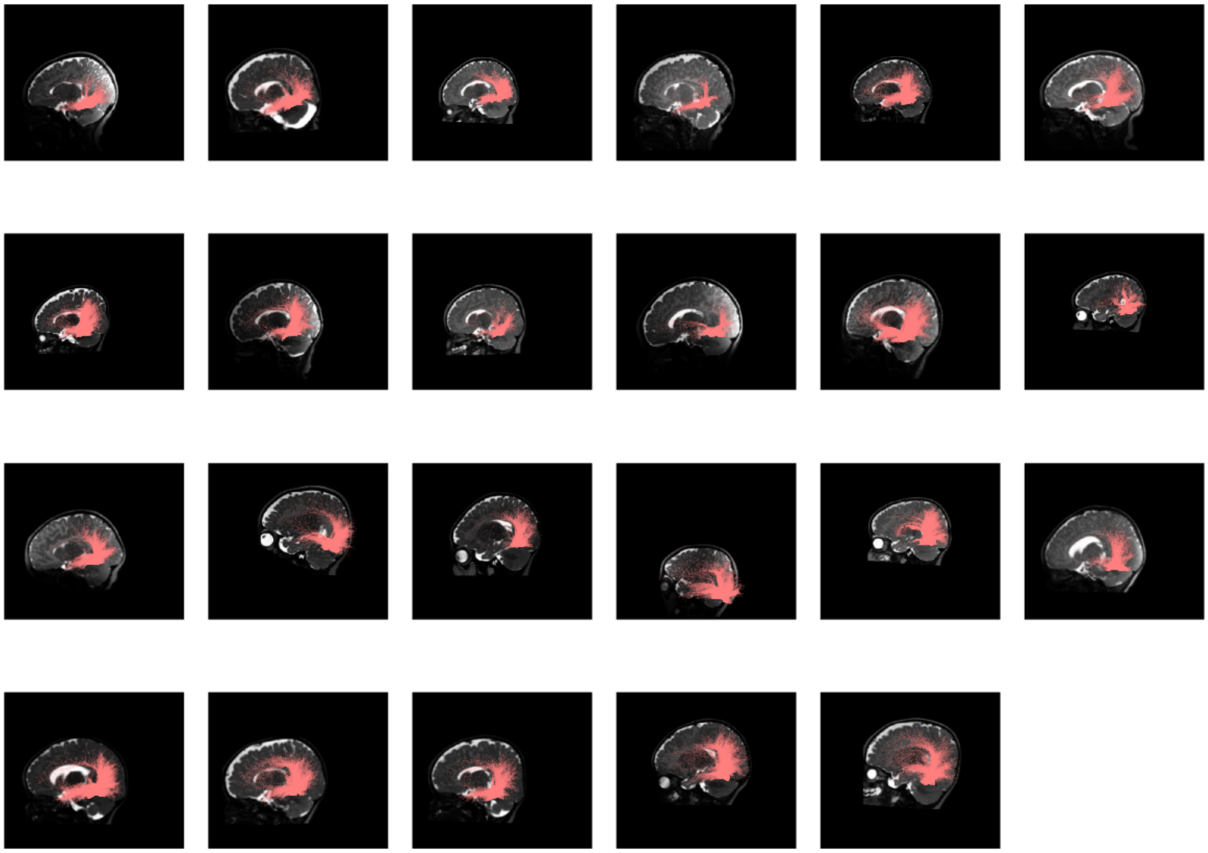

**Supplementary Figure 17.** White matter connections of mFus-faces in the 3-month age group organized by age in the left hemisphere.

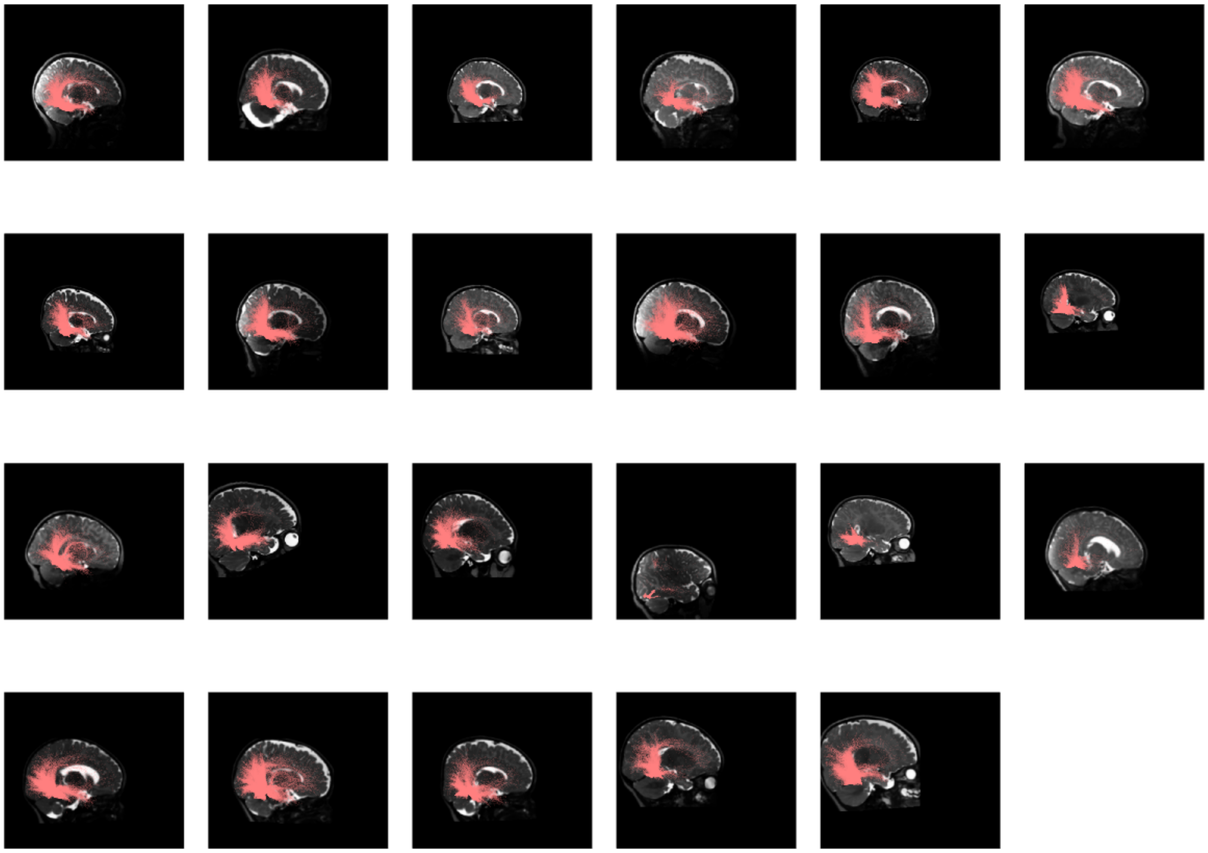

**Supplementary Figure 18.** White matter connections of mFus-faces in the 3-month age group organized by age in the right hemisphere.

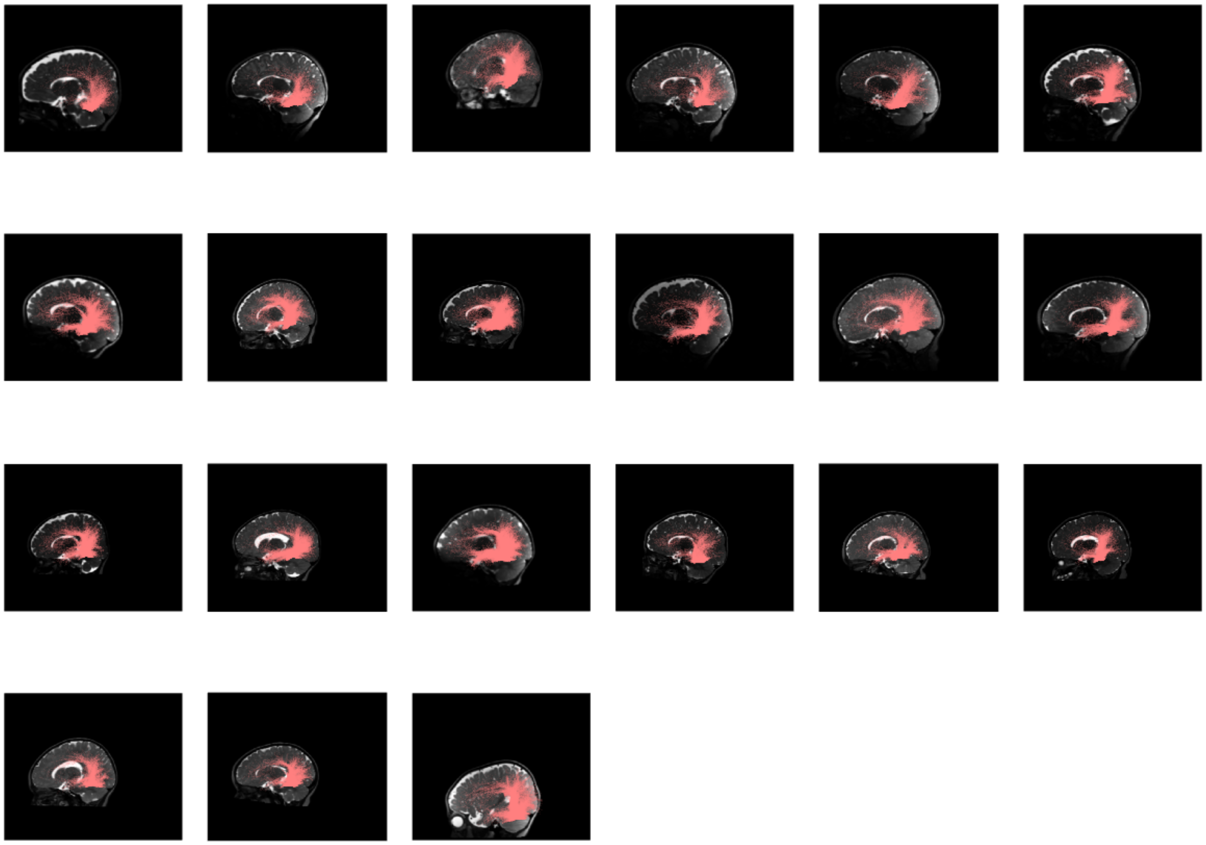

**Supplementary Figure 19.** White matter connections of mFus-faces in the 6-month age group organized by age in the left hemisphere.

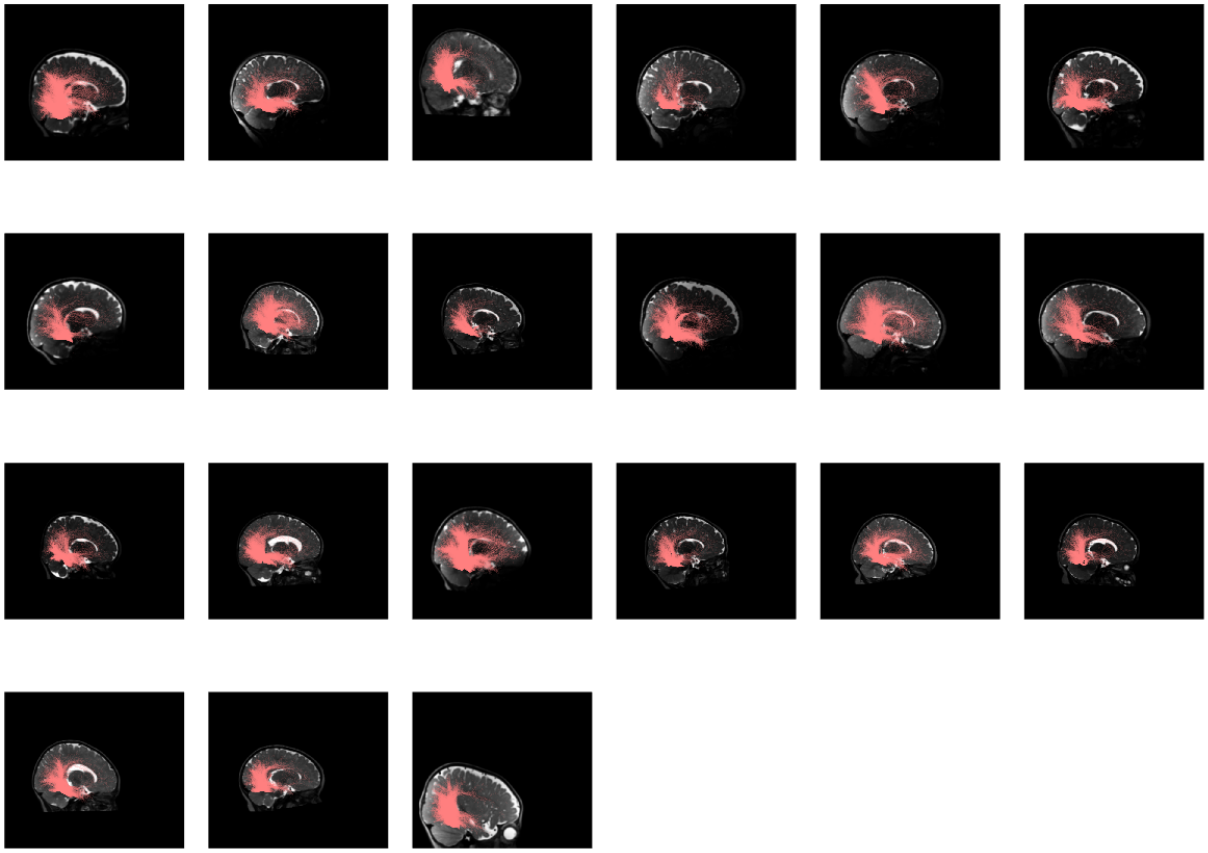

**Supplementary Figure 20.** White matter connections of mFus-faces in the 6-month age group organized by age in the right hemisphere.

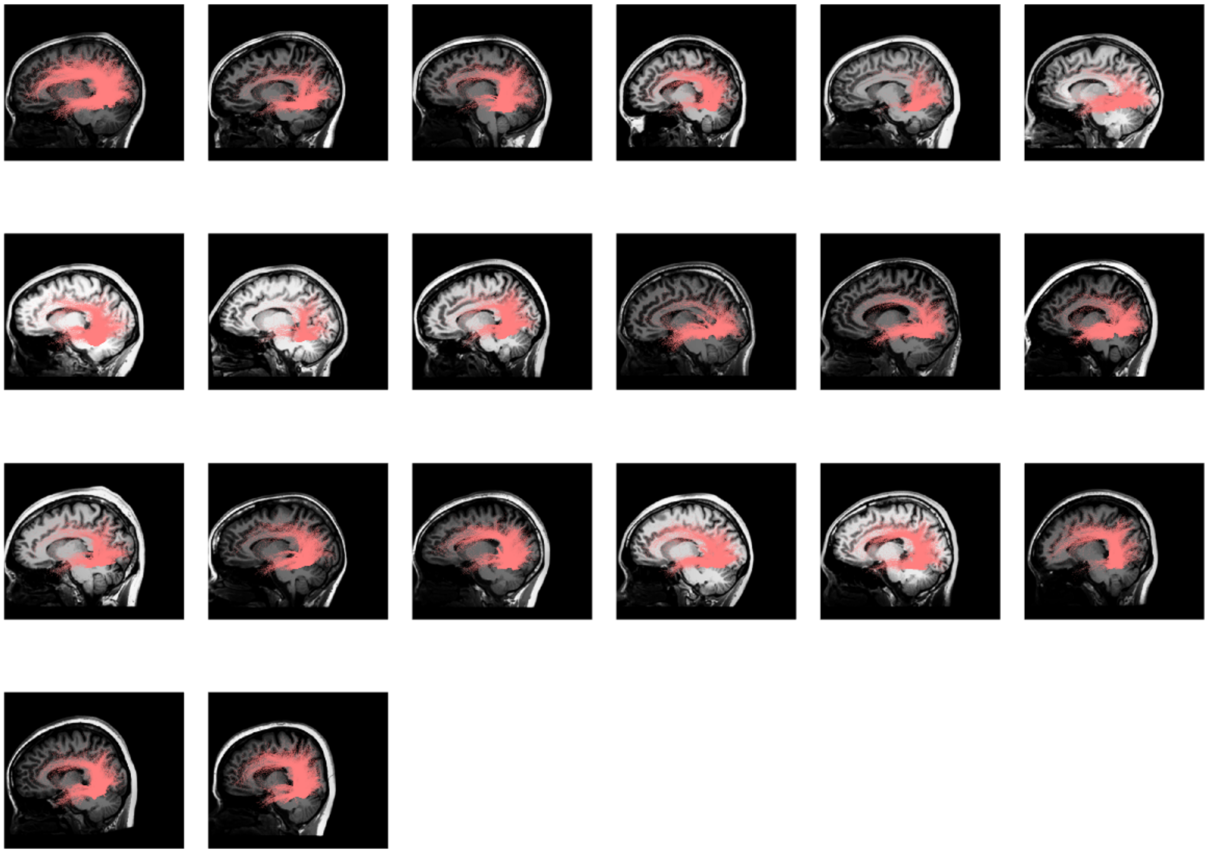

**Supplementary Figure 21.** White matter connections of mFus-faces in the adult age group organized by age in the left hemisphere.

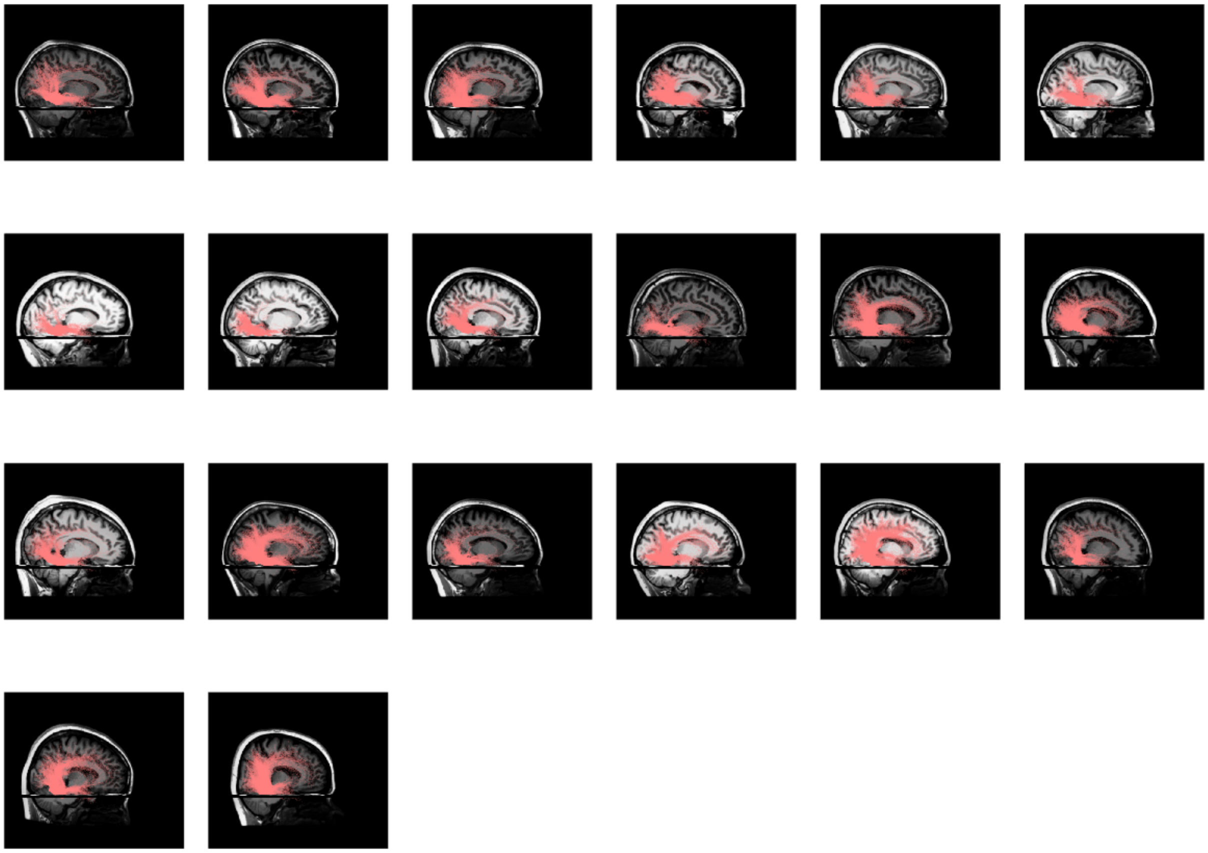

**Supplementary Figure 22.** White matter connections of mFus-faces in the adult age group organized by age in the right hemisphere.

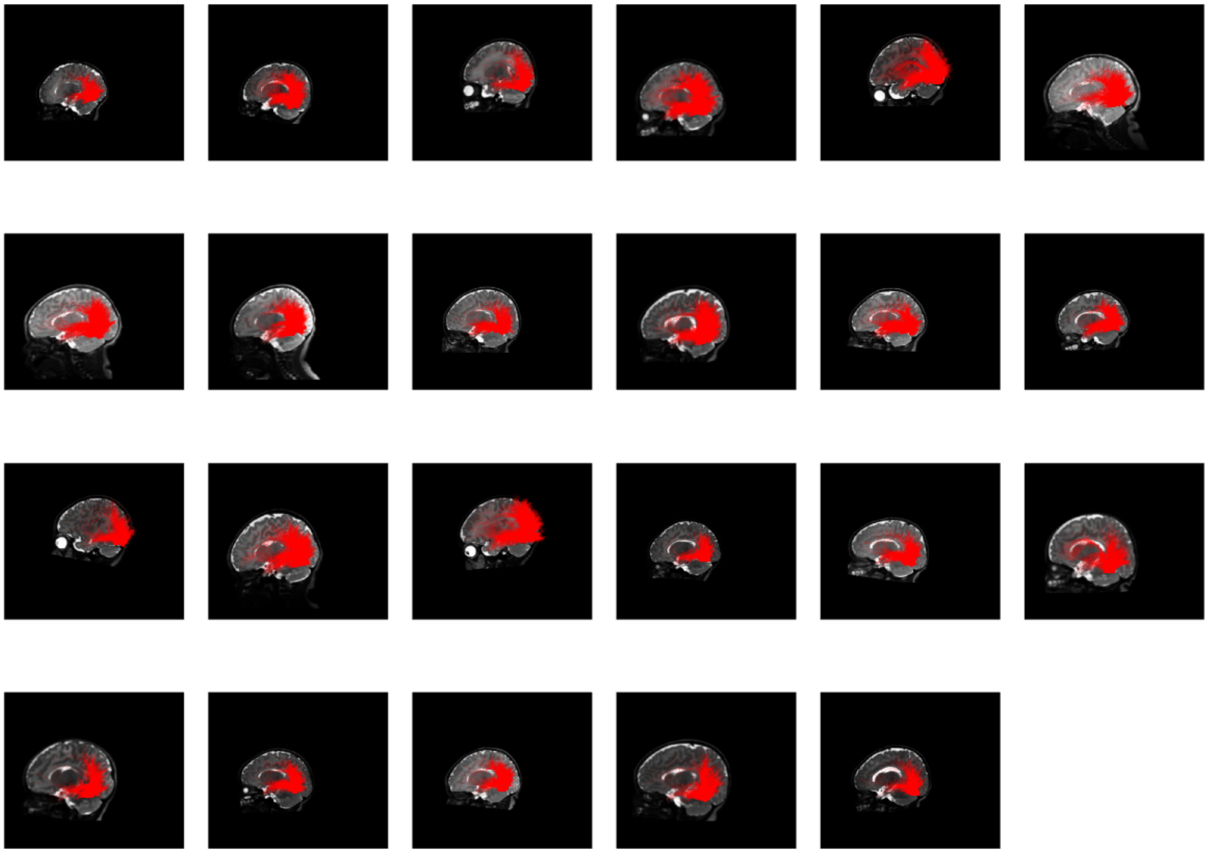

**Supplementary Figure 23.** White matter connections of pFus-faces in the 0-month age group organized by age in the left hemisphere.

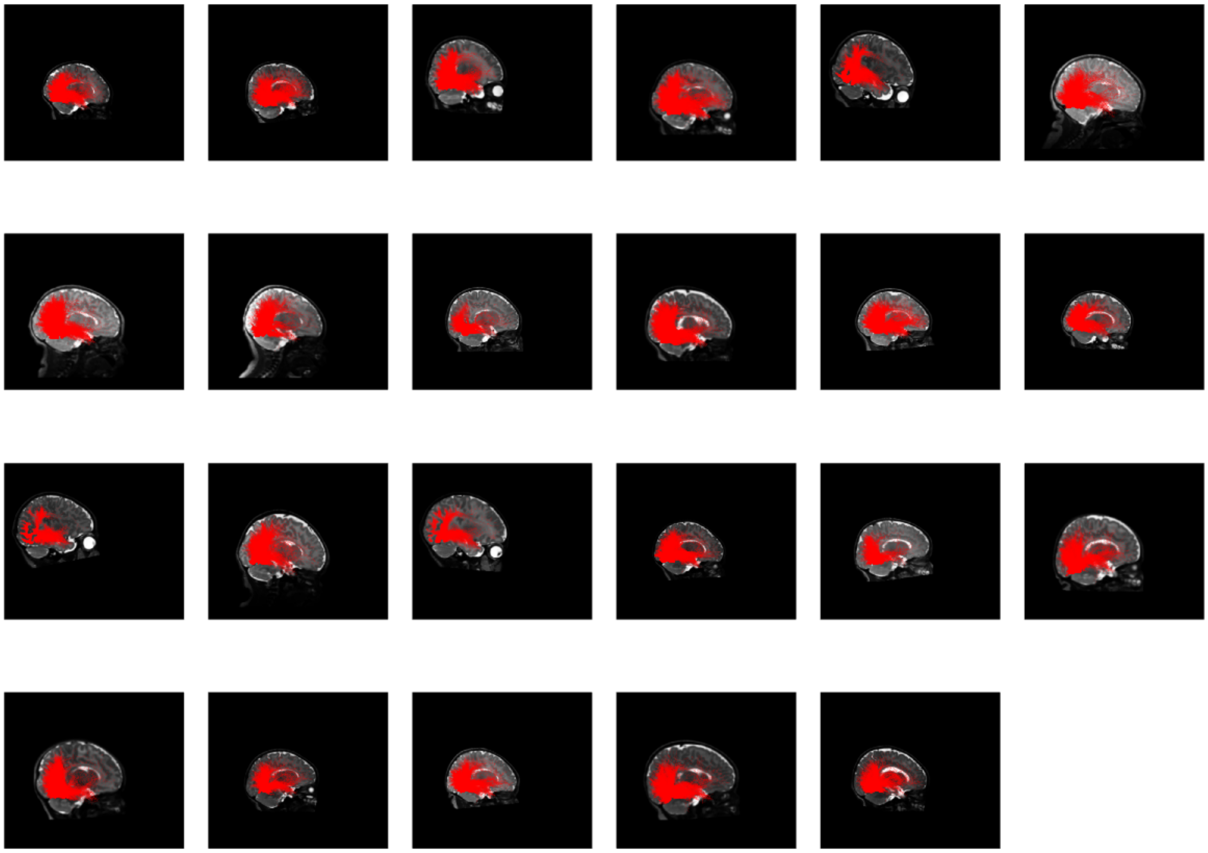

**Supplementary Figure 24.** White matter connections of pFus-faces in the 0-month age group organized by age in the right hemisphere.

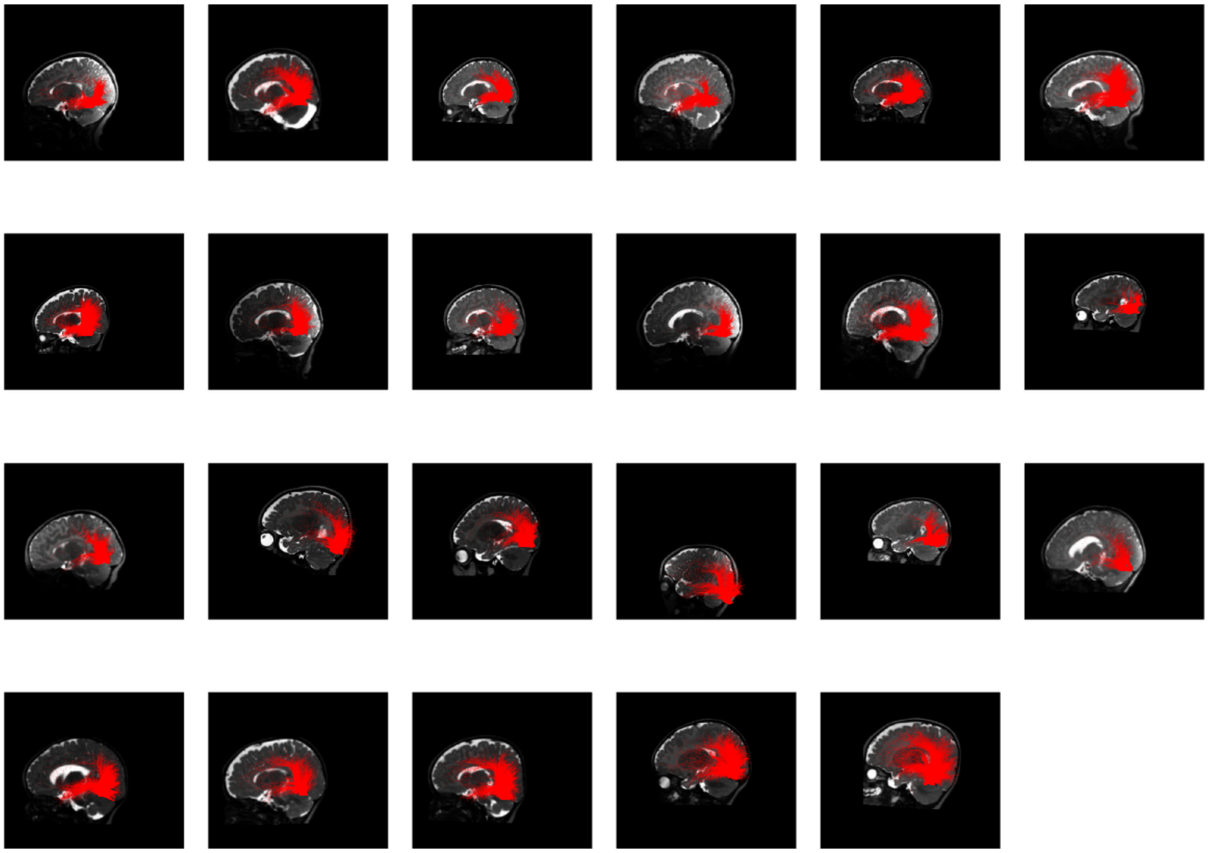

**Supplementary Figure 25.** White matter connections of pFus-faces in the 3-month age group organized by age in the left hemisphere.

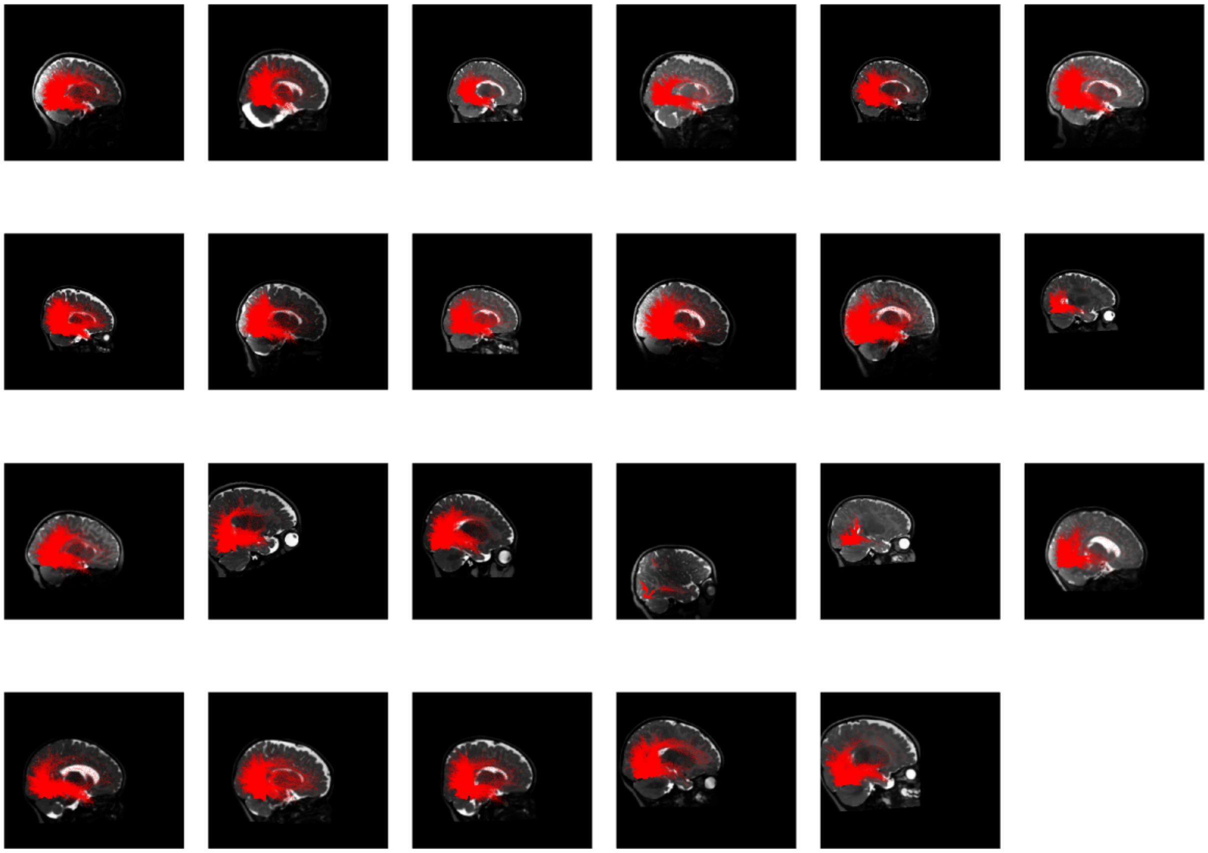

**Supplementary Figure 26.** White matter connections of pFus-faces in the 3-month age group organized by age in the right hemisphere.

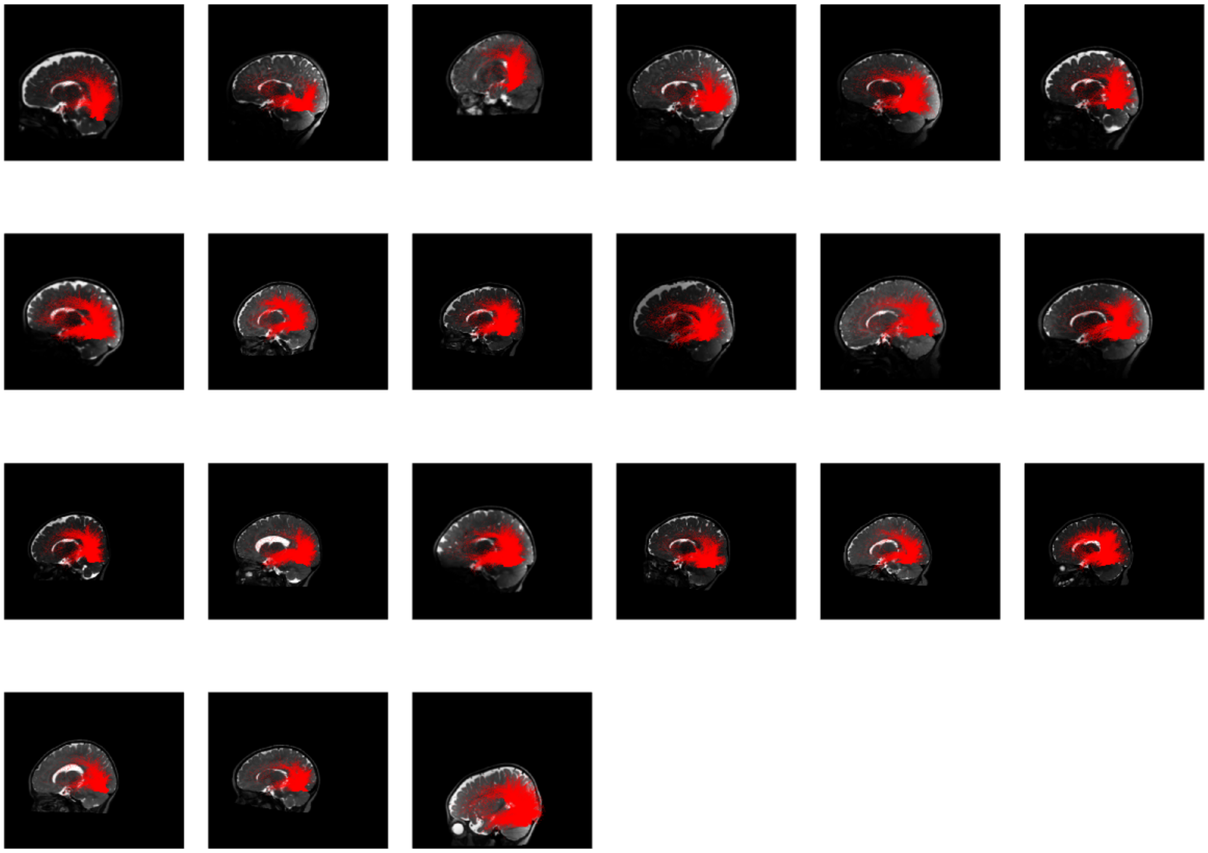

**Supplementary Figure 27.** White matter connections of pFus-faces in the 6-month age group organized by age in the left hemisphere.

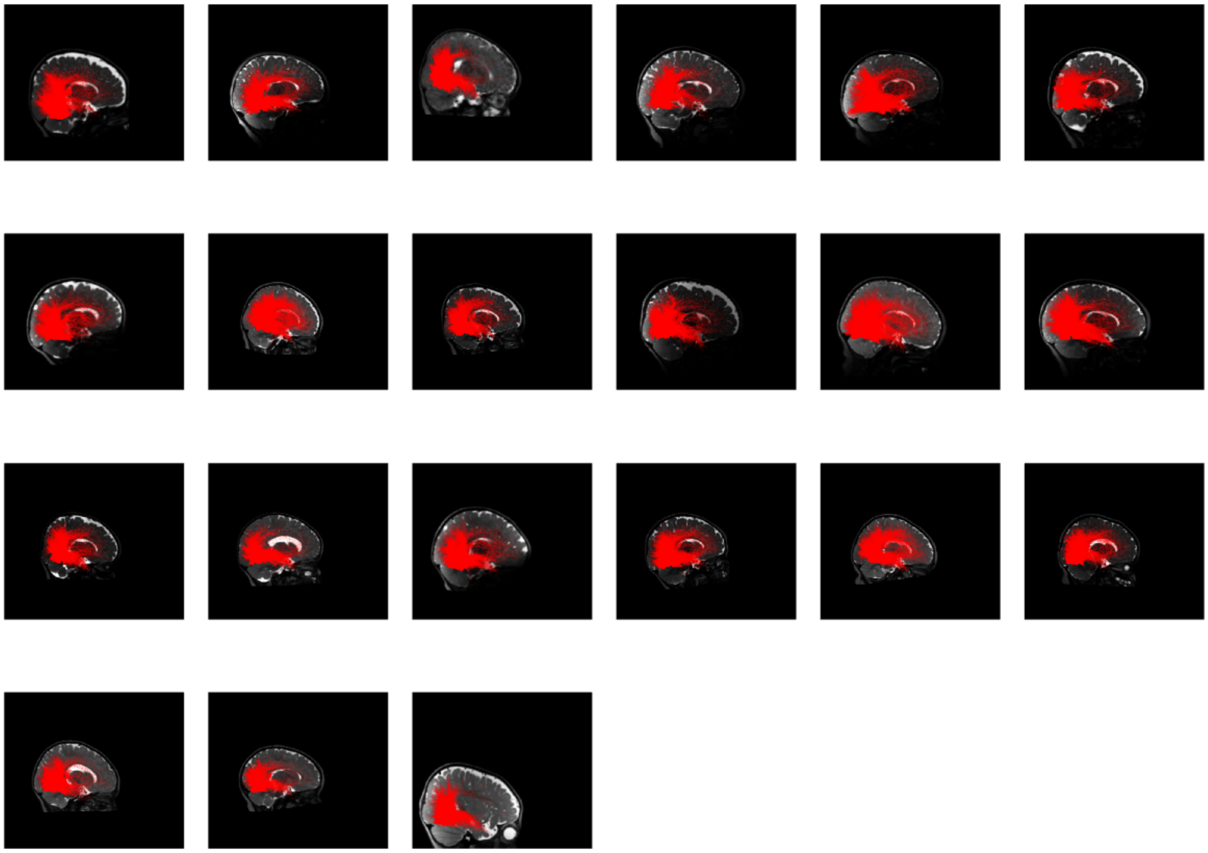

**Supplementary Figure 28.** White matter connections of pFus-faces in the 6-month age group organized by age in the right hemisphere.

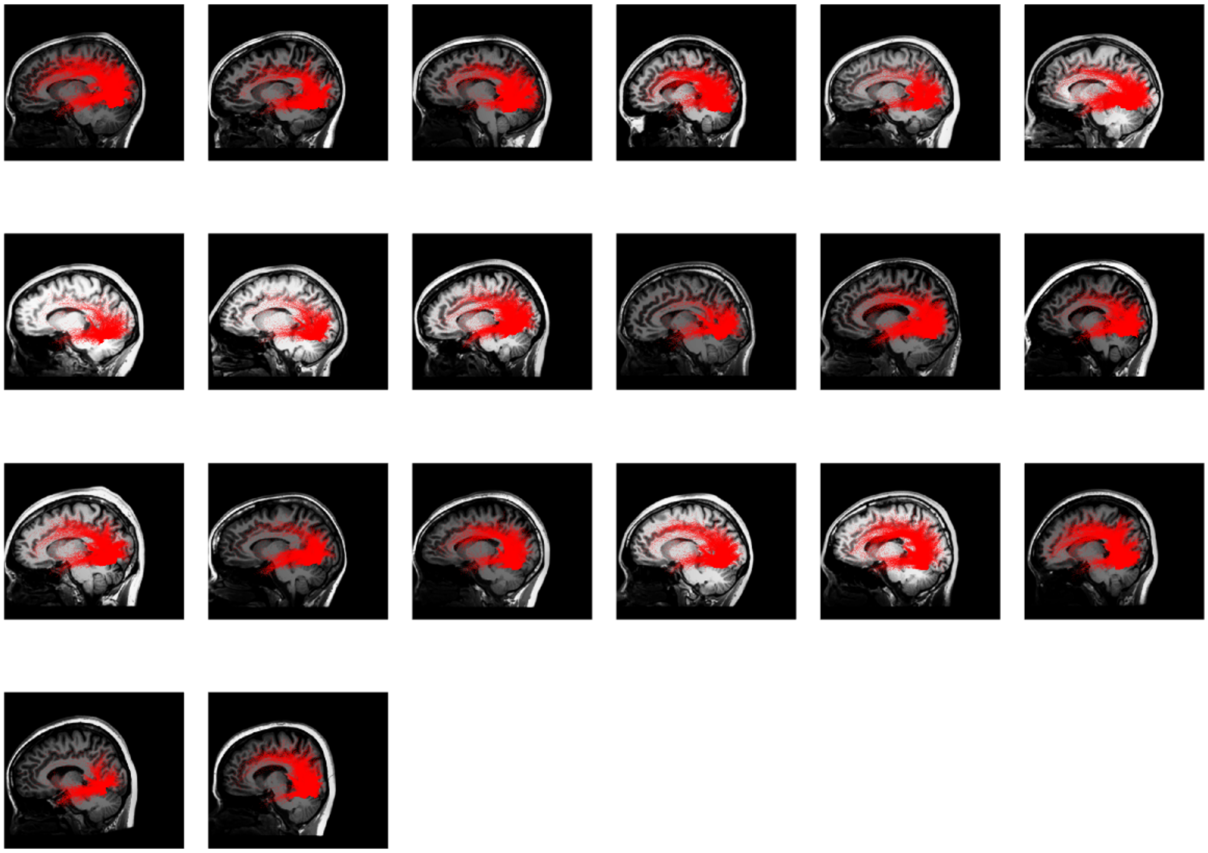

**Supplementary Figure 29.** White matter connections of pFus-faces in the adult age group organized by age in the left hemisphere.

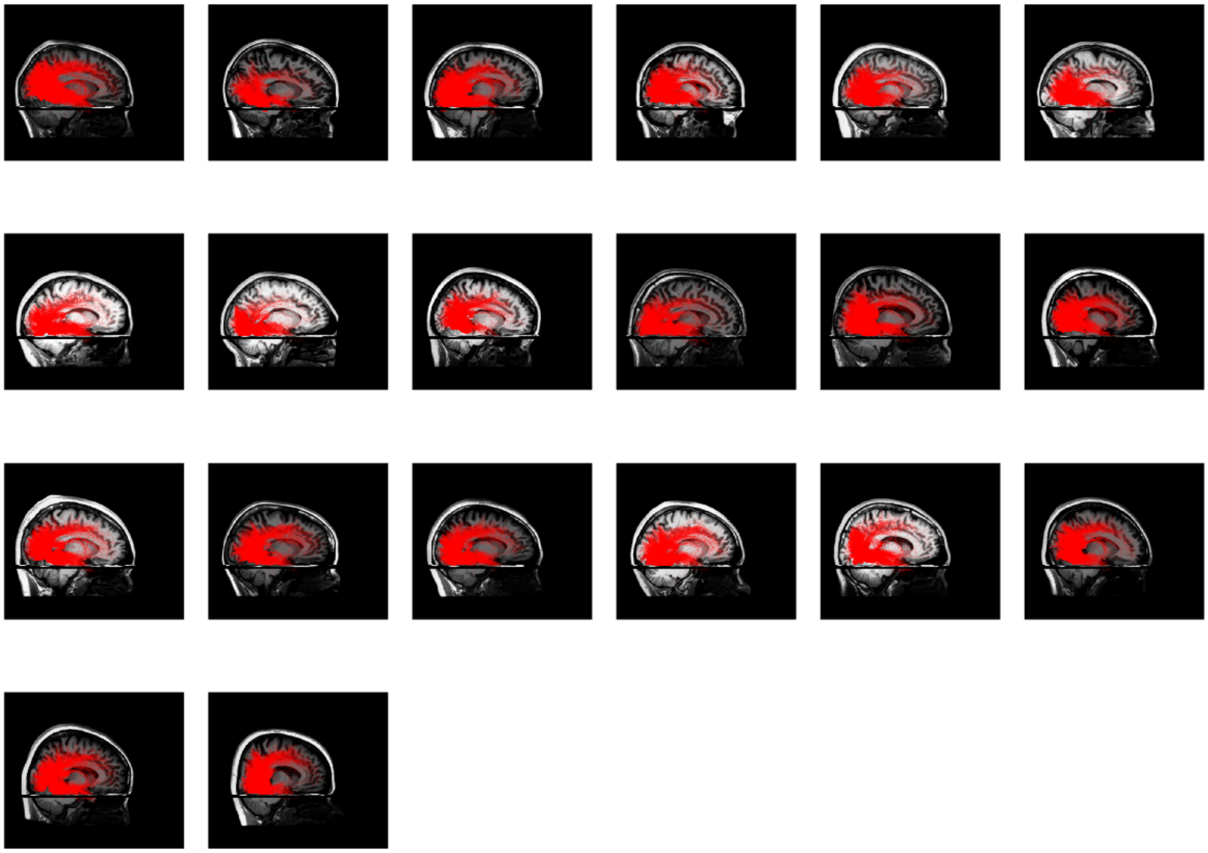

**Supplementary Figure 30.** White matter connections of pFus-faces in the adult age group organized by age in the right hemisphere.

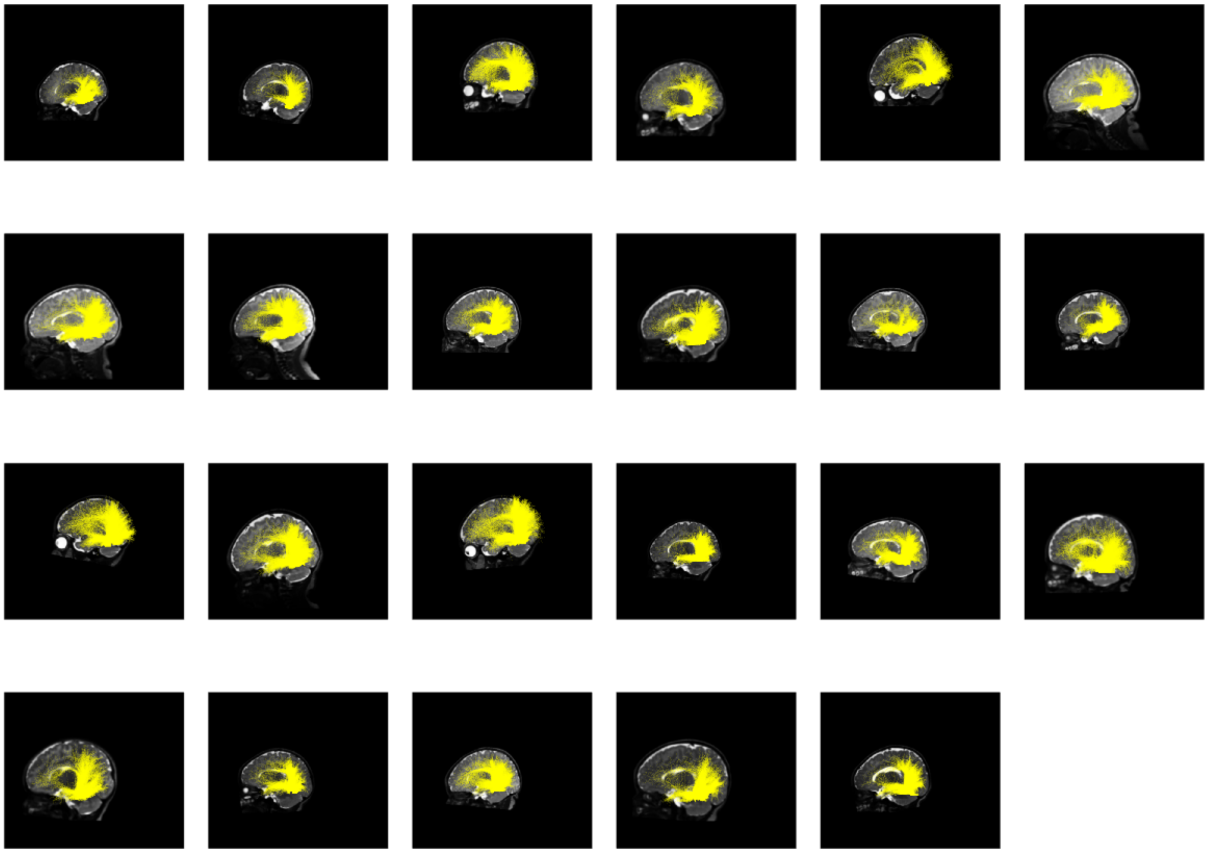

**Supplementary Figure 31.** White matter connections of OTS-bodies words in the 0-month age group organized by age in the left hemisphere.

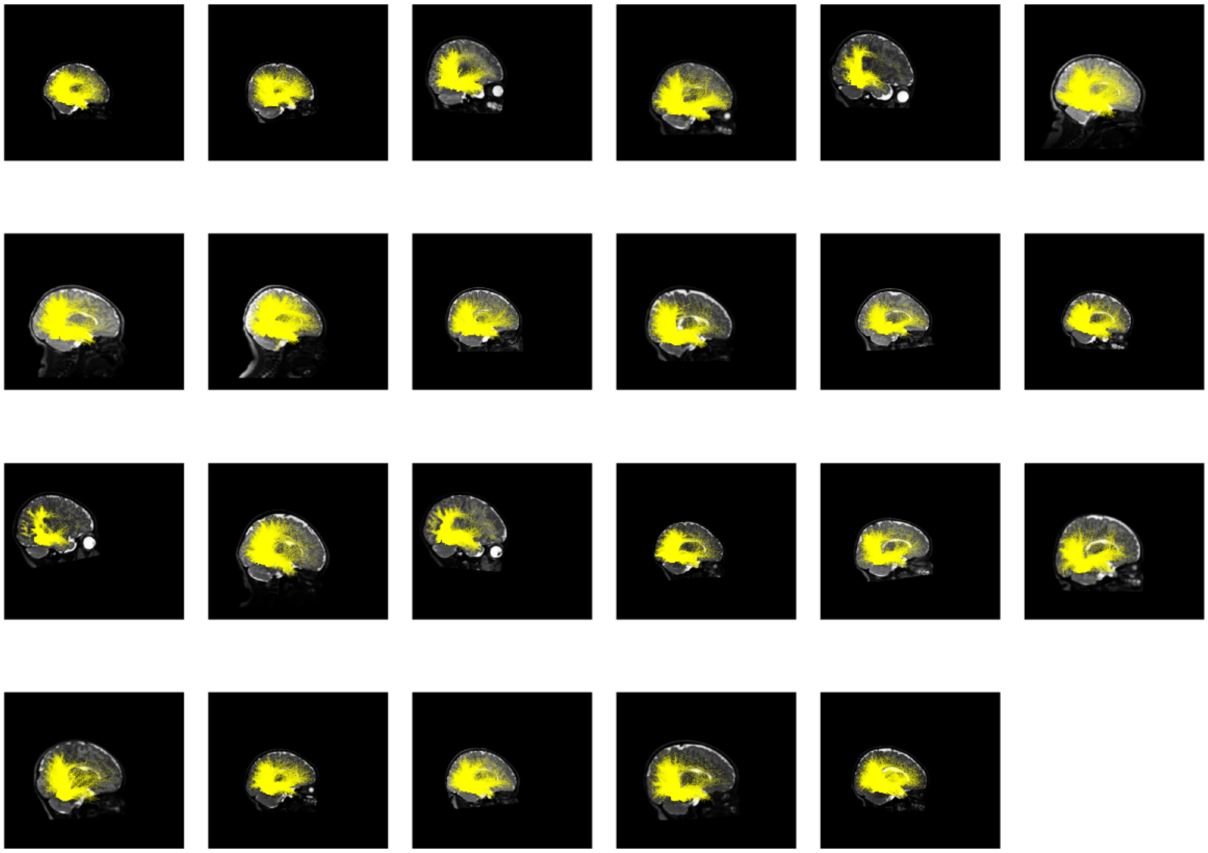

**Supplementary Figure 32.** White matter connections of OTS-bodies words in the 0-month age group organized by age in the right hemisphere.

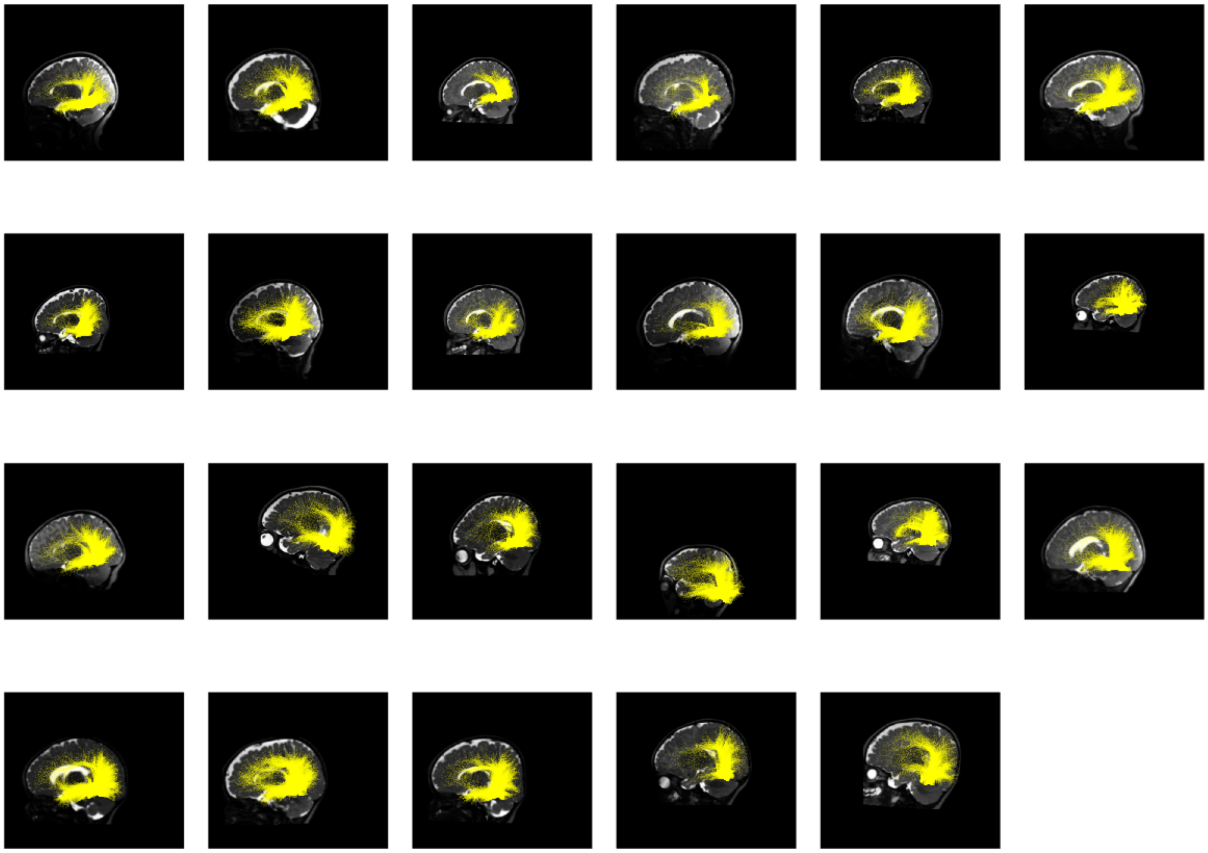

**Supplementary Figure 33.** White matter connections of OTS-bodies words in the 3-month age group organized by age in the left hemisphere.

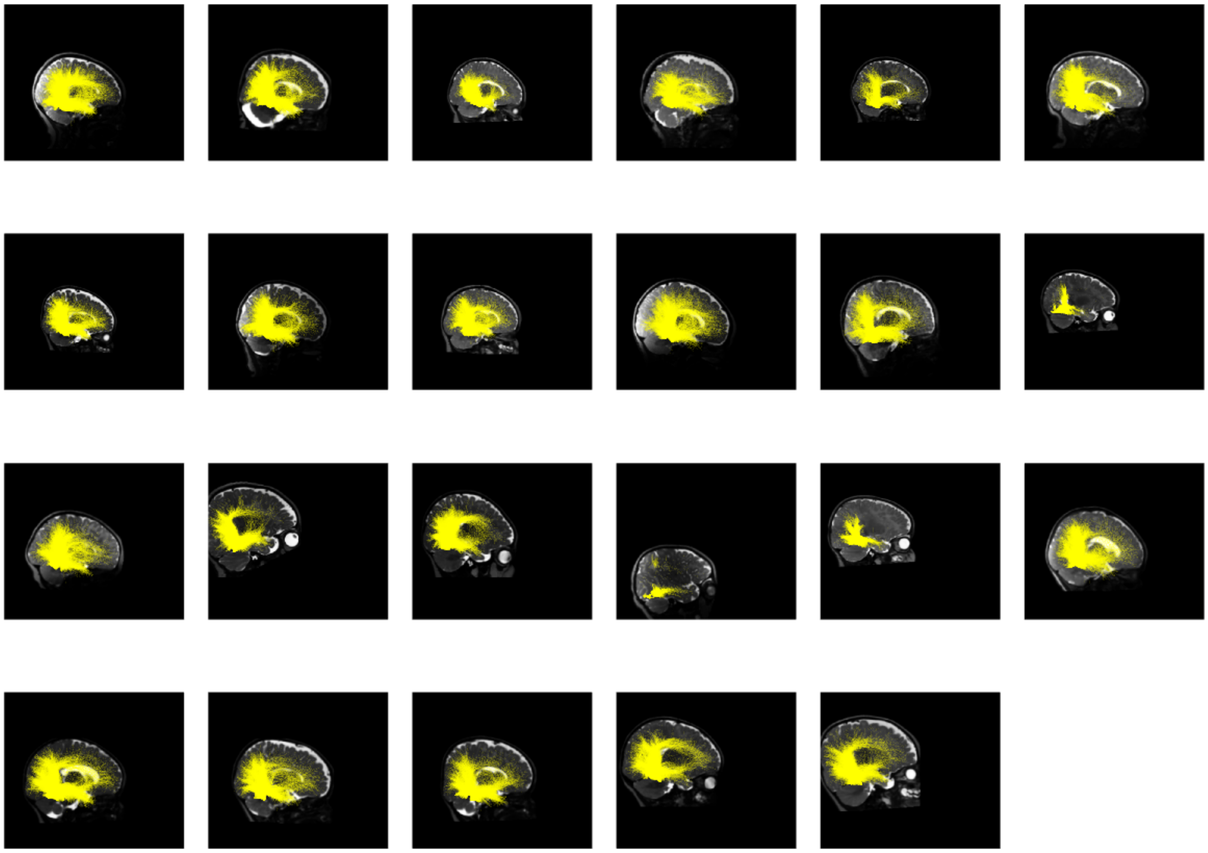

**Supplementary Figure 34.** White matter connections of OTS-bodies words in the 3-month age group organized by age in the right hemisphere.

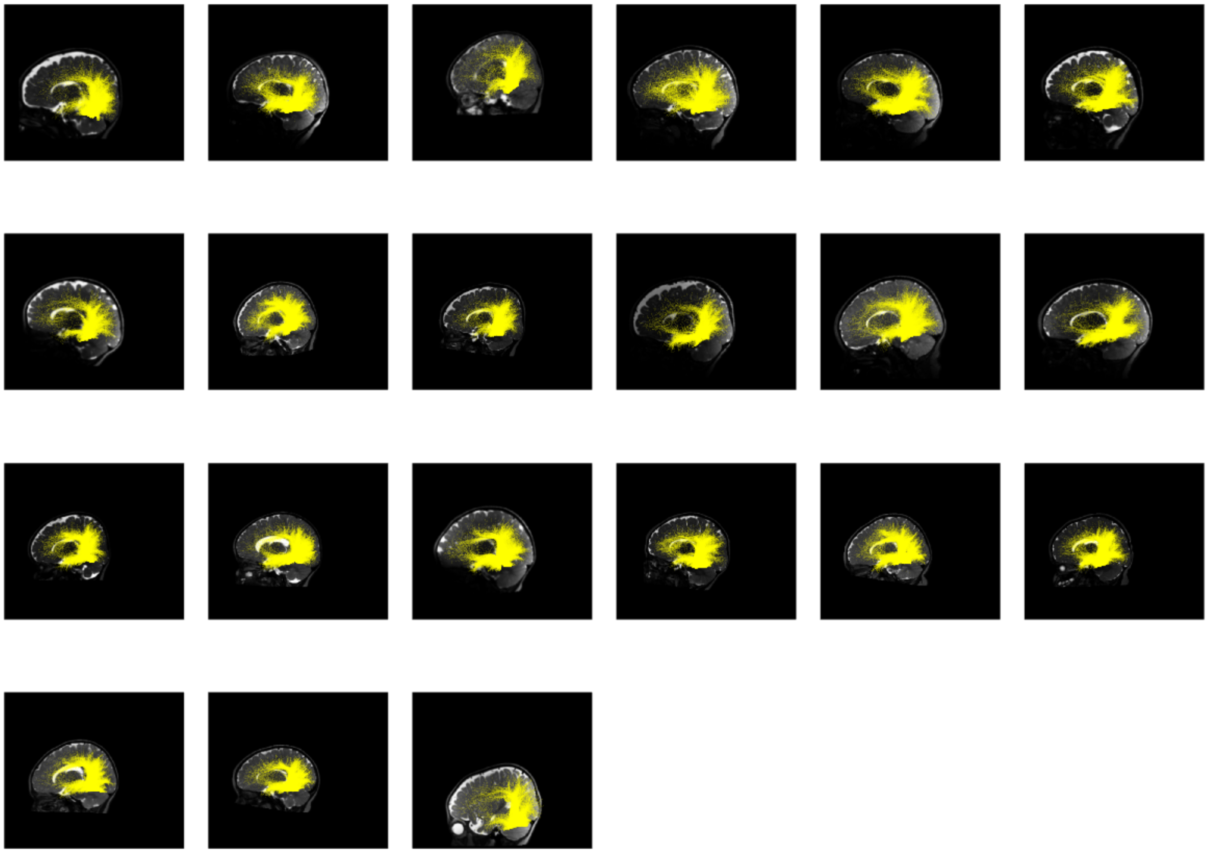

**Supplementary Figure 35.** White matter connections of OTS-bodies words in the 6-month age group organized by age in the left hemisphere.

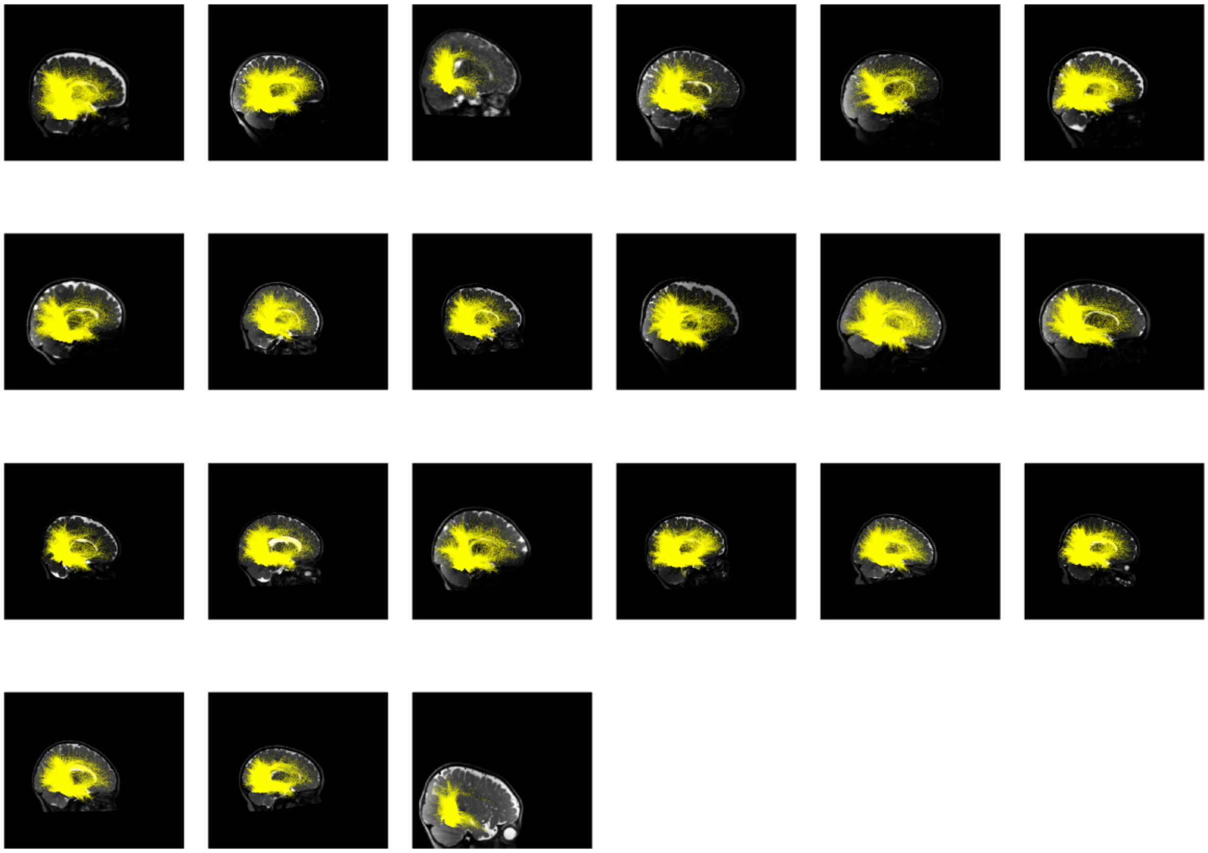

**Supplementary Figure 36.** White matter connections of OTS-bodies words in the 6-month age group organized by age in the right hemisphere.

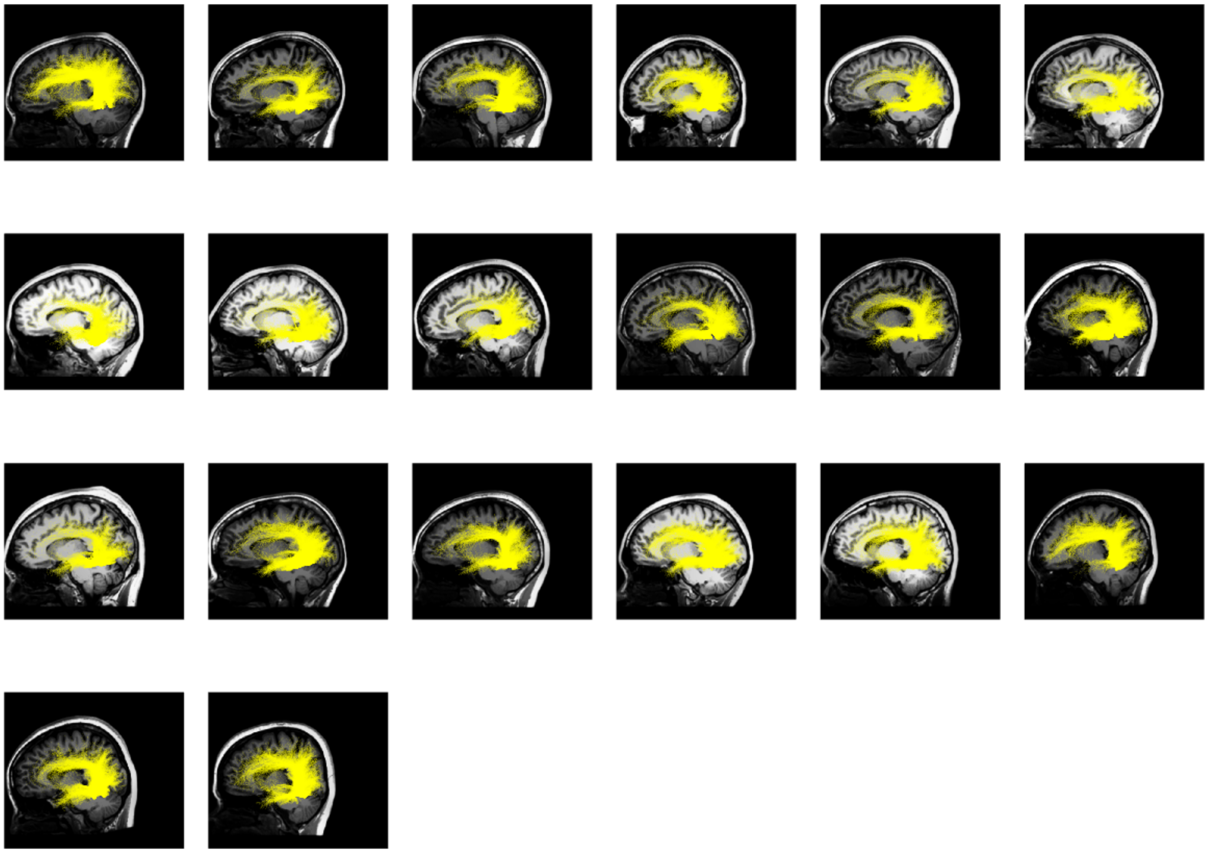

**Supplementary Figure 37.** White matter connections of OTS-bodies in the adult age group organized by age in the left hemisphere.

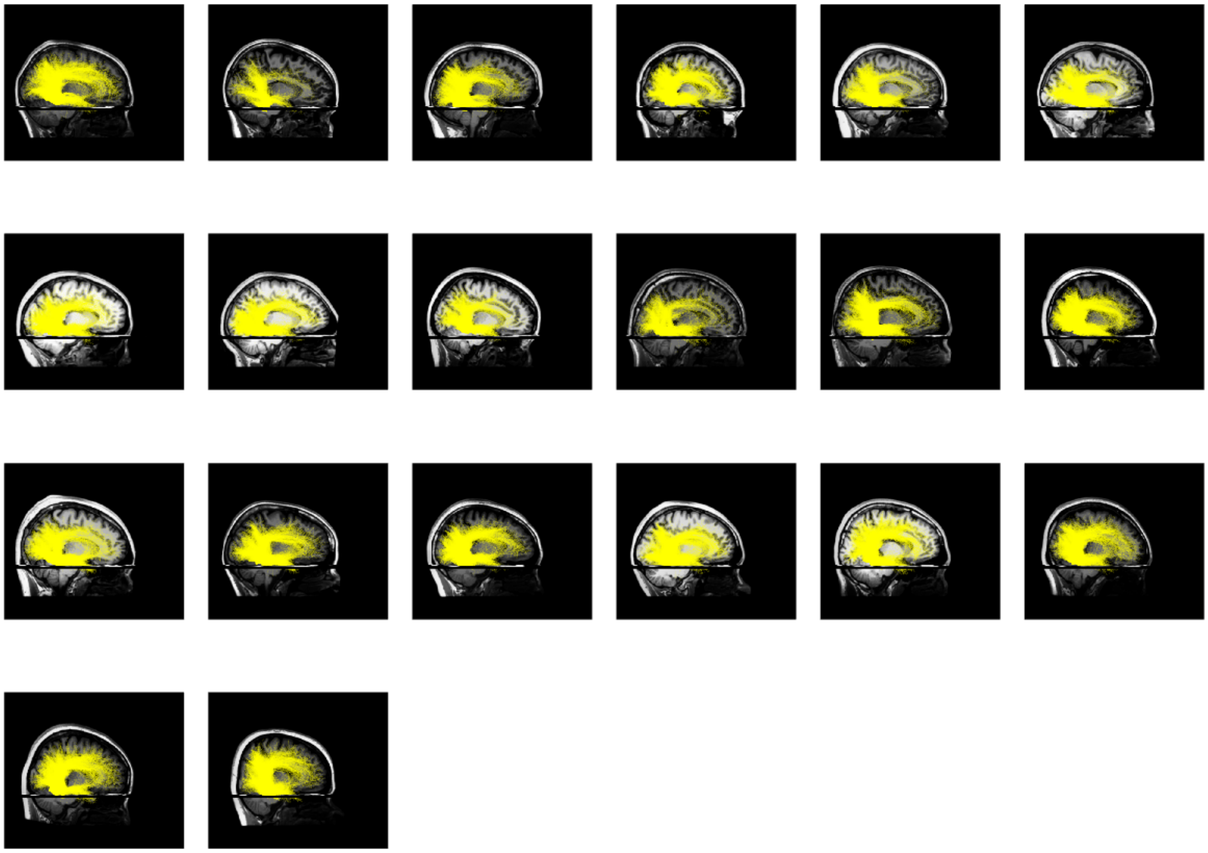

**Supplementary Figure 38.** White matter connections of OTS-bodies in the adult age group organized by age in the right hemisphere.

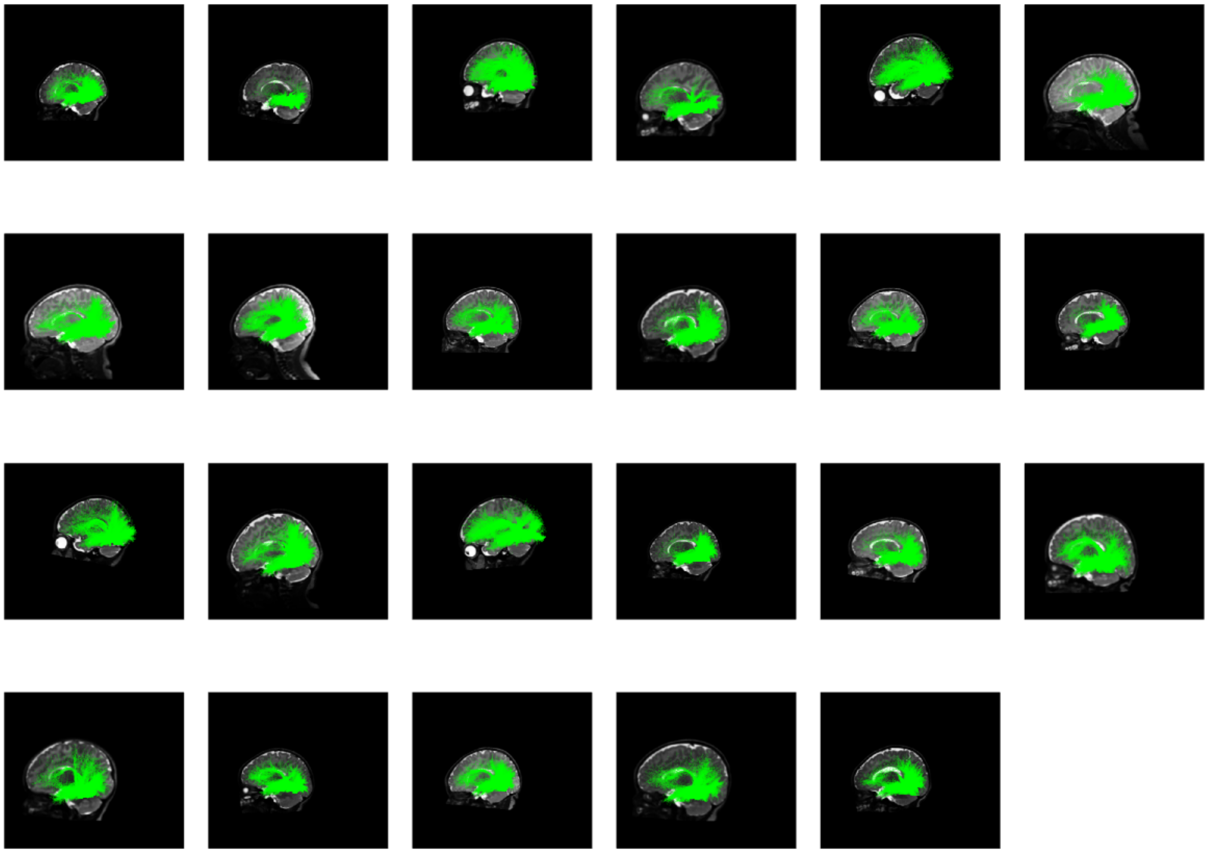

**Supplementary Figure 39.** White matter connections of CoS-places in the 0-month age group organized by age in the left hemisphere.

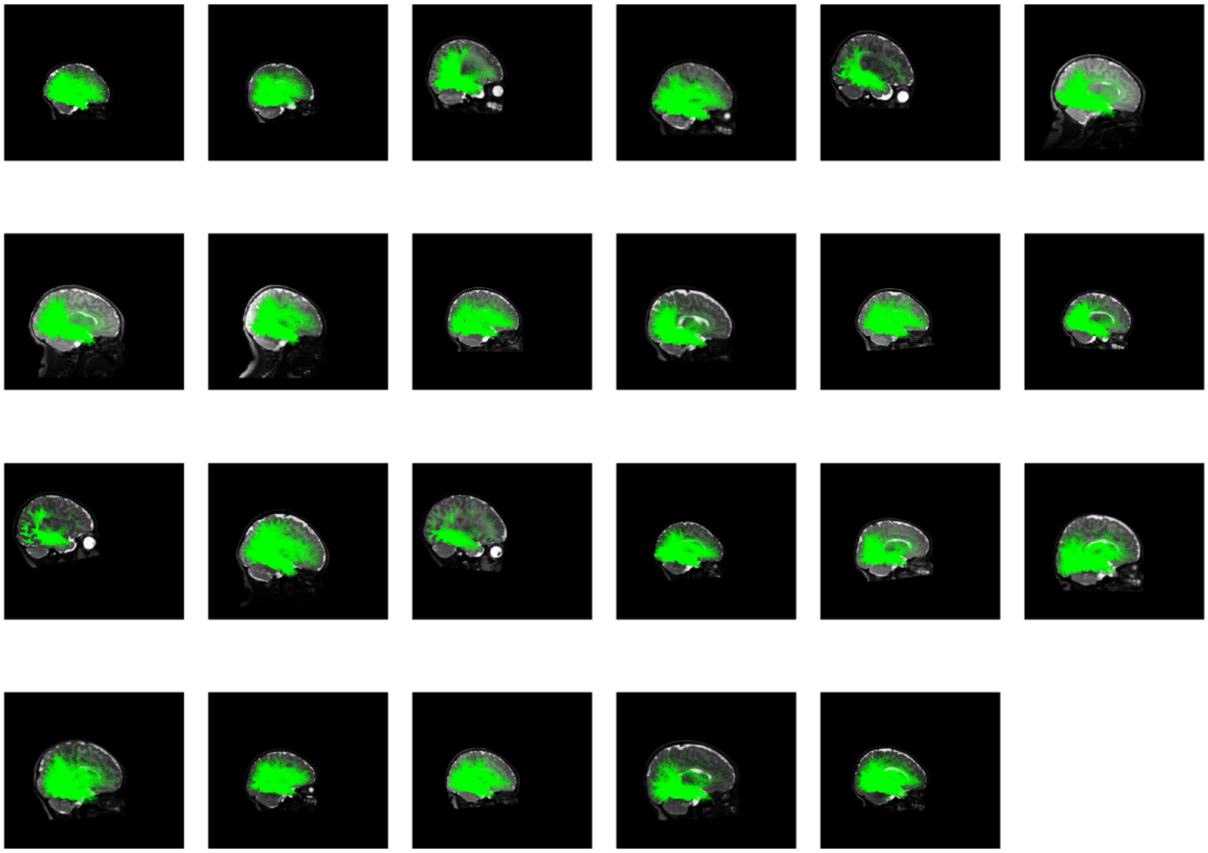

**Supplementary Figure 40.** White matter connections of CoS-places in the 0-month age group organized by age in the right hemisphere.

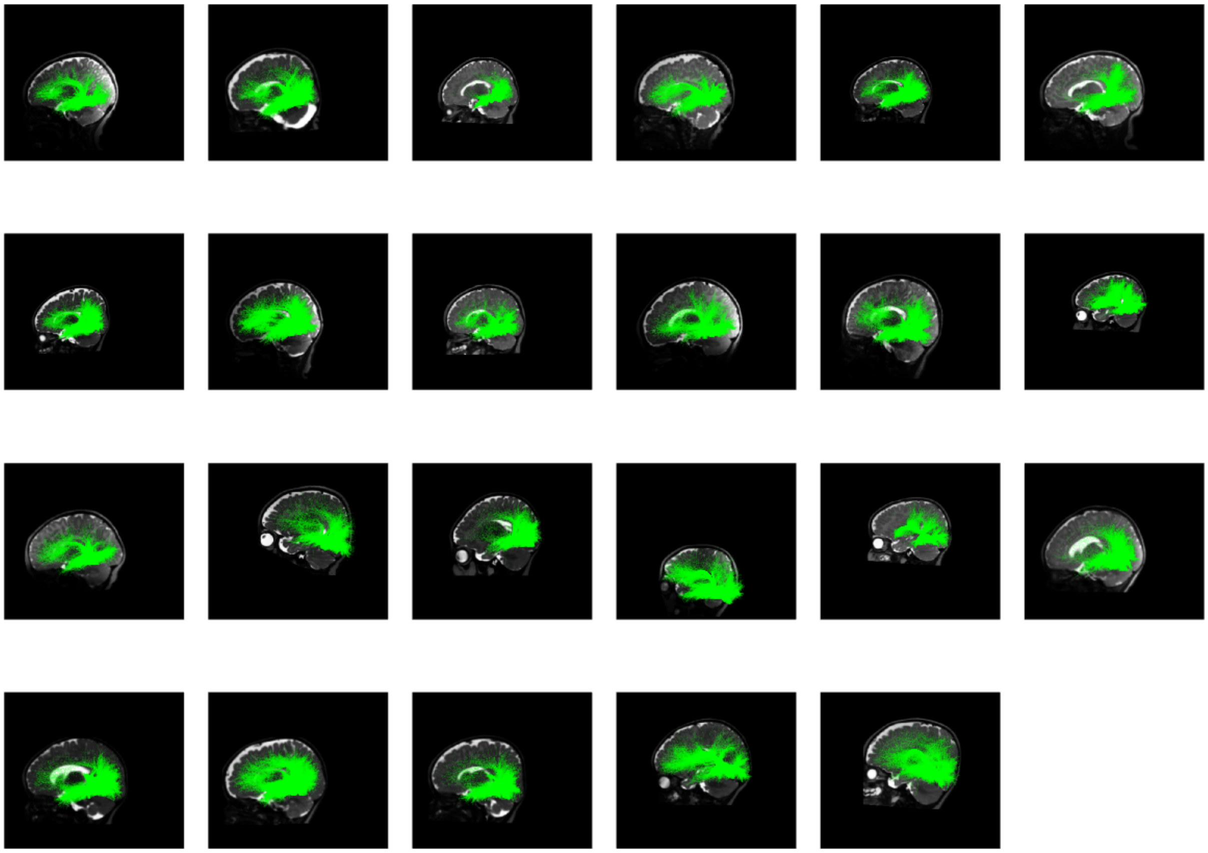

**Supplementary Figure 41.** White matter connections of CoS-places in the 3-month age group organized by age in the left hemisphere.

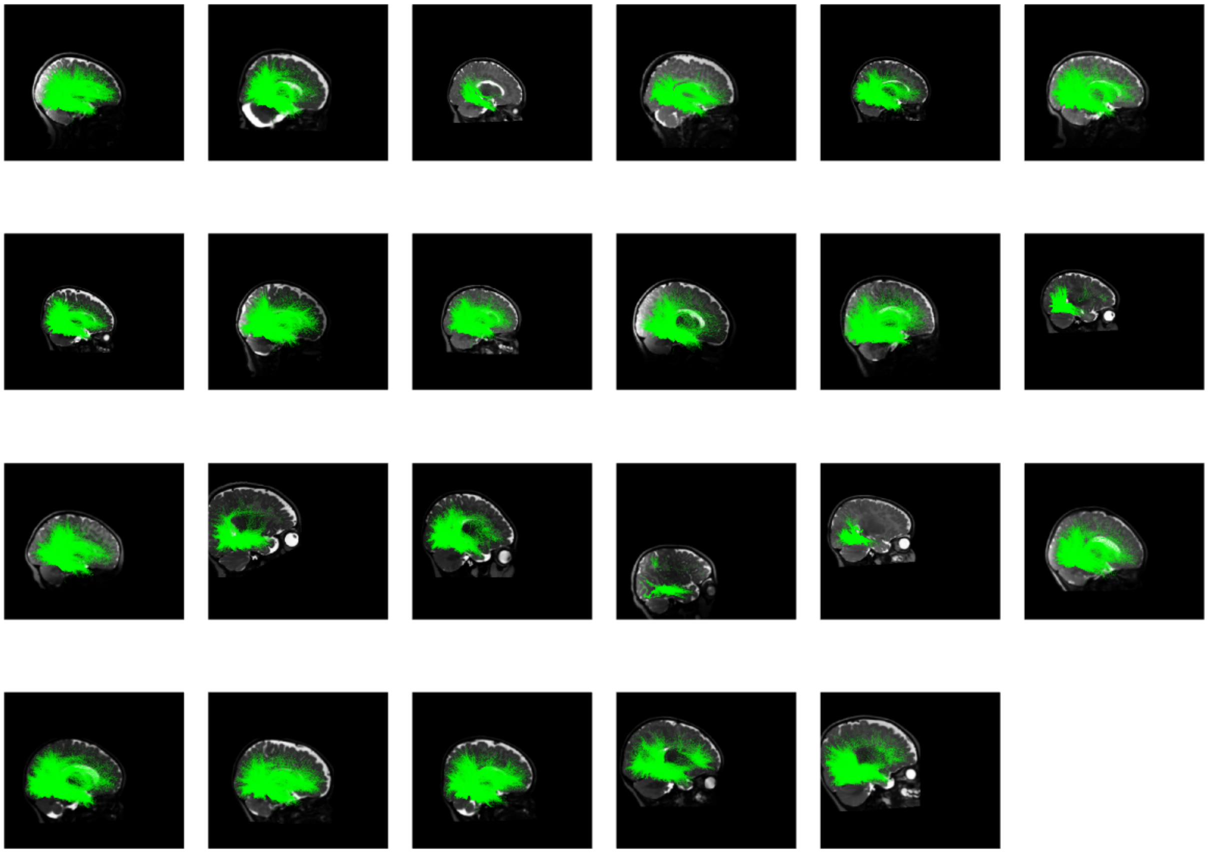

**Supplementary Figure 42.** White matter connections of CoS-places in the 3-month age group organized by age in the right hemisphere.

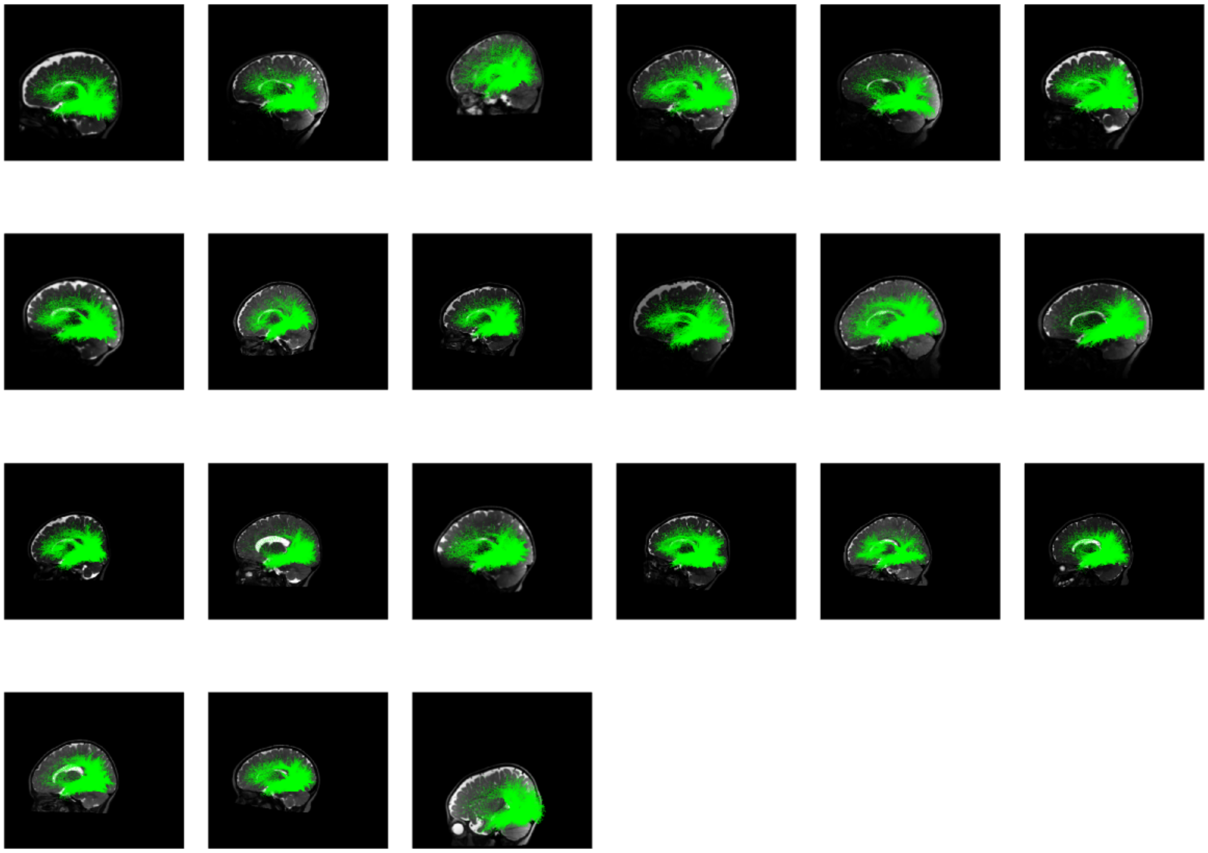

**Supplementary Figure 43.** White matter connections of CoS-places in the 6-month age group organized by age in the left hemisphere.

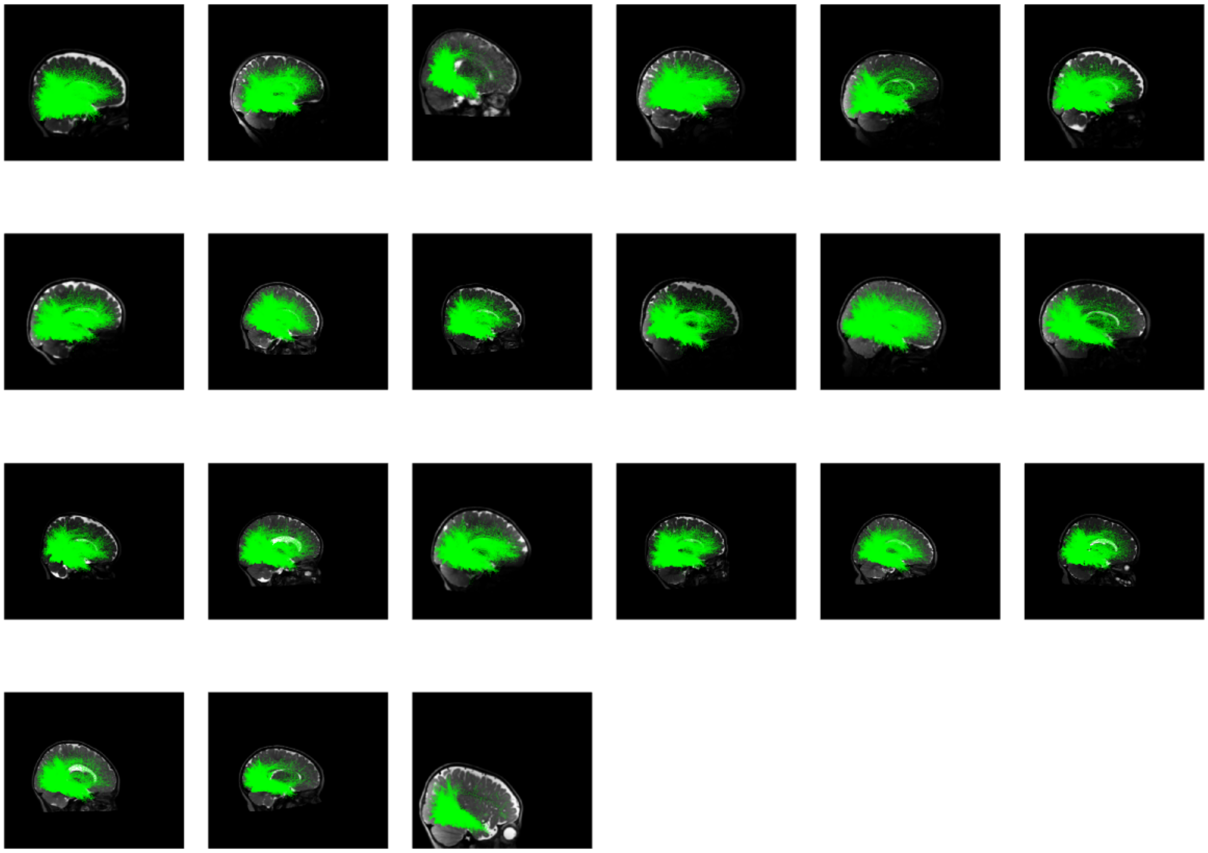

**Supplementary Figure 44.** White matter connections of CoS-places in the 6-month age group organized by age in the right hemisphere.

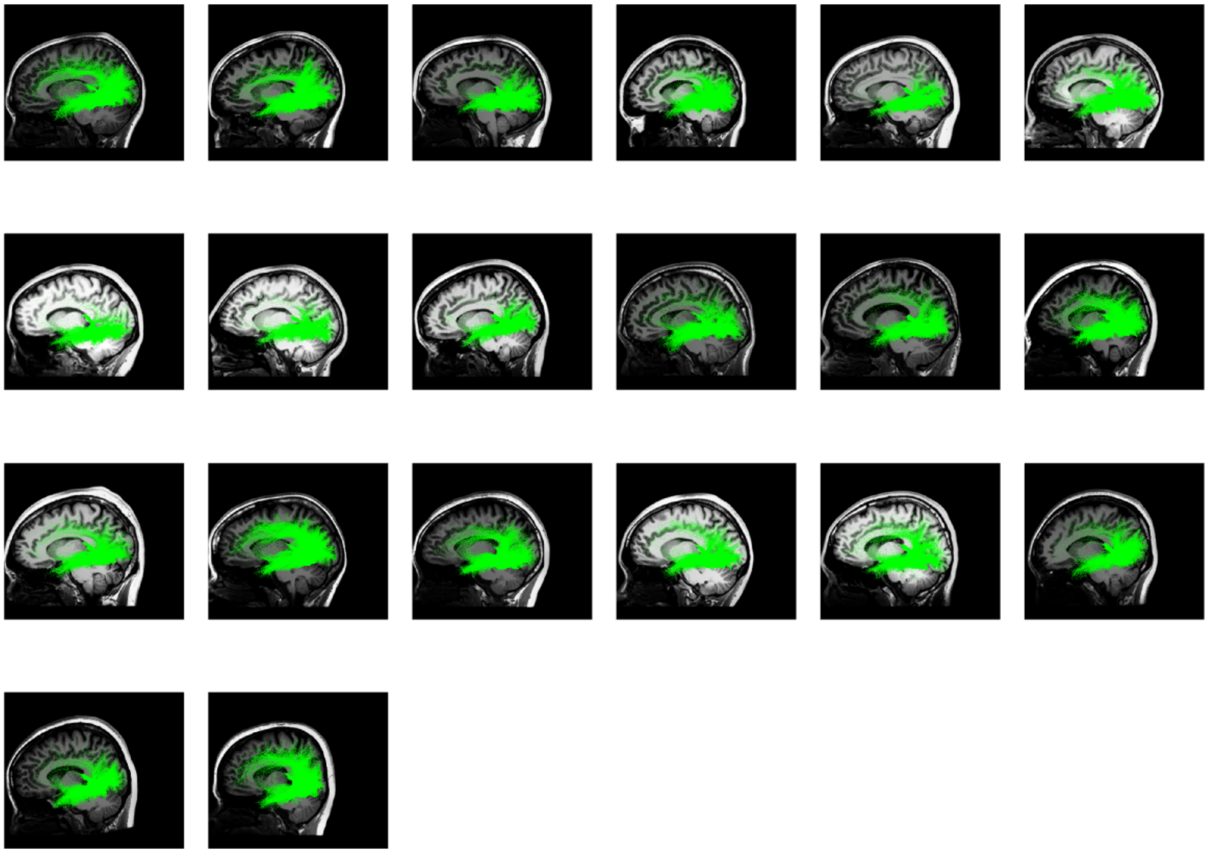

**Supplementary Figure 45.** White matter connections of CoS-places in the adult age group organized by age in the left hemisphere.

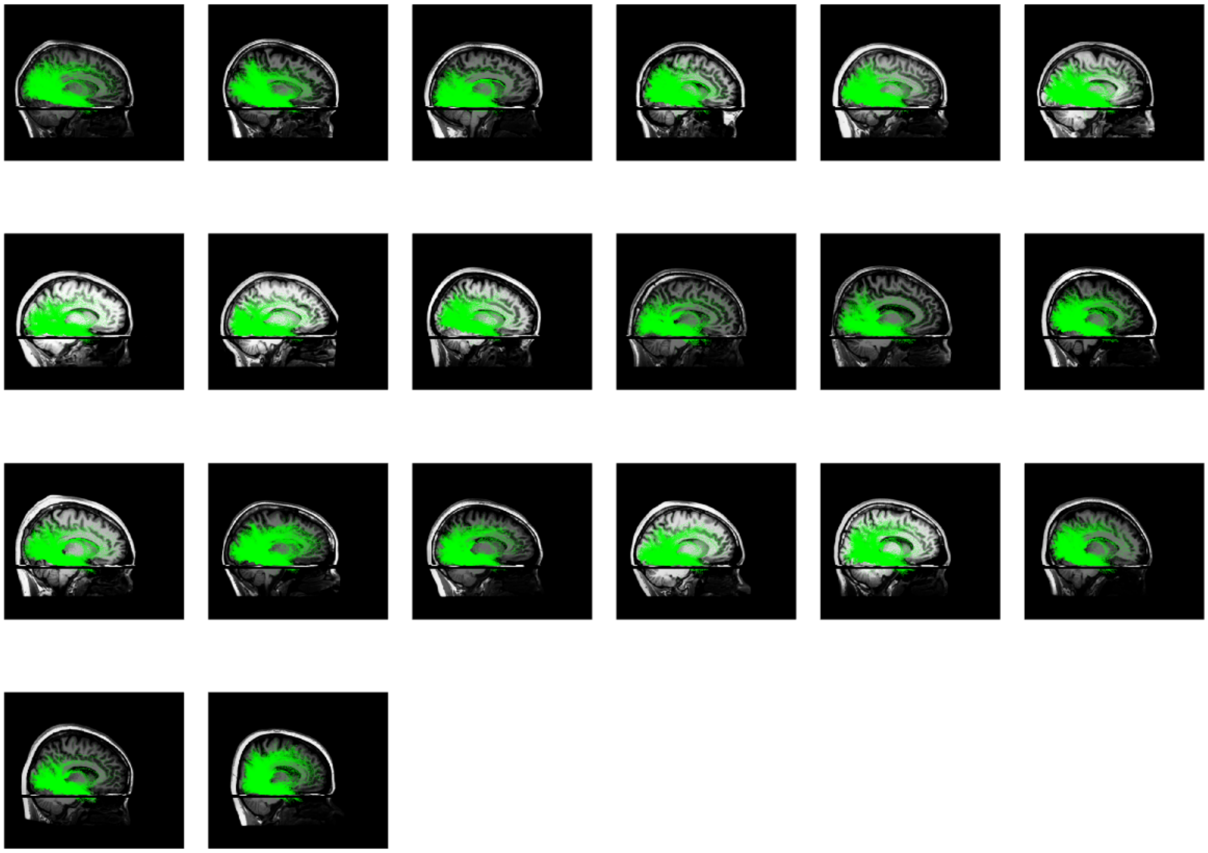

**Supplementary Figure 46.** White matter connections of CoS-places in the adult age group organized by age in the right hemisphere.

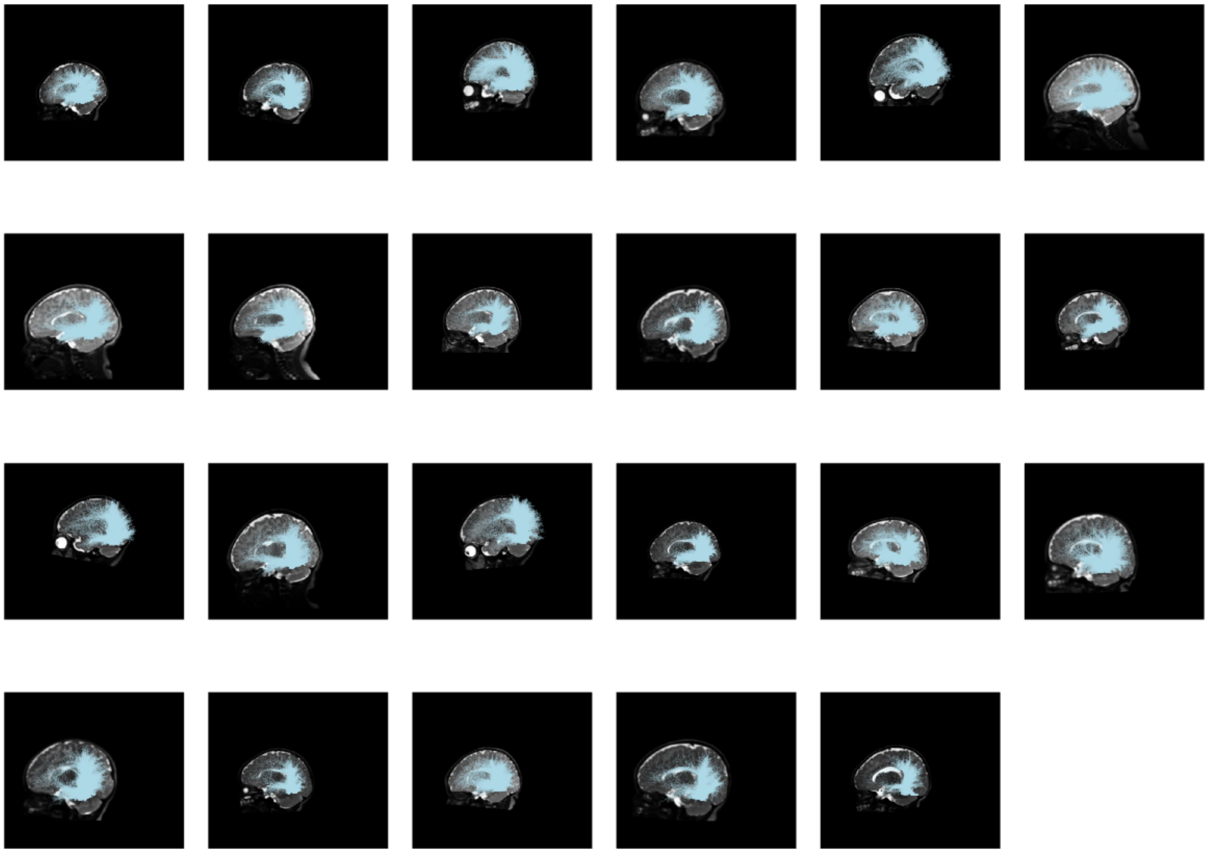

**Supplementary Figure 47.** White matter connections of mOTS-words in the 0-month age group organized by age in the left hemisphere.

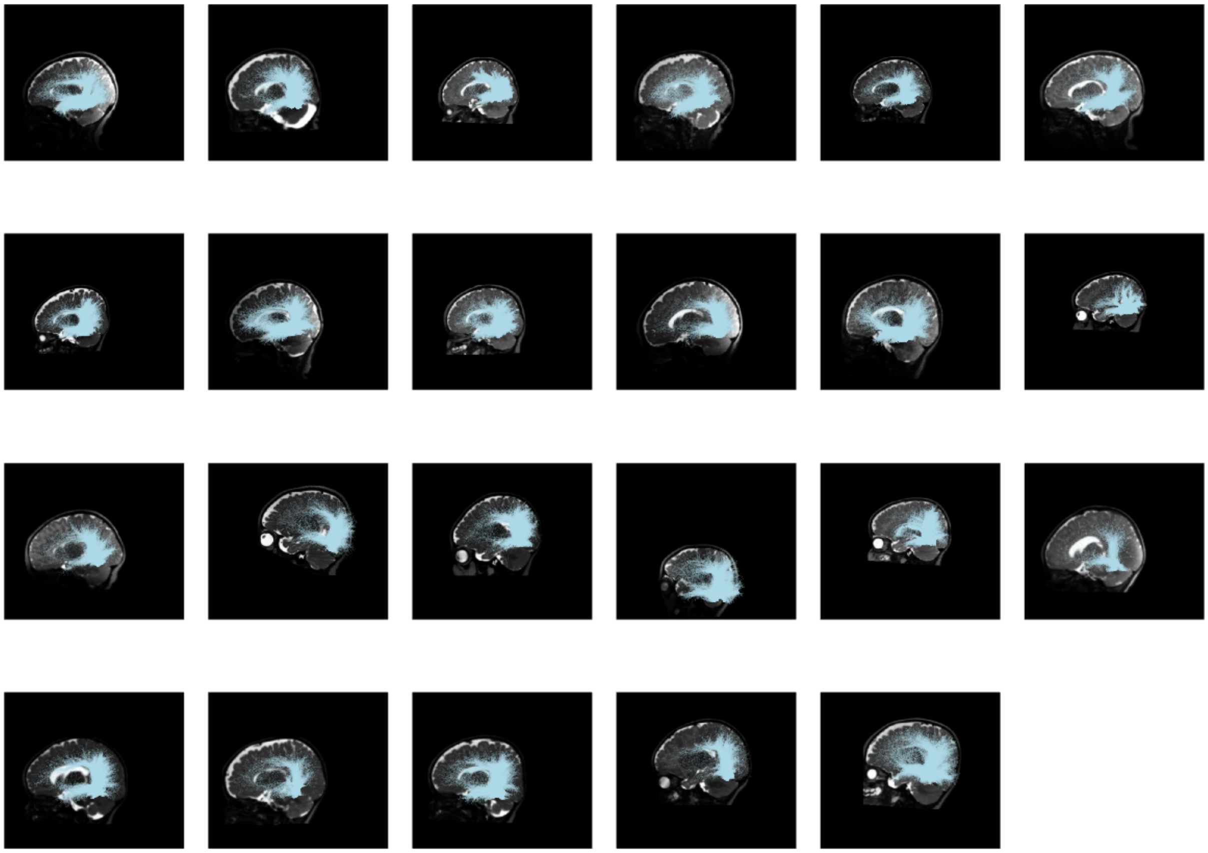

**Supplementary Figure 48.** White matter connections of mOTS-words in the 3-month age group organized by age in the left hemisphere.

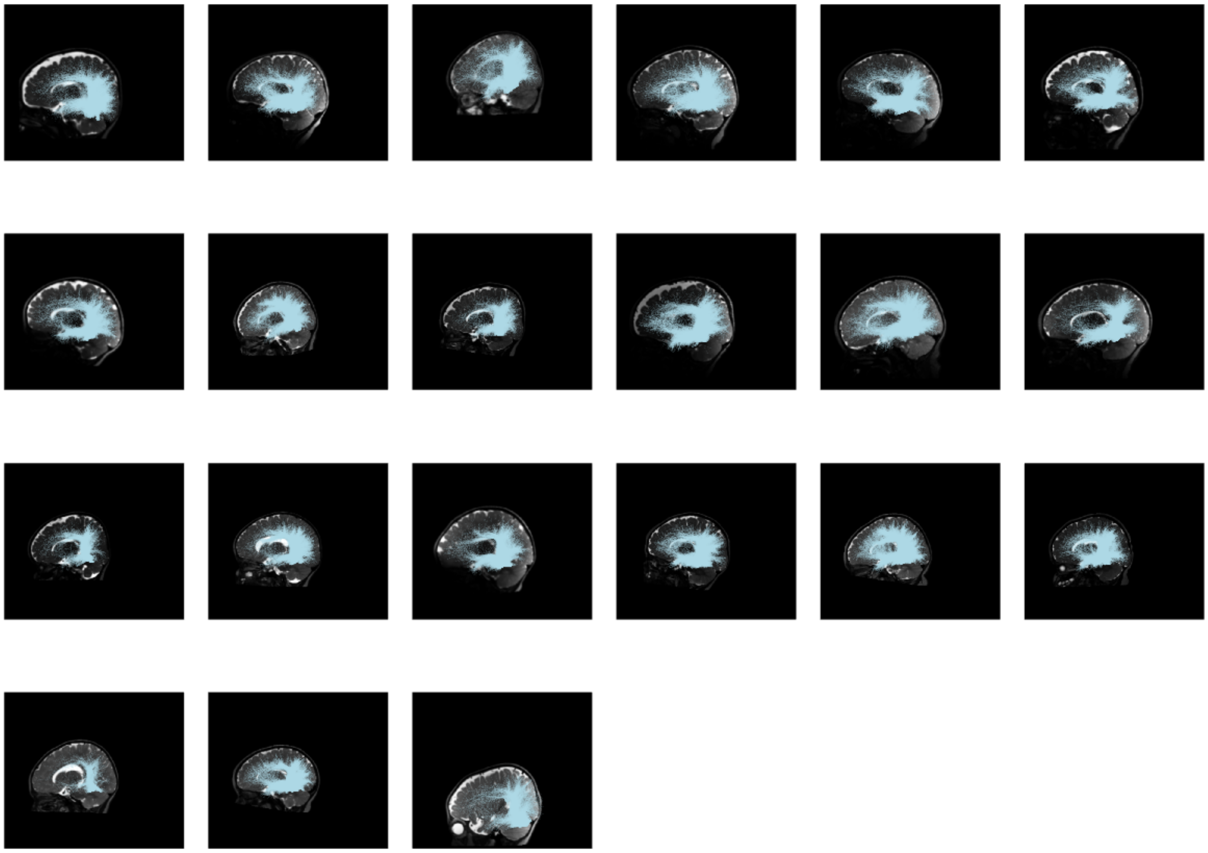

**Supplementary Figure 49.** White matter connections of mOTS-words in the 6-month age group organized by age in the left hemisphere.

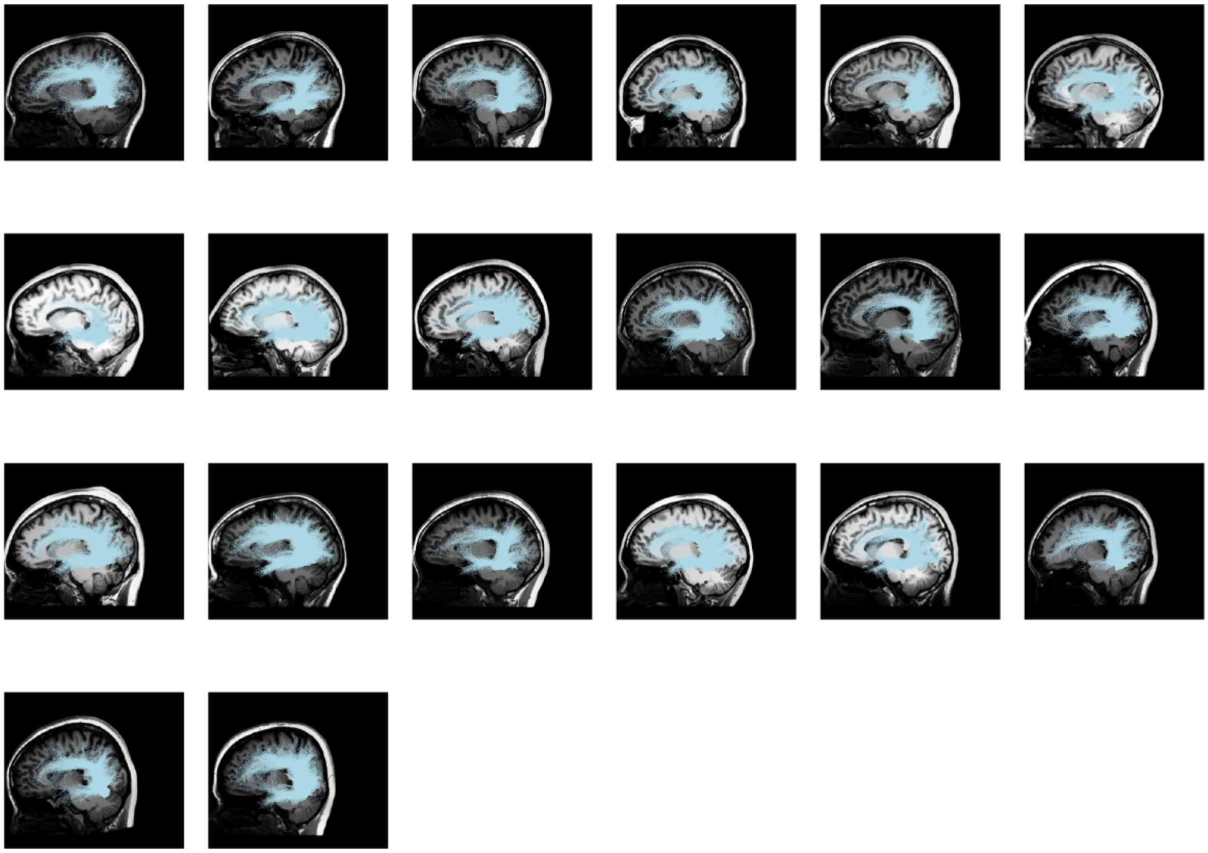

**Supplementary Figure 50.** White matter connections of mOTS-words in the adult age group organized by age in the left hemisphere.

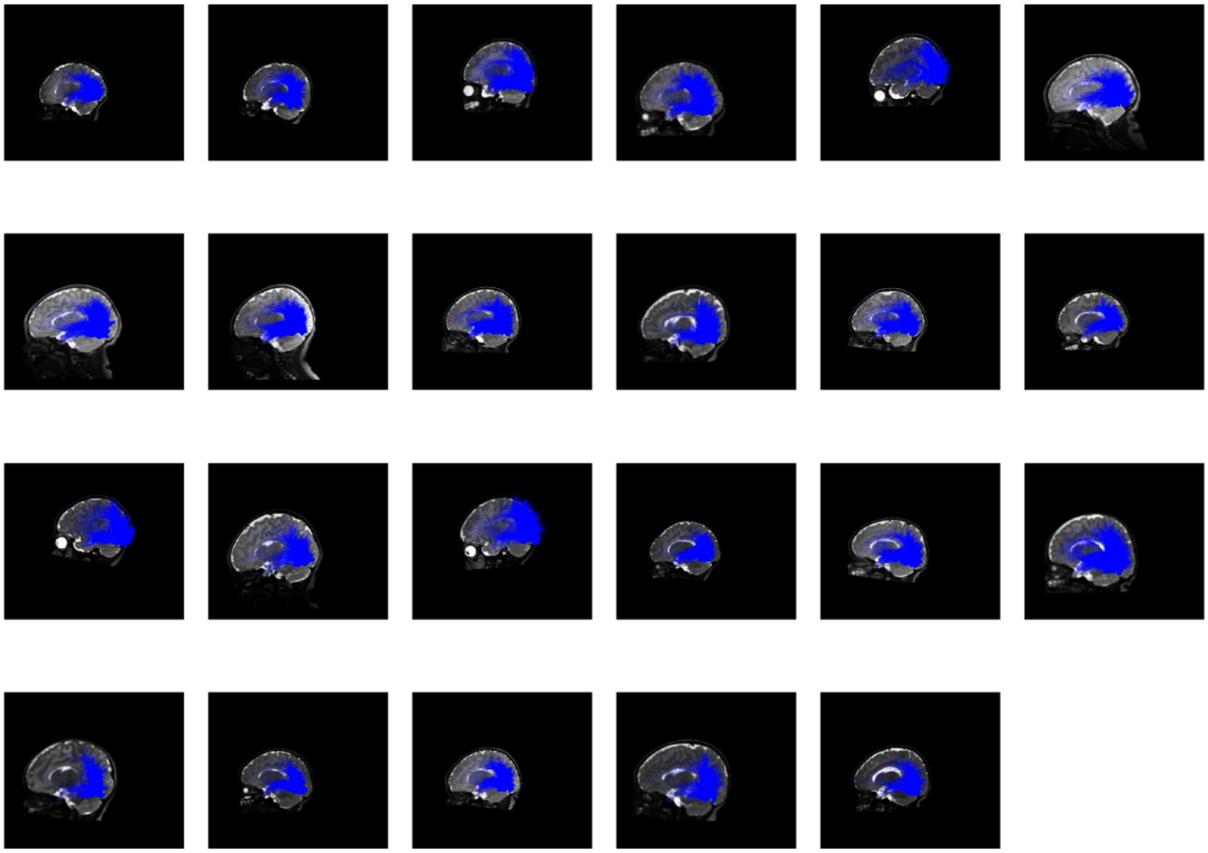

**Supplementary Figure 51.** White matter connections of pOTS-words in the 0-month age group organized by age in the left hemisphere.

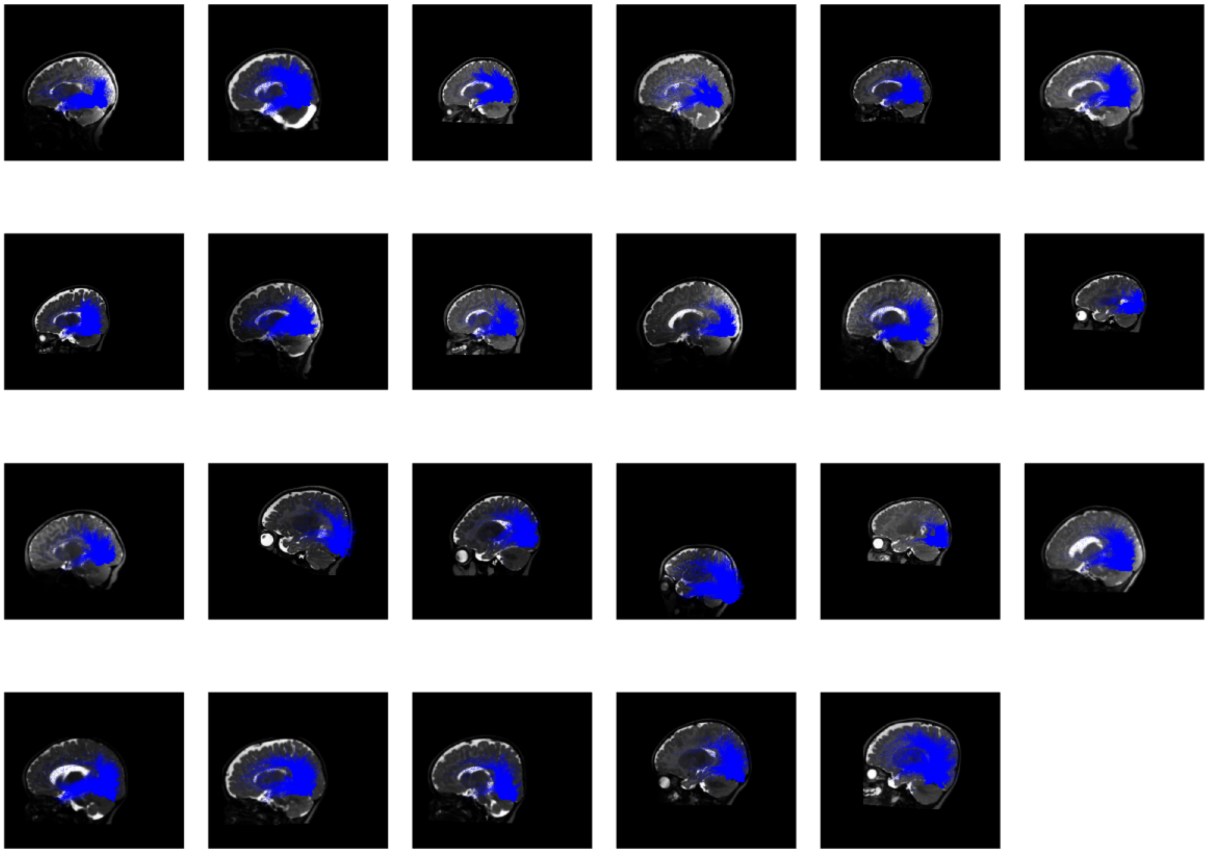

**Supplementary Figure 52.** White matter connections of pOTS-words in the 3-month age group organized by age in the left hemisphere.

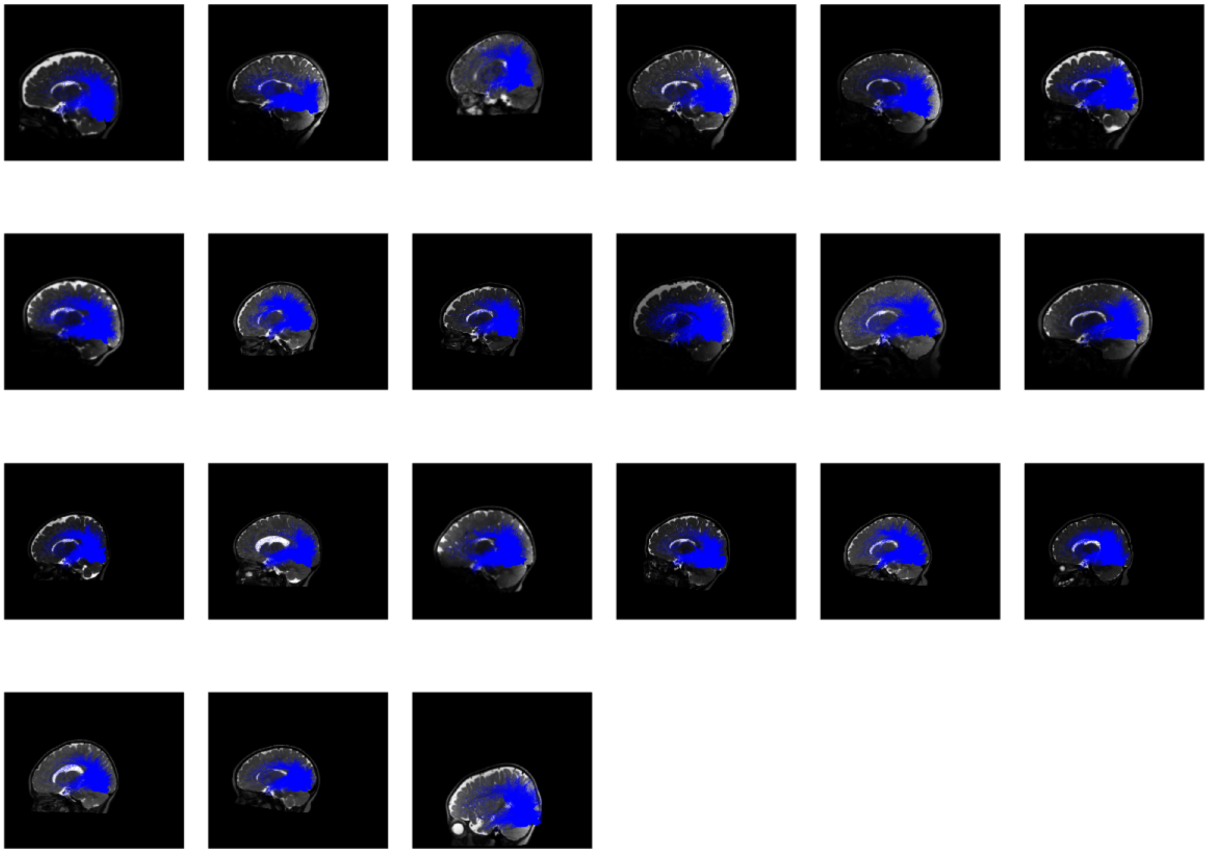

**Supplementary Figure 53.** White matter connections of pOTS-words in the 3-month age group organized by age in the left hemisphere.

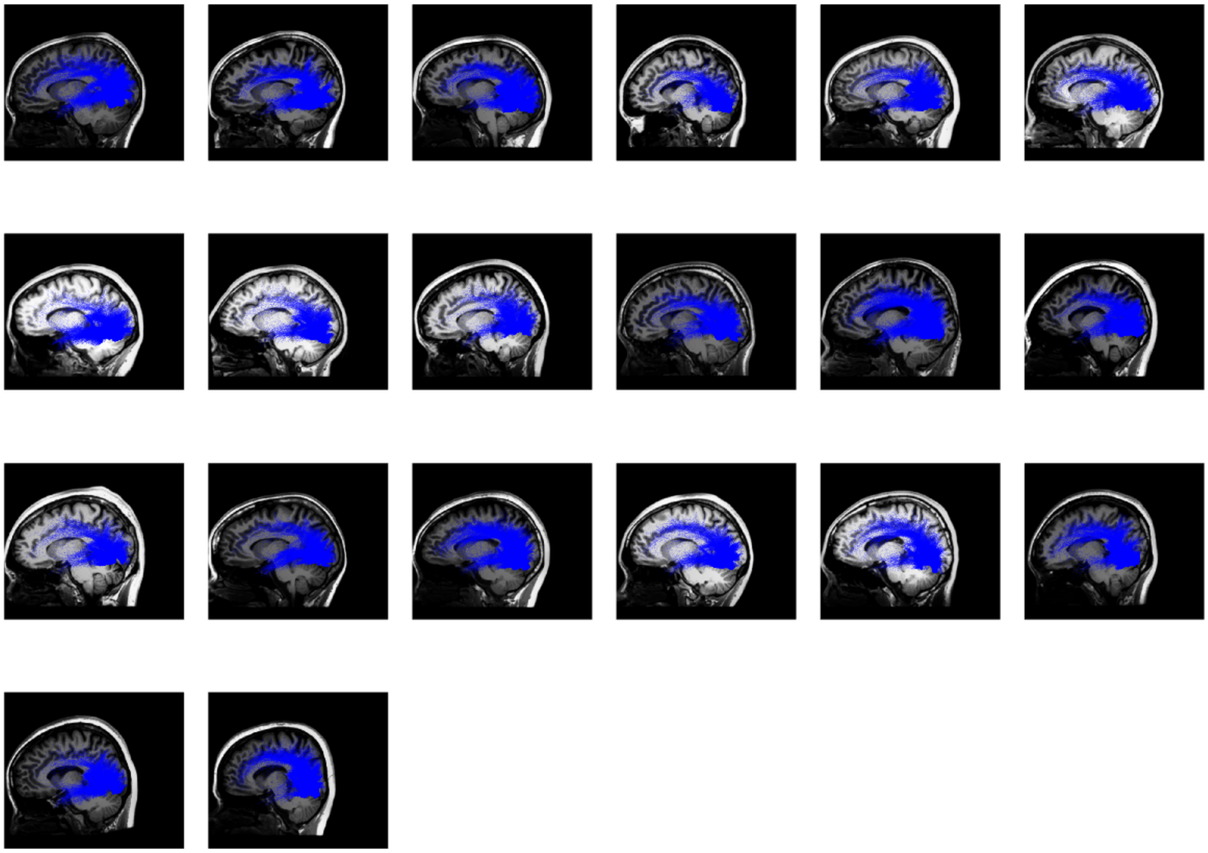

**Supplementary Figure 54.** White matter connections of pOTS-words in the adult age group organized by age in the left hemisphere.

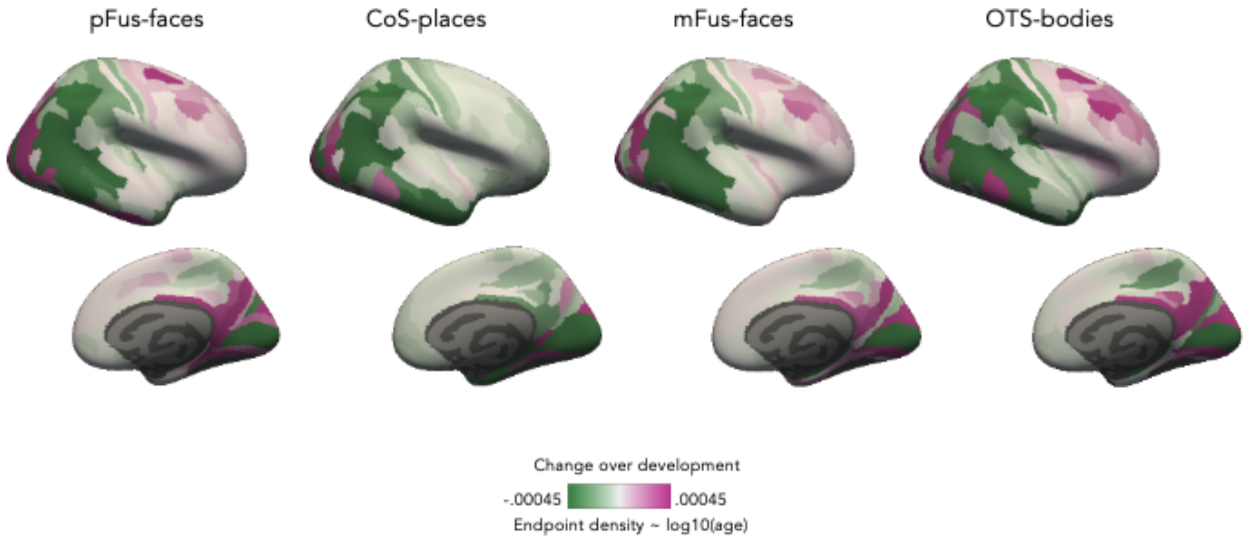


**Supplementary Figure 55.** Change in connectivity endpoint density over development within each Glasser Atlas ROI in the right hemisphere. For each fROI's connectivity profile and Glasser ROI, we fit a linear model predicting endpoint density by log10(age in days). The color of the Glasser ROI indicates the slope. *Green:* decreasing endpoint density over development; *magenta:* increasing endpoint density over development.

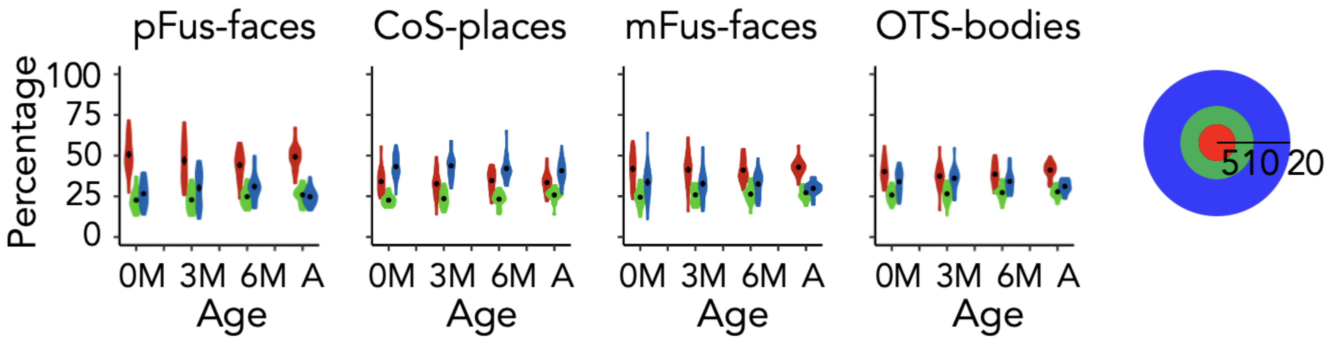


**Supplementary Figure 56.** White matter connections by early visual cortex eccentricity band in the right hemisphere. Quantification of the percentage of connections belonging to each eccentricity band across each age group. *0M:* newborns (n = 23), *3M:* 3 month-olds (n = 23), *6M:* 6 month-olds (n = 21), *A*: adults (n = 21). All panels: *red:* 0-5°, *green:* 5-10°, *blue:* 10-20°, see colorbar at right. Black dot and error bar indicate mean$\pm$ standard error of the mean.

@mFus_bb05.mov

**Supplementary Video 1.** Video of white matter connections between mFus-faces and early visual cortex in an example newborn participant. Streamlines are colored by eccentricity band. *Red:* 0-5°, *green:* 5-10°, *blue:* 10-20°.

@CoS-bb05.mov

**Supplementary Video 2.** Video of white matter connections between CoS-places and early visual cortex in an example newborn participant. Streamlines are colored by eccentricity band. *Red:* 0-5°, *green:* 5-10°, *blue:* 10-20°.

@mFus-adult.mov

**Supplementary Video 3.** Video of white matter connections between mFus-faces and early visual cortex in an example adult. Streamlines are colored by eccentricity band. *Red:* 0-5°, *green:* 5-10°, *blue:* 10-20°.

@Cos-adult.mov

**Supplementary Video 4.** Video of white matter connections between CoS-places and early visual cortex in an example adult participant. Streamlines are colored by eccentricity band. *Red:* 0-5°, *green:* 5-10°, *blue:* 10-20°.

**Supplementary Table 1.** Linear models (lm in R https://www.rdocumentation.org/packages/stats/versions/3.6.2/topics/lm) predicting endpoint density by Glasser ROI and log10(age) for each fROI. P-values are Bonferroni corrected for multiple comparisons.

(a) Main effect of Glasser ROI:
Model comparison: endpoint density ~ log10(age in days)
vs endpoint density ~ Glasser_ROI + log10(age in days)

| **fROI** | **df** | **Sum of Squares** | **F** | ***p*** |
|---|---|---|---|---|
| pFus-faces | 168 | 0.22 | 91.52 | <0.001 |
| pOTS-words | 168 | 1.91 | 454.02 | <0.001 |
| CoS-places | 168 | 1.87 | 591.41 | <0.001 |
| mFus-faces | 168 | 0.05 | 74.94 | <0.001 |
| OTS-bodies | 168 | 0.08 | 81.61 | <0.001 |
| mOTS-words | 168 | 1.04 | 210.14 | <0.001 |

(b) Glasser ROI x Age interaction:
Model comparison: endpoint density ~ Glasser_ROI + log10(age in days)
vs endpoint density ~ GlasserROI X log10(age in days)

| **fROI** | **df** | **Sum of Squares** | **F** | ***p*** |
|---|---|---|---|---|
| pFus-faces | 168 | 0.01 | 3.97 | <0.001 |
| pOTS-words | 168 | 0.03 | 6.36 | <0.001 |
| CoS-places | 168 | 0.03 | 11.86 | <0.001 |

| | | | | |
|---|---|---|---|---|
| mFus-faces | 168 | 0.002 | 3.54 | <0.001 |
| OTS-bodies | 168 | 0.005 | 4.96 | <0.001 |
| mOTS-words | 168 | 0.017 | 3.63 | <0.001 |

**Supplementary Table 2.** Linear models (lm in R https://www.rdocumentation.org/packages/stats/versions/3.6.2/topics/lm) predicting endpoint density by Glasser ROI and log10(corrected_age) for each fROI, where corrected age is the age in days minus the number of days born before 40 weeks of gestation. Two babies (four sessions) had missing gestational ages, and one baby had a negative corrected age, therefore, five sessions are excluded from these analyses. For adults, we do not have data for gestational age, and use age in days in the place of the corrected age. P-values are Bonferroni corrected for multiple comparisons.

(a) Main effect of Glasser ROI:
Model comparison: endpoint density ~ log10(corrected age in days)
vs endpoint density ~ Glasser_ROI + log10(corrected age in days)

| **fROI** | **df** | **Sum of Squares** | **F** | ***p*** |
|---|---|---|---|---|
| pFus-faces | 168 | 0.22 | 89.44 | <0.001 |
| pOTS-words | 168 | 1.88 | 445.17 | <0.001 |
| CoS-places | 168 | 1.86 | 584.80 | <0.001 |
| mFus-faces | 168 | 0.05 | 74.15 | <0.001 |
| OTS-bodies | 168 | 0.08 | 80.41 | <0.001 |
| mOTS-words | 168 | 1.04 | 210.42 | <0.001 |

(b) Glasser ROI x Age interaction:
Model comparison: endpoint density ~ Glasser_ROI + log10(corrected age in days)
vs endpoint density ~ GlasserROI X log10(corrected age in days)

| **fROI** | **df** | **Sum of Squares** | **F** | ***p**** |
|---|---|---|---|---|
| pFus-faces | 168 | 0.009 | 3.79 | <0.001 |
| pOTS-words | 168 | 0.02 | 5.66 | <0.001 |
| CoS-places | 168 | 0.03 | 11.30 | <0.001 |
| mFus-faces | 168 | 0.002 | 3.45 | <0.001 |
| OTS-bodies | 168 | 0.004 | 4.94 | <0.001 |
| mOTS-words | 168 | 0.02 | 3.90 | <0.001 |

**Supplementary Table 3.** Development of Endpoint Density for pFus-faces in the left hemisphere. Slopes and significance for linear models (lm in R https://www.rdocumentation.org/packages/stats/versions/3.6.2/topics/lm) for each Glasser ROI, relating pFus-faces endpoint density in each Glasser ROI to age. Model: endpoint density in Glasser ROI ~ log10(age in days). p-values are uncorrected for multiple comparisons.

| Glasser ROI | Slope | p-value |
|---|---|---|
| 1 | -2.09E-07 | 0.99338142 |
| 2 | -0.0002083 | 0.0574478 |
| 3a | -0.0001187 | 0.37294986 |
| 3b | -0.0002592 | 0.00230718 |
| 4 | 0.00014964 | 0.02324739 |
| 5L | -0.0001438 | 0.00097291 |

| | | |
|---|---|---|
| 5m | -2.53E-05 | 0.01229257 |
| 5mv | -0.0001707 | 0.0002866 |
| 6a | 0.00041474 | 5.41E-05 |
| 6d | 6.69E-05 | 0.01341074 |
| 6ma | 3.59E-05 | 0.06472006 |
| 6mp | -3.70E-05 | 0.35347198 |
| 6r | 5.99E-05 | 0.05449107 |
| 6v | 3.65E-05 | 0.00280483 |
| 7AL | -0.0002767 | 0.00170451 |
| 7Am | -0.000141 | 0.18281998 |
| 7m | 0.00037707 | 0.00148766 |
| 7PC | -8.81E-05 | 0.05272821 |
| 7PL | 0.00015659 | 0.05785285 |
| 7Pm | 7.01E-05 | 0.28791273 |
| 8Ad | 0.00021953 | 7.44E-08 |
| 8Av | 0.00033762 | 4.42E-12 |
| 8BL | 2.32E-05 | 0.00184995 |
| 8BM | 1.93E-05 | 0.00885228 |
| 8C | 0.00055047 | 1.31E-13 |
| 9-46d | 0.00017071 | 8.20E-08 |
| 9a | 3.20E-05 | 0.03919171 |
| 9m | 2.13E-05 | 0.04544286 |
| 9p | 2.79E-05 | 0.00264643 |
| 10d | 9.08E-06 | 0.2987735 |
| 10pp | 1.23E-05 | 0.24626557 |
| 10r | -7.55E-06 | 0.37098372 |
| 10v | 1.31E-05 | 0.04177106 |
| 11l | 7.84E-05 | 0.00200497 |
| 13l | -1.29E-05 | 0.71449167 |
| 23c | -7.35E-05 | 0.26284369 |
| 23d | -2.42E-05 | 0.16573759 |

| | | |
|---|---|---|
| 24dd | -6.04E-06 | 0.81566722 |
| 24dv | 3.75E-05 | 0.00750258 |
| 25 | -1.51E-05 | 0.01761663 |
| 31a | 5.72E-06 | 0.79393709 |
| 31pd | -5.05E-06 | 0.9548965 |
| 31pv | 0.00013413 | 0.09042476 |
| 33pr | 1.74E-06 | 0.53280805 |
| 43 | 2.28E-05 | 0.0004597 |
| 44 | 7.36E-05 | 1.13E-06 |
| 45 | 6.96E-05 | 1.69E-07 |
| 46 | 0.00017952 | 1.26E-09 |
| 47l | 1.02E-05 | 0.00144494 |
| 47m | 3.67E-06 | 0.45902841 |
| 47s | 3.01E-06 | 0.51286675 |
| 52 | -0.0003113 | 0.03136651 |
| 55b | 0.00013095 | 6.22E-06 |
| A1 | -2.32E-05 | 0.03400918 |
| A4 | -6.62E-05 | 0.00536725 |
| A5 | -6.17E-05 | 0.00453509 |
| a9-46v | 4.40E-05 | 0.05142711 |
| a10p | 4.65E-05 | 3.92E-05 |
| a24 | -6.32E-06 | 0.29810705 |
| a24pr | 8.39E-06 | 0.21983492 |
| a32pr | 1.78E-05 | 0.01266449 |
| a47r | 7.73E-05 | 8.88E-05 |
| AAIC | -4.15E-05 | 0.08836918 |
| AIP | -0.0004529 | 0.00745212 |
| AVI | 7.83E-05 | 0.00310564 |
| d23ab | 0.00013037 | 0.0935627 |
| d32 | 4.63E-06 | 0.56044889 |
| DVT | -5.92E-05 | 0.83870568 |

| | | |
|---|---|---|
| EC | -4.82E-06 | 0.68549606 |
| FEF | 0.00015764 | 1.35E-05 |
| FOP1 | 1.11E-05 | 0.00651079 |
| FOP2 | 6.71E-05 | 0.0041442 |
| FOP3 | 5.24E-05 | 0.0306128 |
| FOP4 | 0.000147 | 0.00013654 |
| FOP5 | 0.00014905 | 5.33E-08 |
| FST | 0.00027541 | 0.91724568 |
| H | 0.00034793 | 0.12740708 |
| i6-8 | 0.00016286 | 3.67E-08 |
| IFJa | 0.00024407 | 7.60E-08 |
| IFJp | 0.0001866 | 0.00030513 |
| IFSa | 0.00014258 | 3.88E-09 |
| IFSp | 0.00010232 | 5.69E-08 |
| Ig | 4.71E-05 | 0.02334003 |
| IP0 | 0.00097995 | 0.01005299 |
| IP1 | -0.0005362 | 0.05552553 |
| IP2 | -0.0008111 | 0.00052184 |
| IPS1 | -0.0002297 | 0.44863197 |
| LBelt | -0.0001296 | 0.02191268 |
| LIPd | -0.0008406 | 0.00133628 |
| LIPv | -0.0001407 | 0.09063391 |
| LO1 | 0.00148681 | 0.13240192 |
| LO2 | 0.00165584 | 0.09334997 |
| LO3 | 0.0010383 | 0.06957662 |
| MBelt | -0.0001525 | 0.00065466 |
| MI | -7.05E-05 | 0.01256127 |
| MIP | -0.00011 | 0.64899889 |
| MST | -0.0031381 | 0.01933475 |
| MT | -1.73E-05 | 0.96516077 |
| OFC | 1.37E-05 | 0.39906078 |

| | | |
|---|---|---|
| OP1 | 1.19E-05 | 0.68570702 |
| OP2-3 | 5.53E-05 | 0.21499906 |
| OP4 | 1.69E-05 | 0.1093269 |
| p9-46v | 0.00021787 | 1.60E-10 |
| p10p | 2.02E-05 | 0.03443541 |
| p24 | 2.10E-06 | 0.61407698 |
| p24pr | 2.37E-06 | 0.69589233 |
| p32 | -3.18E-06 | 0.64718462 |
| p32pr | 2.63E-05 | 0.01099533 |
| p47r | 6.22E-05 | 0.00181224 |
| PBelt | -0.0001116 | 0.00428385 |
| PCV | -0.0001933 | 0.00349511 |
| PeEc | 6.86E-05 | 0.13366908 |
| PEF | 4.50E-05 | 0.11153424 |
| PF | -0.0001723 | 0.01844862 |
| PFcm | -4.06E-05 | 0.39540053 |
| PFm | -0.0008419 | 0.01053865 |
| PFop | -4.97E-06 | 0.85196107 |
| PFt | -0.0001814 | 0.02102498 |
| PGi | -0.0017626 | 0.0011029 |
| PGp | 0.00189586 | 3.10E-05 |
| PGs | -1.02E-05 | 0.97822421 |
| PHA1 | 0.00067176 | 2.68E-15 |
| PHT | -0.0012493 | 0.00120075 |
| PI | -2.94E-05 | 0.66992671 |
| Pir | -7.28E-05 | 0.01919062 |
| pOFC | -2.33E-05 | 0.04942619 |
| PoI1 | -3.03E-05 | 0.52467075 |
| PoI2 | -9.86E-05 | 0.00066285 |
| POS1 | 0.00052004 | 1.03E-08 |
| POS2 | 0.00061776 | 0.00138497 |

| | | |
|---|---|---|
| PreS | 0.00111406 | 8.67E-14 |
| ProS | -0.000531 | 0.2354044 |
| PSL | -9.92E-05 | 0.00509584 |
| RI | -0.0001358 | 0.01150878 |
| RSC | 0.00074679 | 1.81E-10 |
| s6-8 | 4.44E-05 | 3.71E-06 |
| s32 | -4.77E-06 | 0.30383799 |
| SCEF | 3.05E-05 | 0.10164402 |
| SFL | 2.24E-05 | 0.14006858 |
| STGa | -4.23E-05 | 0.00505938 |
| STSda | -0.0002265 | 0.00028429 |
| STSdp | -0.0003712 | 0.00021977 |
| STSva | -0.000255 | 0.09759018 |
| STSvp | -0.0010191 | 0.00031424 |
| STV | -1.17E-05 | 0.22466533 |
| TA2 | -4.87E-05 | 0.05520746 |
| TE1a | -0.0001769 | 0.26337709 |
| TE1m | -0.0002722 | 0.05336723 |
| TE1p | -0.0005211 | 0.02362634 |
| TE2a | 0.00060681 | 0.02399751 |
| TGd | -9.54E-05 | 0.54061283 |
| TGv | 2.91E-05 | 0.68183832 |
| TPOJ1 | -0.0002342 | 0.0269904 |
| TPOJ2 | -0.0010167 | 0.01236571 |
| TPOJ3 | -0.0007176 | 0.11670988 |
| V1 | -0.0024928 | 0.07417306 |
| V2 | -3.36E-05 | 0.94226373 |
| V3 | -0.000241 | 0.74401188 |
| V3A | 0.00031572 | 0.33617086 |
| V3B | 0.0008191 | 0.1850266 |
| V3CD | 0.00064895 | 0.276009 |

| | | |
|---|---|---|
| V4 | -0.0084617 | 0.01415976 |
| V4t | 0.00063202 | 0.484999 |
| V6 | -1.20E-04 | 0.68656348 |
| V6A | 0.00014674 | 0.08970808 |
| V7 | 0.00014331 | 0.16184257 |
| v23ab | 0.00031909 | 1.46E-11 |
| VIP | -4.45E-05 | 0.59524904 |
| VMV1 | 0.00104255 | 5.55E-08 |

**Supplementary Table 4.** Development of Endpoint Density for OTS-words in the left hemisphere. Slopes and significance for the models relating pOTS endpoint density in each Glasser ROI to age. Model: endpoint density in Glasser ROI ~ log10(age in days).

| Glasser ROI | Slope | p-value |
|---|---|---|
| 1 | -2.65E-05 | 0.26761147 |
| 2 | -0.0003624 | 0.00166429 |
| 3a | -0.0003147 | 0.02554091 |
| 3b | -0.0003762 | 2.64E-05 |
| 4 | 3.34E-05 | 0.56820547 |
| 5L | -0.0001797 | 0.00087161 |
| 5m | -3.10E-05 | 0.00382829 |
| 5mv | -0.0002391 | 3.56E-05 |
| 6a | 0.00035784 | 0.00013971 |
| 6d | 6.38E-05 | 0.02384247 |
| 6ma | 4.35E-05 | 0.02854697 |
| 6mp | -7.46E-05 | 0.08882267 |
| 6r | 4.36E-05 | 0.04202929 |
| 6v | 4.34E-05 | 7.53E-05 |
| 7AL | -0.000356 | 0.00083796 |
| 7Am | -0.0002316 | 0.03331897 |
| 7m | 0.00036462 | 0.00201761 |

| | | |
|---|---|---|
| 7PC | -0.0001507 | 0.0028716 |
| 7PL | 0.00016528 | 0.01583931 |
| 7Pm | 4.14E-05 | 0.52473975 |
| 8Ad | 0.00020286 | 6.58E-07 |
| 8Av | 0.0002871 | 6.60E-14 |
| 8BL | 2.34E-05 | 0.00054113 |
| 8BM | 1.40E-05 | 0.04265927 |
| 8C | 0.00043331 | 1.88E-14 |
| 9-46d | 0.00013462 | 1.30E-06 |
| 9a | 2.73E-05 | 0.01975729 |
| 9m | 9.62E-06 | 0.26952225 |
| 9p | 2.86E-05 | 0.00019573 |
| 10d | 1.00E-05 | 0.13298772 |
| 10pp | 1.46E-05 | 0.02598574 |
| 10r | -4.20E-06 | 0.56494537 |
| 10v | 6.05E-06 | 0.23345281 |
| 11l | 6.57E-05 | 0.00056967 |
| 13l | -2.60E-06 | 0.89231748 |
| 23c | -0.0001155 | 0.13323476 |
| 23d | -2.55E-05 | 0.12984337 |
| 24dd | 8.71E-07 | 0.97636546 |
| 24dv | 4.83E-05 | 0.00547808 |
| 25 | -1.35E-05 | 0.02960392 |
| 31a | 3.79E-06 | 0.86212625 |
| 31pd | -7.48E-05 | 0.43274233 |
| 31pv | 0.00010308 | 0.17648061 |
| 33pr | 1.72E-06 | 0.45492514 |
| 43 | 1.38E-05 | 0.01564612 |
| 44 | 3.85E-05 | 4.09E-05 |
| 45 | 5.91E-05 | 2.11E-06 |
| 46 | 0.0001744 | 4.94E-08 |

| | | |
|---|---|---|
| 47l | 7.74E-06 | 0.00214227 |
| 47m | 6.77E-06 | 0.03311264 |
| 47s | 2.97E-06 | 0.46911409 |
| 52 | -0.0004083 | 0.00280468 |
| 55b | 0.00011373 | 2.09E-06 |
| A1 | -2.90E-05 | 0.00661352 |
| A4 | -7.06E-05 | 0.00244598 |
| A5 | -5.35E-05 | 0.020791 |
| a9-46v | 4.84E-05 | 0.00175688 |
| a10p | 3.30E-05 | 0.00075119 |
| a24 | -1.84E-06 | 0.71751707 |
| a24pr | 1.31E-05 | 0.00078292 |
| a32pr | 9.70E-06 | 0.05589053 |
| a47r | 5.36E-05 | 1.76E-05 |
| AAIC | -6.15E-05 | 0.00238964 |
| AIP | -0.0006372 | 7.97E-05 |
| AVI | 5.27E-05 | 0.00257184 |
| d23ab | 7.80E-05 | 0.28430917 |
| d32 | 4.54E-06 | 0.47295839 |
| DVT | 4.73E-05 | 0.85475735 |
| EC | -4.50E-06 | 0.58301595 |
| FEF | 1.09E-04 | 6.77E-05 |
| FOP1 | 9.01E-06 | 0.00537511 |
| FOP2 | 4.43E-05 | 0.00407796 |
| FOP3 | 5.09E-05 | 0.01254299 |
| FOP4 | 0.00011622 | 1.90E-05 |
| FOP5 | 0.00012289 | 8.80E-10 |
| FST | -0.0019634 | 0.43234508 |
| H | 0.00018939 | 0.29156638 |
| i6-8 | 0.00014761 | 3.41E-09 |
| IFJa | 0.00017054 | 4.29E-10 |

| | | |
|---|---|---|
| IFJp | 0.00014506 | 0.00230991 |
| IFSa | 0.00014263 | 2.35E-07 |
| IFSp | 8.93E-05 | 4.34E-10 |
| Ig | 1.88E-05 | 0.22219836 |
| IP0 | 0.00012852 | 0.69997448 |
| IP1 | -0.0008109 | 0.00213795 |
| IP2 | -0.0009613 | 1.24E-06 |
| IPS1 | -0.0004299 | 0.15782761 |
| LBelt | -0.0001387 | 0.00901026 |
| LIPd | -0.0010383 | 0.00011456 |
| LIPv | -0.0001791 | 0.023753 |
| LO1 | -0.0015539 | 0.38739515 |
| LO2 | 0.00386545 | 0.08763558 |
| LO3 | 0.00018701 | 0.67065123 |
| MBelt | -0.0001694 | 0.0001485 |
| MI | -6.35E-05 | 0.00324279 |
| MIP | -0.0002266 | 0.28455485 |
| MST | -0.0052575 | 0.0072882 |
| MT | -0.0012403 | 0.17383943 |
| OFC | 1.01E-05 | 0.39214674 |
| OP1 | -3.13E-05 | 0.23623648 |
| OP2-3 | -1.10E-05 | 0.71864328 |
| OP4 | 2.86E-06 | 0.6814291 |
| p9-46v | 0.00022249 | 1.70E-12 |
| p10p | 1.50E-05 | 0.05403318 |
| p24 | 3.24E-06 | 0.20842725 |
| p24pr | 6.18E-06 | 0.24312754 |
| p32 | -2.82E-06 | 0.58160945 |
| p32pr | 2.96E-05 | 0.0003801 |
| p47r | 5.43E-05 | 0.00013813 |
| PBelt | -0.0001576 | 8.13E-05 |

| | | |
|---|---|---|
| PCV | -0.0003023 | 0.00034656 |
| PeEc | -1.09E-05 | 0.74668023 |
| PEF | 1.83E-05 | 0.45566478 |
| PF | -0.0002297 | 0.00100597 |
| PFcm | -9.59E-05 | 0.01026343 |
| PFm | -0.0009737 | 0.00014435 |
| PFop | -3.36E-05 | 0.06747014 |
| PFt | -0.0002437 | 0.00071361 |
| PGi | -0.0025078 | 1.47E-06 |
| PGp | 0.00050469 | 0.16858995 |
| PGs | -0.0008165 | 0.02346779 |
| PHA1 | 0.0005485 | 3.02E-12 |
| PHT | -0.0014131 | 0.0017162 |
| PI | -8.14E-05 | 0.11967755 |
| Pir | -9.55E-05 | 0.00023405 |
| pOFC | -3.78E-05 | 0.00080481 |
| PoI1 | -4.72E-05 | 0.25255449 |
| PoI2 | -0.0001141 | 5.48E-06 |
| POS1 | 0.00060619 | 2.45E-10 |
| POS2 | 0.00044982 | 0.01262105 |
| PreS | 0.00088531 | 4.46E-15 |
| ProS | -0.0004977 | 0.22819163 |
| PSL | -0.0001564 | 8.84E-05 |
| RI | -0.0002181 | 4.74E-05 |
| RSC | 0.00079201 | 9.74E-10 |
| s6-8 | 3.78E-05 | 1.85E-06 |
| s32 | -5.88E-06 | 0.08401212 |
| SCEF | 1.77E-05 | 0.35687849 |
| SFL | 3.15E-05 | 0.02602392 |
| STGa | -3.84E-05 | 0.00506822 |
| STSda | -0.0001989 | 0.00172309 |

| | | |
|---|---|---|
| STSdp | -0.0004197 | 5.29E-06 |
| STSva | -0.000296 | 0.01442929 |
| STSvp | -0.0008698 | 0.00013761 |
| STV | -2.26E-05 | 0.03755479 |
| TA2 | -4.77E-05 | 0.00195014 |
| TE1a | -0.0002618 | 0.0639053 |
| TE1m | -0.0002246 | 0.040491 |
| TE1p | -0.0005502 | 0.09495084 |
| TE2a | 7.75E-05 | 0.67927798 |
| TGd | -0.0002593 | 0.02679301 |
| TGv | -5.61E-05 | 0.26543937 |
| TPOJ1 | -0.0003175 | 0.00490712 |
| TPOJ2 | -0.001097 | 0.00127906 |
| TPOJ3 | -0.0011158 | 0.00962074 |
| V1 | -0.002028 | 0.26222115 |
| V2 | -0.0009303 | 0.3353736 |
| V3 | -0.0042583 | 0.03963417 |
| V3A | 0.00088292 | 0.05060855 |
| V3B | 0.00045082 | 0.52042951 |
| V3CD | 0.00074277 | 0.12894642 |
| V4 | -0.014837 | 0.00024997 |
| V4t | -0.0017957 | 0.35045646 |
| V6 | 7.36E-05 | 0.8534415 |
| V6A | 0.00027238 | 0.00073752 |
| V7 | 0.00021102 | 0.04994608 |
| v23ab | 0.00032375 | 1.76E-09 |
| VIP | -9.10E-05 | 0.24159397 |
| VMV1 | 0.00101473 | 3.82E-05 |

**Supplementary Table 5.** Development of Endpoint Density for CoS-places in the left hemisphere. Slopes and significance for linear models (lm in R https://www.rdocumentation.org/packages/stats/versions/3.6.2/topics/lm) for each

Glasser ROI, relating CoS-places endpoint density in each Glasser ROI to age. Model: endpoint density in Glasser ROI ~ log10(age in days). p-values are uncorrected for multiple comparisons.

| Glasser ROI | slope | p-value |
|---|---|---|
| 1 | -2.75E-05 | 0.21519045 |
| 2 | -0.000136 | 0.01127528 |
| 3a | -0.000101 | 0.04367822 |
| 3b | -0.0001444 | 0.00996932 |
| 4 | -7.27E-05 | 0.07539221 |
| 5L | -4.22E-05 | 0.00163511 |
| 5m | -9.92E-06 | 0.01542328 |
| 5mv | -8.17E-05 | 0.00016011 |
| 6a | 6.81E-05 | 0.01911534 |
| 6d | 5.58E-06 | 0.60828319 |
| 6ma | -4.94E-06 | 0.48309606 |
| 6mp | -3.72E-05 | 0.00165086 |
| 6r | -9.22E-06 | 0.40323232 |
| 6v | 5.96E-06 | 0.33595803 |
| 7AL | -0.000121 | 0.00186461 |
| 7Am | -8.60E-05 | 0.09309486 |
| 7m | 5.70E-05 | 0.20295238 |
| 7PC | -2.99E-05 | 0.17920482 |
| 7PL | -6.24E-06 | 0.87176637 |
| 7Pm | -3.39E-05 | 0.38850457 |
| 8Ad | 1.11E-06 | 0.96126002 |
| 8Av | 6.07E-05 | 0.00550741 |
| 8BL | -1.39E-05 | 0.04759706 |
| 8BM | -2.29E-05 | 0.00160949 |
| 8C | 9.36E-05 | 0.02077319 |
| 9-46d | -0.0001833 | 0.01157035 |
| 9a | -6.48E-05 | 0.04566399 |

| | | |
|---|---|---|
| 9m | -5.92E-05 | 0.00857369 |
| 9p | -3.39E-05 | 0.02528392 |
| 10d | -7.88E-05 | 0.00064751 |
| 10pp | -0.0001095 | 0.00061652 |
| 10r | -9.16E-05 | 0.00105604 |
| 10v | -8.00E-05 | 0.00057029 |
| 11l | -0.0003008 | 4.77E-05 |
| 13l | -0.0003577 | 0.00012793 |
| 23c | -0.0001261 | 0.00118794 |
| 23d | -3.09E-05 | 0.05214842 |
| 24dd | -1.69E-05 | 0.04931556 |
| 24dv | -1.16E-05 | 0.25295472 |
| 25 | -7.54E-05 | 0.00213719 |
| 31a | -2.31E-06 | 0.79686303 |
| 31pd | -0.0001062 | 0.10238728 |
| 31pv | -0.0002121 | 0.20643315 |
| 33pr | -6.81E-06 | 0.03792324 |
| 43 | 3.99E-06 | 0.13049542 |
| 44 | -5.60E-06 | 0.39385133 |
| 45 | -1.96E-05 | 0.02155223 |
| 46 | -0.0001185 | 0.02287843 |
| 47l | -8.39E-06 | 0.12974232 |
| 47m | -3.73E-05 | 0.01417484 |
| 47s | -4.08E-05 | 0.01394026 |
| 52 | -0.0001259 | 0.39970834 |
| 55b | -4.72E-06 | 0.7420379 |
| A1 | -2.27E-05 | 0.00768765 |
| A4 | -5.28E-05 | 0.04799626 |
| A5 | -2.75E-05 | 0.23454044 |
| a9-46v | -0.0001554 | 0.00033418 |
| a10p | -0.0001801 | 0.00010936 |

| | | |
|---|---|---|
| a24 | -2.64E-05 | 0.02876316 |
| a24pr | -4.22E-06 | 0.46310687 |
| a32pr | -1.28E-05 | 0.0141394 |
| a47r | -0.00013 | 0.00014568 |
| AAIC | -1.00E-04 | 0.01000909 |
| AIP | -0.0001911 | 0.00346255 |
| AVI | -0.0001289 | 0.02118192 |
| d23ab | -5.60E-05 | 0.32746306 |
| d32 | -1.89E-05 | 0.11868347 |
| DVT | -0.0002508 | 0.30493163 |
| EC | -0.0002918 | 5.11E-05 |
| FEF | 1.04E-05 | 0.51037577 |
| FOP1 | -4.61E-07 | 0.84057363 |
| FOP2 | 2.90E-06 | 0.66380968 |
| FOP3 | -2.13E-06 | 0.80032299 |
| FOP4 | -4.45E-05 | 0.05960754 |
| FOP5 | -2.23E-05 | 0.13691087 |
| FST | -0.0004553 | 0.63029913 |
| H | -0.0058596 | 0.00177475 |
| i6-8 | 3.46E-05 | 0.00330155 |
| IFJa | 2.16E-05 | 0.11353914 |
| IFJp | 5.90E-06 | 0.82730584 |
| IFSa | -7.56E-05 | 0.01487092 |
| IFSp | -1.71E-05 | 0.17883893 |
| Ig | 5.20E-06 | 0.41456732 |
| IP0 | 0.00012932 | 0.45528767 |
| IP1 | -0.0001104 | 0.26373042 |
| IP2 | -0.0002541 | 0.00507329 |
| IPS1 | -0.0002351 | 0.03339326 |
| LBelt | -8.68E-05 | 0.00152744 |
| LIPd | -0.0002684 | 0.00825674 |

| | | |
|---|---|---|
| LIPv | -9.66E-05 | 0.03910791 |
| LO1 | -0.0003652 | 0.41071589 |
| LO2 | 0.00087045 | 0.01180142 |
| LO3 | 8.81E-05 | 0.74923298 |
| MBelt | -0.0001256 | 0.00043786 |
| MI | -0.0001018 | 0.00015153 |
| MIP | -0.0001968 | 0.04880457 |
| MST | -0.0011039 | 0.02128533 |
| MT | -0.0003428 | 0.10647977 |
| OFC | -0.0001923 | 0.00033047 |
| OP1 | -1.81E-05 | 0.2198457 |
| OP2-3 | 2.08E-06 | 0.90628994 |
| OP4 | -8.73E-06 | 0.074588 |
| p9-46v | -4.11E-05 | 0.14311801 |
| p10p | -9.97E-05 | 0.00099175 |
| p24 | -4.68E-06 | 0.18836018 |
| p24pr | -8.15E-06 | 0.12700169 |
| p32 | -5.38E-05 | 0.00295877 |
| p32pr | -1.40E-05 | 0.0983749 |
| p47r | -0.0001112 | 0.00167036 |
| PBelt | -0.0001078 | 0.01038015 |
| PCV | -8.18E-05 | 0.03948914 |
| PeEc | -0.0002929 | 0.17913937 |
| PEF | -1.14E-05 | 0.40213917 |
| PF | -8.31E-05 | 0.08362275 |
| PFcm | -4.25E-05 | 0.09543694 |
| PFm | -0.0002547 | 0.02105968 |
| PFop | -1.34E-05 | 0.10741174 |
| PFt | -7.25E-05 | 0.01878706 |
| PGi | -0.0007353 | 0.01052192 |
| PGp | 0.00040037 | 0.15251842 |

| | | |
|---|---|---|
| PGs | -1.26E-05 | 0.93481143 |
| PHA1 | 0.00249768 | 0.00820388 |
| PHT | -0.000361 | 0.27853945 |
| PI | -6.66E-05 | 0.38511008 |
| Pir | -0.0001184 | 0.01462204 |
| pOFC | -0.0002206 | 0.00034507 |
| PoI1 | -7.73E-05 | 0.27541271 |
| PoI2 | -0.0001238 | 0.00216916 |
| POS1 | -8.66E-05 | 0.81263991 |
| POS2 | -0.0001073 | 0.37518384 |
| PreS | -0.0052084 | 1.50E-05 |
| ProS | 3.50E-04 | 0.83476386 |
| PSL | -6.20E-05 | 0.00305524 |
| RI | -8.29E-05 | 0.07455268 |
| RSC | -0.0009773 | 0.00060478 |
| s6-8 | 1.95E-06 | 0.68377205 |
| s32 | -4.39E-05 | 0.00383964 |
| SCEF | -2.27E-05 | 0.00796147 |
| SFL | -2.23E-05 | 0.02068972 |
| STGa | -6.71E-05 | 0.0080951 |
| STSda | -0.0002431 | 0.0014616 |
| STSdp | -0.0003377 | 0.00591163 |
| STSva | -0.0004033 | 0.08758512 |
| STSvp | -0.000776 | 0.00168482 |
| STV | -1.74E-05 | 0.02906888 |
| TA2 | -7.16E-05 | 0.00921531 |
| TE1a | -0.000503 | 0.13514632 |
| TE1m | -0.0002613 | 0.30325695 |
| TE1p | 0.00075419 | 0.01838276 |
| TE2a | 0.00059829 | 0.2685112 |
| TGd | -0.0003173 | 0.40646868 |

| | | |
|---|---|---|
| TGv | -0.0001262 | 0.66455138 |
| TPOJ1 | -0.0002435 | 0.00027368 |
| TPOJ2 | -0.0008556 | 0.00814426 |
| TPOJ3 | -0.00042 | 0.13173245 |
| V1 | -0.0060931 | 0.08345273 |
| V2 | 0.00125124 | 0.33351673 |
| V3 | -0.0026854 | 0.05285109 |
| V3A | 0.00024869 | 0.13887691 |
| V3B | -0.0003863 | 0.04890702 |
| V3CD | -0.0002354 | 0.30825418 |
| V4 | -0.0041386 | 0.02435586 |
| V4t | 1.85E-05 | 0.95504518 |
| V6 | 3.98E-05 | 0.80307863 |
| V6A | 0.00010519 | 0.00184301 |
| V7 | 9.24E-05 | 0.01012315 |
| v23ab | 8.51E-06 | 0.89364107 |
| VIP | -4.23E-05 | 0.40289246 |
| VMV1 | 0.01662464 | 3.91E-12 |

**Supplementary Table 6.** Development of Endpoint Density for mFus-faces in the left hemisphere. Slopes and significance for linear models (lm in R https://www.rdocumentation.org/packages/stats/versions/3.6.2/topics/lm) for each Glasser ROI, relating mFus-faces endpoint density in each Glasser ROI to age. Model: endpoint density in Glasser ROI ~ log10(age in days). p-values are uncorrected for multiple comparisons.

| Glasser ROI | Slope | p-value |
|---|---|---|
| 1 | -2.51E-05 | 0.54428889 |
| 2 | -0.0002125 | 0.06088184 |
| 3a | -8.61E-05 | 0.49406147 |
| 3b | -0.0003283 | 0.00023081 |
| 4 | 9.44E-05 | 0.16647953 |
| 5L | -0.0001131 | 0.00091245 |

| | | |
|---|---|---|
| 5m | -2.15E-05 | 0.00647186 |
| 5mv | -0.0001912 | 2.08E-05 |
| 6a | 0.00029426 | 0.0004866 |
| 6d | 3.62E-05 | 0.11851896 |
| 6ma | 5.41E-06 | 0.74233159 |
| 6mp | -6.15E-05 | 0.00880254 |
| 6r | 0.00012143 | 0.00025202 |
| 6v | 8.52E-05 | 1.72E-05 |
| 7AL | -0.000217 | 0.04774239 |
| 7Am | -2.45E-05 | 0.8378981 |
| 7m | 0.00057721 | 1.86E-05 |
| 7PC | -4.57E-05 | 0.39492267 |
| 7PL | 0.0002835 | 0.00374137 |
| 7Pm | 0.00013155 | 0.1069562 |
| 8Ad | 0.00011268 | 0.00185402 |
| 8Av | 0.00043927 | 1.21E-08 |
| 8BL | 1.29E-05 | 0.16217596 |
| 8BM | -7.11E-06 | 0.50091822 |
| 8C | 0.00079673 | 2.74E-10 |
| 9-46d | 0.0001194 | 0.00216124 |
| 9a | -1.93E-06 | 0.93571014 |
| 9m | 1.49E-05 | 0.36078677 |
| 9p | 7.79E-06 | 0.53193903 |
| 10d | 1.65E-05 | 0.26888011 |
| 10pp | 6.02E-06 | 0.69634574 |
| 10r | -1.63E-05 | 0.23967191 |
| 10v | -8.59E-06 | 0.40235367 |
| 11l | 7.57E-05 | 0.04501512 |
| 13l | -4.50E-06 | 0.88914576 |
| 23c | -0.0002595 | 0.00300653 |
| 23d | -6.45E-05 | 0.02107262 |

| | | |
|---|---|---|
| 24dd | -2.89E-05 | 0.06991563 |
| 24dv | -6.73E-06 | 0.69697357 |
| 25 | -1.29E-05 | 0.20389175 |
| 31a | -2.08E-05 | 0.49416554 |
| 31pd | 0.0001568 | 0.16097452 |
| 31pv | 0.00027742 | 0.00924252 |
| 33pr | -1.41E-06 | 0.74734143 |
| 43 | 5.03E-05 | 1.57E-05 |
| 44 | 0.00011227 | 4.14E-08 |
| 45 | 8.33E-05 | 7.46E-08 |
| 46 | 0.00023351 | 1.66E-07 |
| 47l | 2.70E-05 | 0.00017676 |
| 47m | 3.35E-06 | 0.60641206 |
| 47s | 2.03E-05 | 0.06652223 |
| 52 | -0.000137 | 0.39766563 |
| 55b | 0.00014195 | 7.20E-06 |
| A1 | -1.11E-05 | 0.05296647 |
| A4 | -8.98E-05 | 0.17375687 |
| A5 | -8.36E-05 | 0.05130074 |
| a9-46v | 5.02E-05 | 0.06194484 |
| a10p | 2.12E-05 | 0.28276824 |
| a24 | -3.79E-06 | 0.68765354 |
| a24pr | -3.66E-06 | 0.46784541 |
| a32pr | 3.92E-08 | 0.99507823 |
| a47r | 9.45E-05 | 0.00038108 |
| AAIC | -5.57E-05 | 0.08932402 |
| AIP | -0.0005363 | 0.00619157 |
| AVI | 0.00014827 | 0.00022609 |
| d23ab | 0.00030292 | 0.00201282 |
| d32 | -1.09E-05 | 0.2031556 |
| DVT | -4.09E-05 | 0.81277685 |

| | | |
|---|---|---|
| EC | -7.78E-06 | 0.76173672 |
| FEF | 0.00020268 | 0.00011422 |
| FOP1 | 2.52E-05 | 0.00144453 |
| FOP2 | 0.00012255 | 1.16E-05 |
| FOP3 | 0.00012649 | 8.84E-05 |
| FOP4 | 0.00026068 | 1.55E-07 |
| FOP5 | 0.00026459 | 8.38E-07 |
| FST | 0.00178219 | 0.16290276 |
| H | -9.12E-05 | 0.82157075 |
| i6-8 | 0.00018254 | 9.49E-09 |
| IFJa | 0.00035942 | 1.74E-11 |
| IFJp | 0.00028828 | 2.54E-06 |
| IFSa | 0.00020274 | 1.36E-08 |
| IFSp | 0.00018632 | 1.76E-07 |
| Ig | 8.11E-05 | 0.00239512 |
| IP0 | 0.00047479 | 0.08313963 |
| IP1 | -8.11E-05 | 0.73797684 |
| IP2 | -0.000719 | 0.00045079 |
| IPS1 | -0.0002018 | 0.30779081 |
| LBelt | -0.0001025 | 0.00671563 |
| LIPd | -0.0006219 | 0.00450367 |
| LIPv | -4.53E-05 | 0.61897237 |
| LO1 | 0.00029145 | 0.46075332 |
| LO2 | 0.00075523 | 0.00124903 |
| LO3 | 0.0004818 | 0.28367489 |
| MBelt | -0.0001236 | 0.00104081 |
| MI | -7.48E-05 | 0.03893573 |
| MIP | 9.04E-05 | 0.69327315 |
| MST | -0.0006774 | 0.23102651 |
| MT | 0.00027631 | 0.26656011 |
| OFC | -1.96E-05 | 0.36674314 |

| | | |
|---|---|---|
| OP1 | 4.58E-05 | 0.23140088 |
| OP2-3 | 0.00012314 | 0.02642348 |
| OP4 | 2.21E-05 | 0.06186222 |
| p9-46v | 0.00030602 | 1.32E-07 |
| p10p | -1.82E-07 | 0.9913342 |
| p24 | 2.67E-06 | 0.55229697 |
| p24pr | -1.87E-05 | 0.03802315 |
| p32 | -1.81E-05 | 0.09029329 |
| p32pr | -1.38E-05 | 0.22236076 |
| p47r | 9.10E-05 | 0.00089205 |
| PBelt | -0.0001305 | 0.13274833 |
| PCV | -9.77E-05 | 0.25630982 |
| PeEc | 4.91E-05 | 0.70605738 |
| PEF | 5.63E-05 | 0.05650041 |
| PF | -0.000189 | 0.05900935 |
| PFcm | -8.34E-05 | 0.18970053 |
| PFm | -0.0007072 | 0.01811461 |
| PFop | -6.02E-06 | 0.82496099 |
| PFt | -0.0001734 | 0.03223103 |
| PGi | -0.0012346 | 0.02653926 |
| PGp | 0.00083994 | 0.05288936 |
| PGs | 0.00055468 | 0.15199313 |
| PHA1 | 0.00057088 | 5.49E-06 |
| PHT | -0.0015715 | 0.00491371 |
| PI | -0.0001305 | 0.23639014 |
| Pir | -9.60E-05 | 0.06828074 |
| pOFC | -5.18E-05 | 0.00396892 |
| PoI1 | -3.60E-05 | 0.68069214 |
| PoI2 | -9.84E-05 | 0.01085657 |
| POS1 | 0.00063512 | 3.49E-11 |
| POS2 | 0.00095007 | 8.46E-05 |

| | | |
|---|---|---|
| PreS | 0.00053586 | 0.00046359 |
| ProS | -0.0005157 | 0.0882672 |
| PSL | -0.0002115 | 0.00604394 |
| RI | -0.0003503 | 0.00553181 |
| RSC | 0.00084313 | 3.80E-10 |
| s6-8 | 2.09E-05 | 0.03173768 |
| s32 | -1.19E-05 | 0.09738145 |
| SCEF | -2.64E-05 | 0.07490733 |
| SFL | 6.17E-06 | 0.65730012 |
| STGa | -8.69E-05 | 0.00859461 |
| STSda | -0.0002696 | 0.0021254 |
| STSdp | -0.0003695 | 0.0496406 |
| STSva | -0.0005315 | 0.09665279 |
| STSvp | -0.0013162 | 0.0002906 |
| STV | -3.43E-05 | 0.05544183 |
| TA2 | -0.0001268 | 0.00159838 |
| TE1a | -0.0005883 | 0.03309857 |
| TE1m | -0.0005811 | 0.03491273 |
| TE1p | -0.0009084 | 0.10061759 |
| TE2a | 0.00044917 | 0.31684037 |
| TGd | -0.0001702 | 0.53836094 |
| TGv | 1.19E-05 | 0.90923757 |
| TPOJ1 | -0.000474 | 0.00156419 |
| TPOJ2 | -0.0016643 | 0.00086792 |
| TPOJ3 | 4.82E-05 | 0.9054877 |
| V1 | 0.00019988 | 0.77586529 |
| V2 | 0.00077056 | 0.00809789 |
| V3 | 0.00064478 | 0.06296363 |
| V3A | -5.43E-05 | 0.67989568 |
| V3B | -5.84E-05 | 0.84914685 |
| V3CD | -1.77E-05 | 0.95083746 |

| | | |
|---|---|---|
| V4 | -0.0009192 | 0.27854304 |
| V4t | 0.00025705 | 0.43723411 |
| V6 | -0.0001569 | 0.17955203 |
| V6A | 2.24E-05 | 0.68598632 |
| V7 | 1.97E-05 | 0.71477063 |
| v23ab | 0.00032347 | 3.87E-11 |
| VIP | 0.00014078 | 0.15751247 |
| VMV1 | 0.00051827 | 0.00080369 |

**Supplementary Table 7.** Development of Endpoint Density for OTS-bodies in the left hemisphere. Slopes and significance for linear models (lm in R https://www.rdocumentation.org/packages/stats/versions/3.6.2/topics/lm) for each Glasser ROI, relating OTS-bodies endpoint density in each Glasser ROI to age. Model: endpoint density ~ log10(age in days). p-values are uncorrected for multiple comparisons.

| Glasser ROI | Slope | p-value |
|---|---|---|
| 1 | -6.45E-05 | 0.13908334 |
| 2 | -0.0005186 | 0.00153575 |
| 3a | -0.0002293 | 0.11628034 |
| 3b | -0.0005556 | 1.05E-05 |
| 4 | 6.70E-05 | 0.3548793 |
| 5L | -0.0001977 | 0.0001712 |
| 5m | -4.26E-05 | 0.00018571 |
| 5mv | -0.000323 | 9.14E-08 |
| 6a | 0.00026739 | 0.00425353 |
| 6d | 2.63E-05 | 0.37072785 |
| 6ma | -8.39E-06 | 0.61035814 |
| 6mp | -0.0001112 | 0.00032089 |
| 6r | 0.00015627 | 1.26E-04 |
| 6v | 0.00010114 | 6.41E-07 |
| 7AL | -0.0003708 | 0.00724287 |
| 7Am | -0.0001631 | 0.21585673 |
| 7m | 0.00069435 | 9.55E-08 |

| | | |
|---|---|---|
| 7PC | -0.0002135 | 0.01692127 |
| 7PL | 0.00028369 | 0.00137163 |
| 7Pm | 0.00019526 | 0.0136224 |
| 8Ad | 0.00013811 | 1.91E-04 |
| 8Av | 0.00055751 | 1.04E-08 |
| 8BL | 7.16E-06 | 0.4147813 |
| 8BM | -9.86E-06 | 0.33224966 |
| 8C | 0.00095166 | 8.91E-11 |
| 9-46d | 8.79E-05 | 0.06611318 |
| 9a | 3.23E-06 | 0.89052381 |
| 9m | 1.75E-06 | 0.9183377 |
| 9p | 8.52E-06 | 0.51186993 |
| 10d | 3.92E-06 | 0.7949581 |
| 10pp | 5.02E-06 | 0.78021381 |
| 10r | -3.12E-05 | 0.03914266 |
| 10v | -8.24E-06 | 0.48543045 |
| 11l | 6.85E-05 | 0.10256702 |
| 13l | 2.78E-05 | 0.41713342 |
| 23c | -0.0004138 | 2.26E-05 |
| 23d | -8.77E-05 | 0.00855996 |
| 24dd | -7.28E-05 | 0.00073487 |
| 24dv | -3.71E-05 | 0.08942162 |
| 25 | -2.08E-05 | 0.06389866 |
| 31a | -2.67E-05 | 0.46956713 |
| 31pd | 0.00015295 | 0.16486499 |
| 31pv | 0.00028666 | 0.0122369 |
| 33pr | -3.86E-06 | 0.37615885 |
| 43 | 5.18E-05 | 1.94E-05 |
| 44 | 0.00013328 | 1.62E-11 |
| 45 | 0.00014347 | 1.28E-07 |
| 46 | 0.00024569 | 3.15E-06 |

| | | |
|---|---|---|
| 47l | 3.39E-05 | 0.00043306 |
| 47m | 4.51E-06 | 0.43978416 |
| 47s | 2.72E-05 | 0.00240921 |
| 52 | -1.09E-04 | 0.38670621 |
| 55b | 0.00018945 | 3.50E-06 |
| A1 | -2.80E-05 | 0.20730787 |
| A4 | -9.77E-05 | 0.07299988 |
| A5 | -6.51E-05 | 0.06040978 |
| a9-46v | 5.64E-05 | 0.05571752 |
| a10p | 2.15E-05 | 0.40023281 |
| a24 | -1.21E-05 | 0.30179695 |
| a24pr | 2.55E-06 | 0.66127171 |
| a32pr | 2.07E-06 | 0.72411644 |
| a47r | 0.00010961 | 0.00029989 |
| AAIC | -3.07E-05 | 0.3915287 |
| AIP | -0.0009152 | 9.52E-05 |
| AVI | 0.00024376 | 2.70E-06 |
| d23ab | 0.00030602 | 0.00185482 |
| d32 | -3.98E-06 | 0.71986763 |
| DVT | -3.25E-05 | 0.83559168 |
| EC | -4.82E-05 | 0.34195887 |
| FEF | 0.00024477 | 0.00028226 |
| FOP1 | 3.17E-05 | 9.00E-06 |
| FOP2 | 0.00018242 | 1.75E-07 |
| FOP3 | 0.00022244 | 4.05E-08 |
| FOP4 | 0.0003873 | 1.12E-10 |
| FOP5 | 0.00039092 | 4.33E-07 |
| FST | 0.00302444 | 0.10937567 |
| H | -0.0005297 | 0.36587235 |
| i6-8 | 0.00019959 | 3.68E-09 |
| IFJa | 0.00049759 | 5.96E-11 |

| | | |
|---|---|---|
| IFJp | 0.00040629 | 9.23E-07 |
| IFSa | 0.00027011 | 5.85E-08 |
| IFSp | 0.0002503 | 2.56E-09 |
| Ig | 0.00010056 | 5.67E-05 |
| IP0 | 0.0001022 | 0.6644022 |
| IP1 | -8.43E-05 | 0.73800056 |
| IP2 | -0.0009646 | 7.41E-05 |
| IPS1 | -0.0001968 | 0.23736549 |
| LBelt | -0.0001863 | 0.02954404 |
| LIPd | -0.0008614 | 0.0008802 |
| LIPv | -0.0001346 | 0.12880634 |
| LO1 | -6.81E-05 | 0.83966859 |
| LO2 | 0.00046957 | 0.01182697 |
| LO3 | 7.09E-05 | 0.84761546 |
| MBelt | -0.0001257 | 0.00017307 |
| MI | -2.37E-05 | 0.52569393 |
| MIP | 7.47E-05 | 0.7239528 |
| MST | -0.0007891 | 0.28144286 |
| MT | 0.00025061 | 0.27024273 |
| OFC | -1.53E-05 | 0.52670671 |
| OP1 | 4.87E-05 | 0.3003206 |
| OP2-3 | 0.00019484 | 0.0050414 |
| OP4 | 2.93E-05 | 0.03774062 |
| p9-46v | 0.00035519 | 8.27E-09 |
| p10p | -8.70E-06 | 0.67353565 |
| p24 | 7.06E-06 | 0.11405032 |
| p24pr | -2.51E-05 | 0.0347772 |
| p32 | -2.62E-05 | 0.06636492 |
| p32pr | -1.66E-05 | 0.18475925 |
| p47r | 0.00010613 | 0.00279726 |
| PBelt | -0.0001447 | 0.04578319 |

| | | |
|---|---|---|
| PCV | -0.000213 | 0.03642791 |
| PeEc | 9.20E-05 | 0.5726744 |
| PEF | 7.84E-05 | 0.02612348 |
| PF | -0.0002645 | 0.01664226 |
| PFcm | -0.0001068 | 0.15742922 |
| PFm | -0.0008683 | 0.01447437 |
| PFop | -4.96E-05 | 0.08827253 |
| PFt | -0.0003199 | 0.00064802 |
| PGi | -0.0012648 | 0.03975206 |
| PGp | 0.00058632 | 0.14890361 |
| PGs | 0.00062699 | 0.10622648 |
| PHA1 | 0.00063637 | 0.00014716 |
| PHT | -0.0029084 | 0.00278043 |
| PI | -0.0001181 | 0.29385347 |
| Pir | -9.08E-05 | 0.0926628 |
| pOFC | -7.07E-05 | 0.01185913 |
| PoI1 | 2.36E-05 | 0.80369685 |
| PoI2 | -7.15E-05 | 0.0508298 |
| POS1 | 0.00077283 | 6.59E-15 |
| POS2 | 0.00096983 | 5.20E-05 |
| PreS | 0.00042058 | 0.00324843 |
| ProS | -0.0004021 | 0.16782986 |
| PSL | -0.000394 | 0.0012306 |
| RI | -0.0006529 | 0.00091456 |
| RSC | 0.00096836 | 6.47E-12 |
| s6-8 | 2.32E-05 | 0.02492618 |
| s32 | -1.88E-05 | 0.02304909 |
| SCEF | -3.77E-05 | 0.02914942 |
| SFL | 1.37E-06 | 0.9211945 |
| STGa | -7.54E-05 | 0.00814143 |
| STSda | -0.0002486 | 0.00147378 |

| | | |
|---|---|---|
| STSdp | -0.0004354 | 0.01087942 |
| STSva | -0.0005172 | 0.04234007 |
| STSvp | -0.0018401 | 0.00031428 |
| STV | -7.79E-05 | 0.00275333 |
| TA2 | -9.06E-05 | 0.0040287 |
| TE1a | -0.0007029 | 0.01006699 |
| TE1m | -0.0011452 | 0.00061256 |
| TE1p | -0.0027556 | 0.00290927 |
| TE2a | 0.0002624 | 0.68622923 |
| TGd | -0.0002061 | 0.4634956 |
| TGv | 7.54E-05 | 0.44870677 |
| TPOJ1 | -0.0009931 | 0.00122355 |
| TPOJ2 | -0.0022402 | 6.84E-06 |
| TPOJ3 | -7.45E-05 | 0.86735994 |
| V1 | 0.00064 | 0.29974788 |
| V2 | 0.00097076 | 0.00075568 |
| V3 | 0.00079367 | 0.01238726 |
| V3A | 1.86E-06 | 0.98738153 |
| V3B | -0.0002508 | 0.41118423 |
| V3CD | -1.03E-04 | 0.68743816 |
| V4 | -0.000657 | 0.41135711 |
| V4t | 1.05E-05 | 0.97675397 |
| V6 | -0.0001376 | 0.1707629 |
| V6A | 3.61E-05 | 0.42231442 |
| V7 | 2.27E-05 | 0.5627882 |
| v23ab | 0.00042686 | 2.34E-15 |
| VIP | 3.13E-05 | 0.75061537 |
| VMV1 | 0.00062193 | 2.24E-07 |

**Supplementary Table 8.** Development of Endpoint Density for mOTS-words in the left hemisphere. Slopes and significance for linear models (lm in R https://www.rdocumentation.org/packages/stats/versions/3.6.2/topics/lm) for each

Glasser ROI, relating mOTS-words endpoint density in each Glasser ROI to age. Model: endpoint density in Glasser ROI ~ log10(age in days). p-values are uncorrected for multiple comparisons.

| Glasser ROI | Slope | p-value |
|---|---|---|
| 1 | -5.01E-05 | 0.31912944 |
| 2 | -0.0007516 | 0.0001441 |
| 3a | -0.0003346 | 0.06030991 |
| 3b | -0.0007335 | 7.51E-07 |
| 4 | 6.16E-05 | 0.41928078 |
| 5L | -0.0002676 | 4.57E-05 |
| 5m | -5.84E-05 | 5.04E-06 |
| 5mv | -0.0004747 | 8.84E-08 |
| 6a | 0.00029228 | 0.00803742 |
| 6d | 2.64E-05 | 0.45323604 |
| 6ma | -2.51E-05 | 0.22494564 |
| 6mp | -0.0001871 | 0.00048509 |
| 6r | 0.00023364 | 5.69E-06 |
| 6v | 0.00015609 | 6.64E-13 |
| 7AL | -0.0004914 | 0.00154699 |
| 7Am | -0.0001385 | 0.31166373 |
| 7m | 0.00090514 | 1.76E-10 |
| 7PC | -0.0002589 | 0.0053204 |
| 7PL | 0.00035506 | 0.00031738 |
| 7Pm | 0.00027577 | 0.00586826 |
| 8Ad | 0.00023046 | 1.83E-05 |
| 8Av | 0.00069378 | 7.33E-14 |
| 8BL | 1.36E-05 | 0.31579802 |
| 8BM | -7.53E-06 | 0.59344588 |
| 8C | 0.00123852 | 6.02E-18 |
| 9-46d | 0.0001484 | 0.00453179 |
| 9a | 1.23E-05 | 0.70010818 |

| | | |
|---|---|---|
| 9m | -9.94E-06 | 0.68756939 |
| 9p | 1.60E-05 | 0.35910664 |
| 10d | -2.51E-06 | 0.8951469 |
| 10pp | -5.94E-06 | 0.76027502 |
| 10r | -4.12E-05 | 0.01079641 |
| 10v | 7.54E-06 | 0.56695419 |
| 11l | 0.00013797 | 0.00014973 |
| 13l | 8.33E-05 | 0.01190187 |
| 23c | -0.0006162 | 1.15E-05 |
| 23d | -0.0001187 | 0.01239856 |
| 24dd | -0.0001104 | 0.00014255 |
| 24dv | -4.30E-05 | 0.08216882 |
| 25 | -2.42E-05 | 0.04768959 |
| 31a | -1.45E-05 | 0.7897534 |
| 31pd | 0.00018683 | 0.21124315 |
| 31pv | 0.00038137 | 0.00997513 |
| 33pr | -6.34E-06 | 0.27771936 |
| 43 | 8.54E-05 | 1.10E-08 |
| 44 | 0.0001873 | 6.77E-17 |
| 45 | 0.0001852 | 6.79E-18 |
| 46 | 0.00041603 | 3.90E-07 |
| 47l | 4.29E-05 | 1.11E-07 |
| 47m | 1.22E-05 | 0.01121792 |
| 47s | 4.42E-05 | 4.94E-06 |
| 52 | -0.0003054 | 0.04054037 |
| 55b | 0.00029051 | 4.70E-08 |
| A1 | -7.58E-05 | 0.00316852 |
| A4 | -0.0001228 | 0.00920997 |
| A5 | -7.73E-05 | 0.00741113 |
| a9-46v | 1.03E-04 | 0.00058731 |
| a10p | 5.14E-05 | 0.05607663 |

| | | |
|---|---|---|
| a24 | -1.36E-05 | 0.25652902 |
| a24pr | 2.41E-07 | 0.9707224 |
| a32pr | 6.98E-06 | 0.42963526 |
| a47r | 0.00016315 | 8.23E-10 |
| AAIC | -3.64E-05 | 0.32554489 |
| AIP | -0.0012427 | 4.21E-06 |
| AVI | 0.00031934 | 4.68E-12 |
| d23ab | 0.00045306 | 0.00074712 |
| d32 | 1.69E-06 | 0.90817059 |
| DVT | -6.09E-05 | 0.71898668 |
| EC | -3.30E-05 | 0.22277025 |
| FEF | 0.00030531 | 1.61E-06 |
| FOP1 | 4.26E-05 | 7.15E-08 |
| FOP2 | 0.00023889 | 1.66E-12 |
| FOP3 | 0.00032133 | 1.51E-13 |
| FOP4 | 0.00057743 | 2.70E-18 |
| FOP5 | 0.00044838 | 4.75E-23 |
| FST | 0.00189127 | 0.57485433 |
| H | -0.0003914 | 0.27244508 |
| i6-8 | 0.00026262 | 8.97E-10 |
| IFJa | 0.00057206 | 1.61E-21 |
| IFJp | 0.000601 | 1.97E-11 |
| IFSa | 0.00035512 | 1.56E-15 |
| IFSp | 0.00031032 | 1.88E-14 |
| Ig | 0.00014596 | 1.79E-05 |
| IP0 | -0.0002373 | 0.33727763 |
| IP1 | -0.0002231 | 0.42205831 |
| IP2 | -0.0010883 | 9.59E-05 |
| IPS1 | -0.0002761 | 0.11977922 |
| LBelt | -0.0003114 | 4.66E-05 |
| LIPd | -0.0010225 | 0.00011417 |

| | | |
|---|---|---|
| LIPv | -0.0002261 | 0.01262674 |
| LO1 | -0.000105 | 0.73532971 |
| LO2 | 0.00015886 | 0.43236792 |
| LO3 | -0.0001619 | 0.71111038 |
| MBelt | -0.0001761 | 1.65E-05 |
| MI | -2.95E-05 | 0.4795723 |
| MIP | -0.0001198 | 0.59698392 |
| MST | -0.0019053 | 0.10560026 |
| MT | -5.00E-05 | 0.80321014 |
| OFC | 4.67E-06 | 0.86084669 |
| OP1 | 9.02E-05 | 0.06950491 |
| OP2-3 | 0.00033956 | 4.54E-06 |
| OP4 | 3.78E-05 | 0.01889582 |
| p9-46v | 0.00045705 | 4.93E-13 |
| p10p | 1.05E-05 | 0.65141707 |
| p24 | 7.94E-06 | 0.09363854 |
| p24pr | -2.92E-05 | 0.02850694 |
| p32 | -3.18E-05 | 0.04306651 |
| p32pr | -2.20E-05 | 0.09234664 |
| p47r | 0.00020467 | 1.08E-09 |
| PBelt | -0.000233 | 0.00250405 |
| PCV | -0.0002873 | 0.03190017 |
| PeEc | 0.00010817 | 0.25730594 |
| PEF | 0.00015257 | 0.00127944 |
| PF | -0.0001612 | 0.2633617 |
| PFcm | -0.000124 | 0.13586392 |
| PFm | -0.0005592 | 0.16728617 |
| PFop | -5.30E-05 | 0.0741977 |
| PFt | -0.0003819 | 6.85E-05 |
| PGi | -0.0012543 | 0.05507022 |
| PGp | 0.00030764 | 0.41760601 |

| | | |
|---|---|---|
| PGs | 0.0004068 | 0.47628724 |
| PHA1 | 0.00064485 | 1.57E-08 |
| PHT | -0.0043903 | 0.18618563 |
| PI | -0.0001829 | 0.10136606 |
| Pir | -0.0001286 | 0.01235508 |
| pOFC | -5.37E-05 | 0.04817059 |
| PoI1 | -0.0001167 | 0.2773499 |
| PoI2 | -0.0001454 | 0.00361702 |
| POS1 | 0.00092764 | 7.19E-14 |
| POS2 | 0.00114263 | 3.56E-06 |
| PreS | 0.00049437 | 7.00E-05 |
| ProS | 6.31E-05 | 0.84518499 |
| PSL | -0.0004863 | 0.0006279 |
| RI | -0.0010146 | 0.00045891 |
| RSC | 0.00150717 | 1.44E-13 |
| s6-8 | 2.47E-05 | 0.05265723 |
| s32 | -2.59E-05 | 0.00215959 |
| SCEF | -5.35E-05 | 0.03498789 |
| SFL | -1.57E-05 | 0.40431034 |
| STGa | -7.71E-05 | 0.00230504 |
| STSda | -0.0003044 | 0.00018277 |
| STSdp | -0.0010247 | 0.00024895 |
| STSva | -0.0004946 | 0.03753667 |
| STSvp | -0.0037341 | 0.00702297 |
| STV | -8.37E-05 | 0.00903208 |
| TA2 | -8.09E-05 | 0.00225101 |
| TE1a | -0.0004248 | 0.02860474 |
| TE1m | -0.0008979 | 0.01428361 |
| TE1p | -0.011576 | 0.02754983 |
| TE2a | 0.00196591 | 0.04190409 |
| TGd | -0.0003496 | 0.14150582 |

| | | |
|---|---|---|
| TGv | -1.99E-05 | 0.8368654 |
| TPOJ1 | -0.0016987 | 0.00071698 |
| TPOJ2 | -0.0033462 | 0.01534868 |
| TPOJ3 | -1.78E-05 | 0.96837091 |
| V1 | -0.0001104 | 0.86591573 |
| V2 | 0.00053705 | 0.06228073 |
| V3 | 0.0003475 | 0.22630971 |
| V3A | 2.81E-05 | 0.8130781 |
| V3B | -0.0003955 | 0.12656164 |
| V3CD | -1.35E-05 | 0.94319501 |
| V4 | -0.0013047 | 0.19220565 |
| V4t | -0.0001216 | 0.70914826 |
| V6 | -8.78E-05 | 0.36303332 |
| V6A | 8.63E-05 | 0.03703133 |
| V7 | 1.37E-05 | 0.68208929 |
| v23ab | 0.0006183 | 1.23E-13 |
| VIP | 2.93E-05 | 0.76466769 |
| VMV1 | 0.00070459 | 3.53E-05 |
